# Characteristics of mathematical modeling languages that facilitate model reuse in systems biology: A software engineering perspective

**DOI:** 10.1101/2019.12.16.875260

**Authors:** Christopher Schölzel, Valeria Blesius, Gernot Ernst, Andreas Dominik

**Affiliations:** THM University of Applied Sciences, Giessen, Germany; Vestre Viken Hospital Trust, Kongsberg, Norway; University of Oslo, Norway

## Abstract

Reproducible, understandable models that can be reused and combined to true multi-scale systems are required to solve the present and future challenges of systems biology. However, many mathematical models are still built for a single purpose and reusing them in a different context can be challenging due to an inflexible monolithic structure, confusing code, missing documentation or other issues. These challenges are very similar to those faced in the engineering of large software systems. It is therefore likely that addressing model design at the software engineering level will also be beneficial in systems biology. To do this, researchers cannot just rely on using an accepted standard language. They need to be aware of the characteristics that make this language desirable and they need guidelines on how to utilize them to make their models more reproducible, understandable, reusable, and extensible. Drawing upon our experience with translating and extending a model of the human baroreflex, we therefore propose a list of desirable language characteristics and provide guidelines and examples for incorporating them in a model: In our opinion, a mathematical modeling language used in systems biology should be modular, human-readable, hybrid (i.e., support multiple formalisms), open, declarative, and support the graphical representation of models. We compare existing modeling languages with respect to these characteristics and show that there is no single best language but that trade-offs always have to be considered. We also illustrate the benefits of the individual language characteristics by translating a monolithic model of the human cardiac conduction system to a modular version using the modeling language Modelica as an example. Our experiment can be seen as emblematic for model reuse in a multi-scale setting. It illustrates how each characteristic, when applied consistently, can facilitate the reuse of the resulting model. We therefore recommend that modelers consider these criteria when choosing a programming language for any biological modeling task and hope that our work sparks a discussion about the importance of software engineering aspects in mathematical modeling languages.

## 1 Introduction

As the understanding of biological systems grows, it becomes more and more apparent that their behavior cannot be reliably predicted without the help of mathematical models. In the past, these models were confined to single phenomena, such as the Hodgkin-Huxley model of the generation of neuronal action potentials [1]. They have served their purpose up to a point where now it is necessary to take into account the upward and downward causations that link all levels of organization in a biological system from genes to proteins to cells to tissue to organs to whole organisms, populations and ecosystems [2]. These causations span effects on multiple scales of space and time, which need to be included in models. This can be achieved by two different approaches. A *micro-level* model combines thousands of individual homogeneous submodels to reach the next higher scale. This approach requires a vast amount of computing power and is therefore usually limited to span a distance of only two scales. More wide-spanning multi-scale models can be achieved by the *multilevel* approach, which combines both macro- and micro-level descriptions of a system by different models [3]. While micro-level parts of such a model may look as described above, the macro-level parts feature heterogeneous descriptions of subsystems and their high-level interactions. For this approach, a wide variety of techniques exist that reduce the computational complexity of resulting models [4]. While both approaches require the reuse of existing models, the multi-level approach additionally involves the combination of independent submodels, which may have been designed for different purposes and in different labs. These submodels may even use different modeling formalisms, thus forming a *multi-class* model [5].

The first step in building a model consisting of several submodels is to regenerate the individual parts from the literature. This can already be a challenge due to several issues with reproducibility in systems biology including incomplete model descriptions, errors in formulas, availability of the code or missing descriptions of experiment setup or design choices [6–8]. A recent study by curators of the BioModels database showed that only 51% of 455 published models were directly reproducible [8]. In an extreme case, Topalidou *et al.* [9] report requiring three months to reproduce a neuroscientific model of the basal ganglia.

We experienced similar reproducibility issues first-hand when we translated the Seidel-Herzel model (SHM)—a macro-level model of the human baroreflex, which is able to simulate many disease conditions and exhibits interesting dynamical properties—into a form that would be more amenable to extension and reuse [10–12]. Even though we could reach out to the author of the model to obtain his original implementation in C, the translation process was still quite challenging. The C code was monolithic and imperative in nature, describing calculation steps instead of mathematical relations and containing details that where not described in the corresponding PhD thesis. We had to carefully extract the meaning of each line of code in order to build a modular, declarative version that produced the same simulation results. For this task, we chose the modeling language Modelica, since it provides a lot of flexibility for modular model design. However, when we wanted to extend the model with a trigger for premature ventricular contractions (PVCs), it turned out that even our new version was not suitable for reuse. In fact, the component that described the cardiac conduction system remained monolithic and lacked a graphical representation, which made it hard to identify the equations and variables that would have to be changed. This situation—having to untangle the semantics and code of an existing model to extend or adjust it for use in a different context—is emblematic of the challenges faced when building multi-scale and especially multi-level models. Our example shows that issues with reproducibility and reuse reach down to the engineering level. The modeling language and the design principles applied to the construction of a model can facilitate or hamper further use. This also holds for the aforementioned case of Topalidou *et al.* [9], since the original model was implemented in Delphi, which is also an imperative language and therefore not well suited for mathematical modeling.

Even though the need for design principles on the engineering level is apparent, most publications about best practices for reproducibility and reusability do not address it. Instead, existing approaches broadly fall into three (overlapping) categories. They tend to focus on a) biological validity [13–18], b) high-level choices of modeling formalisms and techniques [19–21], or c) model documentation, annotation and distribution [7, 22, 23]. Apart from these general discussions about reusability, there also are authors who advocate for individual modeling techniques, such as modular model design [24] in CellML or coupling models via semantic annotations [25]. However, while the latter work is applicable to multiple languages, it only focuses on one, albeit central, part of model design, namely the composition of multiple models or model parts. This means that researchers who want to select a suitable language for their modeling task still have little guidance available. The best assistance for choosing a modeling language currently comes from the list of accepted standards published by the COmputational Modeling in BIology NEtwork (COMBINE) [22, 26]. The COMBINE suggests to use CellML and the Systems Biology Markup Language (SBML) with the main reason that these are standard exchange formats that have a high interoperability among several tools. While this is true and a great improvement over the previous state of the art, standardization and interoperability alone cannot guarantee reusable model design. For example, the BioModels database [27] features curated models in SBML format, but most of these models are monolithic and therefore require further modification if only parts of the model should be reused [28]. In the aforementioned reproducibility study, the curators of this database found that the reproducibility rate for SBML models was only slightly higher (56%) than the overall rate of reproducibility across all models (51%; including models written in SBML, MATLAB, Python, C, R, and other languages) [8]. Our previous example of the translation of the SHM also shows that using a suitable language is a necessary but not sufficient criterion for the model to actually be reusable. Additionally, no single language or even a small set of prescribed languages is likely to cover all use cases which may arise in systems biology, especially when considering multi-class models, which combine entirely different model formalisms [7].

Even when the discussion is restricted to the formalisms of ordinary differential equations (ODEs) and discrete events, there are a multitude of languages to choose from. As mentioned above, the COMBINE lists SBML and CellML as accepted standard languages. Both are markup languages based on the eXtensible Markup Language (XML) and designed to be written and read by software tools and not directly by humans. While SBML has a clear focus on metabolism and cell signaling models, CellML, despite its name, is not targeted towards a specific level of organization. MATLAB is a proprietary domain-specific programming language designed for scientific computing in general, which is also popular in systems biology (https://www.mathworks.com/products/matlab.html). It provides an environment for graphical block diagrams called Simulink (https://www.mathworks.com/products/simulink.html) and a declarative language for designing physical systems called Simscape (https://www.mathworks.com/products/simscape.html). The MATLAB environment SimBiology is another alternative based on block diagrams, which is targeted towards pharmacological models, but can, like SBML, model arbitrary ODE-based dynamical systems (https://de.mathworks.com/products/simbiology.html). While MATLAB is still popular [29, 30], the open-source programming language Python also gains increasing interest in the community [29, 31–34]. Usually models are not built in Python itself, but researchers have created packages such as PySB [33] and the Python Simulator for Cellular systems (PySCeS) [31] that define embedded domain-specific languages (DSLs) which facilitate the creation of mathematical models for specific use cases. With Tellurium [34] there also exists a broader python-based environment that supports multiple COMBINE standards and uses the declarative modeling language Antimony [35]. Another emerging language for the definition of embedded DSLs for mathematical models is Julia, which has a similar focus as Python but is more extensible and tends to have better runtime performance [36]. Finally, Modelica is an open-source declarative modeling language primarily used in engineering [37]. It has a large user base both in industry and research, but is still largely unknown in systems biology. Notable exception include the Physiolibrary [38]—a Modelica library for physiological models—and SBML2Modelica [39]—a tool that translates SBML models to Modelica. This extensive list of language candidates makes it apparent that researchers need guidelines to choose between these candidates and to write model code that actually uses the desirable characteristics of the chosen language.

In recent years there has been increasing interest to apply techniques from software engineering (such as unit testing, version control, or object-oriented programming) to modeling in systems biology [28, 30, 40]. Hellerstein *et al.* [40] even go so far to suggest that systems biologists should rethink the whole modeling process as “model engineering”. To date they are the only authors that we are aware of who actually give explicit guidelines for how to write model code (e.g., they suggest to use human-readable variable names).

In this article we share our experience with extending the SHM and generalize our findings from this example to expand on the idea of model engineering in three ways: First, we propose a list of desirable characteristics that make a model language suitable for building reusable multi-scale models. Second, we give guidelines on how these characteristics can be exploited during model design to increase the reproducibility and reusability of a particular model. Third, we compare state-of-the-art language candidates with respect to the aforementioned characteristics.

From these candidates we chose one, namely Modelica, to demonstrate the reasoning behind the characteristics, guidelines, and language assessment using the example of the cardiac conduction system within the SHM. We transform the existing monolithic model into a modular structure and show how this facilitates the PVC extension. After we present our results we reflect on the impact that each of the language characteristics and the choice of Modelica in particular has on the usefulness of the model.

## 2 Results

### 2.1 Desirable characteristics for a mathematical modeling language for systems biology

The following characteristics were developed from literature review and/or from our personal experience with Modelica and the SHM. The goal of these characteristics is to facilitate the creation and analysis of multi-scale, multi-level and multi-class models. We therefore focus on increasing reproducibility, understandability, reusability, and extensibility. The resulting characteristics are that a modeling language should be modular, human-readable, hybrid, open, declarative and graphical. Each characteristic will be introduced with arguments for its usefulness, a brief set of guidelines on how it may be applied to full effect, examples where this was relevant in our implementation of the SHM, and references to other authors that advocate this feature.

#### Modular

In order to replace or reuse parts of a model, they have to be identified in the code. The modeling language should make this as easy as possible, using separable components with clear interfaces. The number of variables in the interface should be minimal, encapsulating internal implementation details so that using and connecting the component becomes as easy as possible. Some languages facilitate this by allowing the definition of connector components, which group interface variables together, so that the interface has, e.g., a single electrical pin connector instead of two separate variables for current and voltage. Interfaces are important to define intended biological transitions between model components and to document assumptions, even if rigid interfaces can limit reuse. It can even be argued that it is beneficial if a component cannot easily be reused in an environment with different assumptions, since such a switch of assumptions will likely require more change than adding a variable to the interface. For quick experimentation, it can be an advantage if the language allows connecting arbitrary internal variables of components, but published versions of a model should always have a clear interface concept to remain understandable.

Modularization and encapsulation are reliable tools to handle complexity in large software projects, so it is reasonable to expect that they will also be able to manage the complexity of biological systems. Modularity also inherently facilitates reusability, since clearly defined self-contained modules are easier to reuse than a set of equations that has to be extracted from a tightly coupled model. To allow reuse of components within the same model, it must be possible to import multiple instances of a module and assign individual identifiers to them. This can be further facilitated by supporting full object-orientation, allowing a component to inherit variables, equations and possibly annotations from one or more parent components, which define common structure and behavior. Additionally, components are also easier to reuse if individual variables and equations may be overwritten or removed during instantiation and inheritance. Some languages also allow the reuse of models across different languages, tools and platforms by using a standardized exchange format or a standardized interface. In systems biology, SBML is a standard exchange format for hundreds of tools, allowing the use of models in a multitude of different contexts often through the use of translators that convert SBML to different languages. In contrast, the Functional Mock-up Interface (FMI) is a model exchange format maintained by the Modelica Association [41, 42] that focuses more on a unifying interface than a unified language. It is not used to translate models into other languages, but rather to distribute models in an encapsulated format that is independent of the underlying formalism, which is especially interesting for multi-class models.

##### Guidelines

Modules should be small enough to be understandable at first glance, but still self-contained. If a formula or concept is used multiple times in a model, it should be defined as a module once and then referenced. In software engineering this concept is called DRY for “don’t repeat yourself”. Modules should have clearly defined, minimal interfaces, which explicitly state possible connection points to the outside world. Both modules and their interfaces should follow the biological structure of the system. If a module represents more than one biological entity or an equation in the module conflates effects from multiple distinct causes, it might be worth to investigate whether splitting up the corresponding module further might increase its understandability and flexibility for reuse and extension. Interfaces should represent the transfer of some physical quantity between biological entities and should only expose variables whose meanings are clear and do not require an understanding of the module’s internal organization or function. If possible, each module should be tested individually, which is called a “unit test” in software engineering.

##### Importance in SHM modeling task

Since the SHM features a multitude of feedback loops, locating errors was very tedious with the original monolithic model. Systematic debugging became only possible when we isolated the different parts of the system, such as the baroreceptors, and subjected them to controlled input signals to observe the component output. It was also possible to reuse several components within the SHM: The parasympathetic and the sympathetic system share a base class that only leaves the sign of the baroreceptor influence open for definition and the four different release equations for norepinephrine and acetylcholine are also governed by a common base class.

##### References

There is a consensus that multi-scale modeling requires some form of modularity for hierarchical composition [24, 25, 28, 43–45]. More specifically, Hellerstein *et al*. [40] and Mulugeta *et al*. [30] both suggest that object-oriented programming might be an especially promising way to implement modularity. Many researchers advocate for clearly defined interfaces [2, 28, 44, 46], but there is also critique with regard to a loss of flexibility for reuse and the requirement to consider all code-level elements of a model as potential coupling points [25, 29].

#### Human-readable

This characteristic covers two loosely connected aspects: The fundamental readability of model files with a text editor and the readability and understandability of definitions within the model.

Every modeling language has to be both human-readable so that a human can write the code to define a model and machine-readable so that a software tool can interpret that code to run simulations. However, as Figure 1 shows, there is a trade-off between the two and languages can choose to support the one at the cost of the other. On the one end of the spectrum, languages like Antimony or Modelica, whose syntax is closer to natural language and easier to read and write for humans using just a text editor, require more effort for specialized parsers to build abstract syntax trees, which can then be processed by compilers and other software tools. The middle ground is formed by XML-based languages like SBML and CellML. XML files already have a tree structure and parsers for XML exist for virtually all modern programming languages, which lowers the barrier to implement support for an XML-based format in a software tool and therefore increases interoperability between tools. While XML files can still be viewed and edited in a text editor, this requires familiarity with the language and tends to be cumbersome for larger edits. Especially the Mathematical Markup Language (MathML) format used both by SBML and CellML for storing equations can be hard to write and decipher without tool assistance. SBML and CellML therefore rely on software tools that use graphical interfaces or intermediary languages to ease model editing. On the machine-readable end of the spectrum, MATLAB Simulink uses a proprietary binary format that is tailored specifically to the MATLAB software toolchain. This can both reduce storage space and implementation effort for parsers, but also means that it is impossible to inspect model files without the corresponding software.

**Figure 1:**
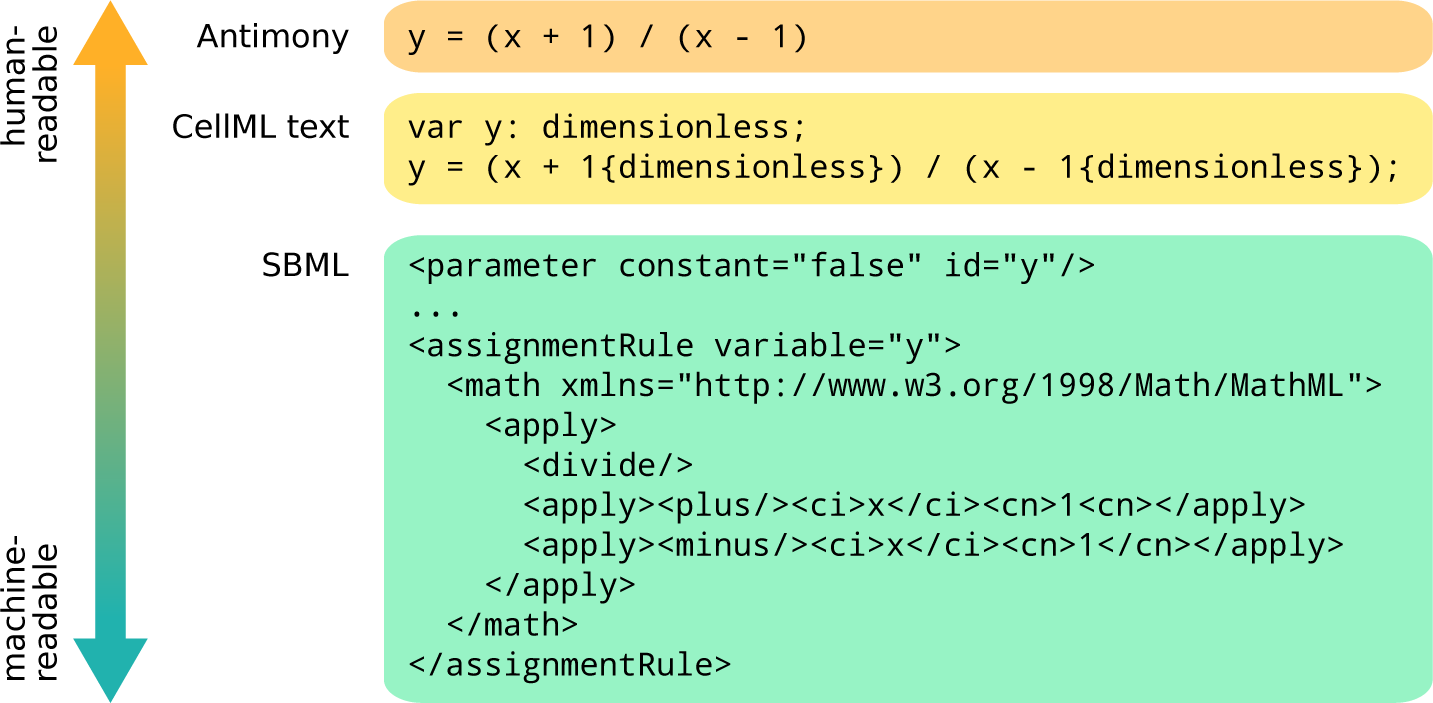
Example of the definition of a variable *y* depending on another pre-existing variable *x* via a simple assignment rule in three modeling languages with different levels of focus on human-readability versus machine-readability. Antimony mainly focuses on how humans would write equations in text form, but requires a specialized parser. CellML Text—an intermediary editing language used by the tool OpenCOR [47]—adds some syntax that is easy to parse by a machine (due to using braces that do not conflict with other symbols in the code), but is not an intuitive representation of unit information for a human unfamiliar with the language. SBML focuses more on machine-readability, since XML can be parsed by the standard libraries of most modern programming languages, ensuring minimal barriers for tool support. However, while the SBML code is still readable and editable in a text editor, it takes some effort and familiarity with the language to decipher the meaning from the symbols.

For model exchange and interoperability between different tools, XML-based formats have the clear advantage that supporting their import or export in a tool requires very little programming effort. This is illustrated by the success of SBML and FMI, which are both based on XML and are supported by over 100 tools each (http://sbml.org/SBML_Software_Guide/SBML_Software_Matrix, https://fmi-standard.org/tools). Increased interoperability also facilitates model reusability, because it becomes more likely, that a researcher who wants to reuse a model can simply import it in their tool of choice without having to translate it to another language first.

However, for model development and for publishing models to other researchers, languages with a strong focus on human-readability are preferable, because they allow tool-independent access to a model and because they are more suitable for version control. Due to their verbose syntax, XML-based languages are typically not designed to be written by humans directly but by software tools, which provide intermediary languages or graphical interfaces to facilitate editing. The translation between these different representations is performed automatically during export and import, which is convenient, but if the feature sets of the exporting and the importing tool do not overlap completely, there is a risk that information is lost. For example, a SBML model written with tool A may include layout information for a graphical representation of the model, but when it is loaded in tool B, which uses a purely equation-based representation, this layout information may be discarded. If tool B does not show a warning message, there is no way for the user to know that the model contained this information unless they look at the SBML code itself. This problem is less likely to occur, if the model is written in a language more focused towards human-readability, which is then also used directly for editing. In this case, both tool A and tool B would display the same code and while tool B does not display the graphical representation, the user would notice the presence of the layout annotations and could choose to view the model in a tool that does support them. Additionally, the more a language focuses on human-readability, the more easily it can be translated to slides, websites, articles and other formats, which makes it easier to communicate the details of a model to other researchers. It can also be archived more safely, as it will still be easily readable decades into the future, even if the tools used to create it and to view its contents will not be available anymore. Finally, version control software can be immensely helpful for tracking errors, for finding the exact versions of a model used to generate plots in an article, and for understanding the rationale behind modeling design choices. Standard solutions like Git operate under the assumption, that the documents under version control are written by humans and that element order, white space, and other details all are results of deliberate choices and therefore carry meaning. This is not the case for XML-based documents written by tools, which can artificially inflate changesets between document versions with management data or structural changes that carry no meaning and therefore obscure the changes that are actually relevant. While there are specialized solutions to distinguish semantic from structural changes in XML documents [48], this is still an active field of research and not yet broadly implemented in version control software. Also, even with these solutions, researchers might only be familiar with the tool that generated the file and not the content of the file itself, which makes it harder for them to localize and understand changes between model versions.

It is possible to combine the benefits of XML-based exchange formats and languages that focus more on human-readability, if these exchange formats are used and published *in addition* to a more human-readable representation of a model. This can be seen as analogous to software packages in general purpose programming languages. Open source software projects are usually both published as some kind of easily installable artifact–a file that not even has to be human-readable at all–and also as human-readable code in an online repository, which can be used to analyze and extend the software.

Moving from the question of the general file format to the content of the file, it can be said that readable code is largely the responsibility of its authors. However, a language may facilitate a clean coding style by providing expressive language constructs and documentation features or hinder it by introducing visual clutter. One example for this is the verbose use of {`dimensionless`} that is required after each constant in an equation in CellML Text as seen in Figure 1. Additionally, languages can also add human-readable documentation strings to variables and components or incorporate an HTML document for a more detailed model description. In contrast to comments in traditional programming languages, which are ignored by the compiler, these documentation features can enrich model presentation across various tools and representations including graphical dialogs or HTML representations within a model database. An example for this can be seen in Figure 2.

##### Guidelines

Model files stored in a version-controlled repository and published in model databases should be written in languages that focus on human-readability. If possible, models should additionally be published in a more easily machine-readable exchange format like XML to lower the barrier for direct reuse. If the language has support for structured documentation that is semantically tied to individual components or variables, this form of documentation should be preferred over unstructured comments. Every parameter, variable, and model component should at least be documented with a short human-readable label. Any non-obvious design choices or complex equations should also be documented.

##### Importance in SHM modeling task

On several occasions during our implementation we accidentally introduced errors in one part of the model while correcting an issue in a different part. To recover from these errors, it was crucial that we could quickly skip through the changes made since the last known working version. This was facilitated by the fact that Modelica focuses on keeping model files easily human-readable. With an XML-based format, we would have had more difficulties to make sense of the differences between versions.

##### References

Hellerstein *et al*. [40] and Zhu *et al*. [43] stress the importance of keeping model files under version control. The authors of Tellurium specifically state that human-readable languages can facilitate reproducibility and exchangeability [34]. Drager et al. [49] found that existing tools struggle to make all the information in the XML-based description of SBML models accessible in a comprehensive form, which led them to develop a tool called SBML2LaTeX, which generates human-readable reports from SBML models.

#### Hybrid

A language is *hybrid* if it supports multiple modeling formalisms and thus multi-class models. The most common form of hybrid models and languages cover both continuous ODEs or differential algebraic equations (DAEs) and discrete events, but other combinations are possible. The distinction between ODEs and DAEs is important here since physical conservation laws, such as conservation of mass or energy, are *algebraic* constraints, which cannot always be formulated with pure ODEs. Incorporating them in a model can, however, have the benefit of making connections between components *acausal*, which means that variables do not have to be designated as input or output and instead the solver can choose the appropriate resolution order. This avoids errors and performance issues related to algebraic loops, simplifies model descriptions, and allows reusing components in different contexts. As with DAEs, support for discrete events also comes in different forms. Many languages support the reinitialization of continuous variables through discrete events.

In this formalism the only discrete part of the model is a set of equations that define boolean values based on the state of the system. When these values switch from false to true, events are generated, which can introduce discontinuities in an otherwise continuous system. For a fully hybrid model that involves more complex discrete parts it is preferable that the language also supports the explicit declaration of discrete variables. The value of these variables remains constant between events, but they may be defined with complex equation systems, which are solved during each event instance. As a result, they can make models that require complex event triggers more understandable as can be seen in Figure 3.

**Figure 2:**
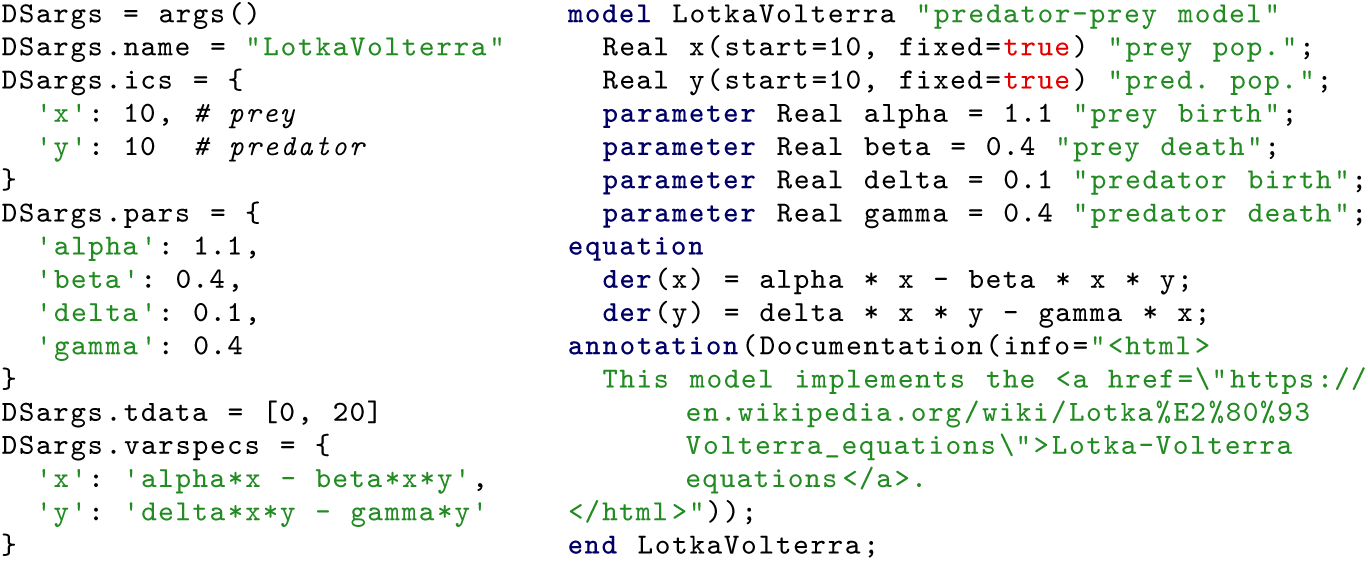
Simple predator-prey model in a language without (left, PyDSTool) and with (right, Modelica) support for documentation strings. Note that while regular Python comments (#) can be used to annotate PyDSTool models, they are ignored by the compiler and are only useful when reading the code directly. Modelica comments are part of the model syntax and can therefore be read by tools to, e.g., provide automated tooltips in dialogs and graphs or to enrich model summaries in databases.

**Figure 3:**
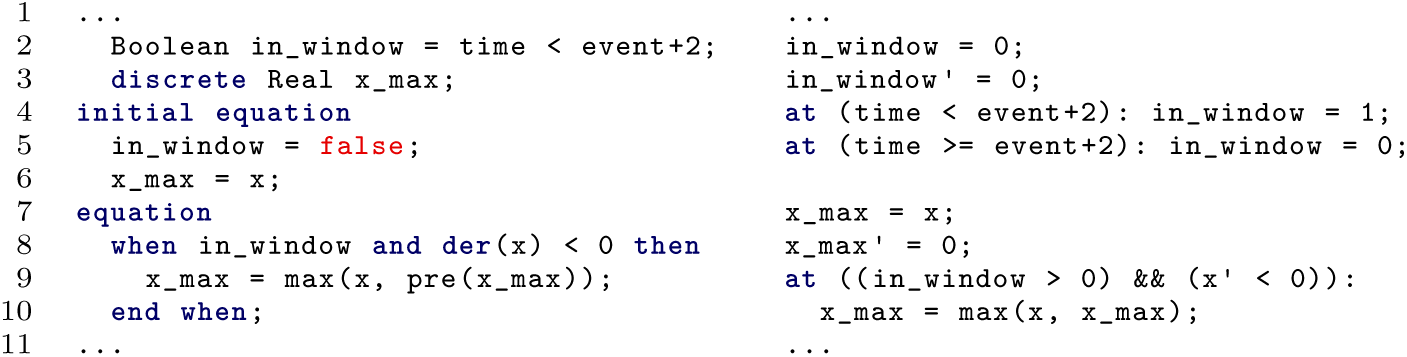
Definition of a discrete variable `x_max`, which captures the peak value of the continuous variable `x` obtained within two seconds after an event event in a language with (left, Modelica) and without (right, Antimony) support for declaring discrete variables. The Modelica model defines a discrete boolean variable `in_window`, to simplify the **when** condition later in the code. The information that this variable is discrete already lies in the type definition as `Boolean`. For real variables like `x_max`, there exists a keyword **discrete**, which determines that the variable value may only change within a **when** equation. The same model structure and semantics can also be achieved in Antimony, but the discrete variables `in_window` and `x_max` each need an explicit rate rule to ensure that their value only changes when an event occurs (lines 3 and 8). Additionally, two events are needed to emulate the boolean variable `in_window`: One to update the value when the condition becomes true (line 4) and one to do so when it becomes false (line 5).

It is important to note that we do not argue that a modeling language should support as many formalisms as possible, but rather a combination of formalisms that go well together. If other formalisms are required, the language should rather aim to allow coupling of models across languages with standardized interfaces such as the FMI [41, 42]. Additionally, there is a trade-off between fully supporting a modeling formalism, such as ODEs or DAEs and being able to assign a domain-specific meaning to language constructs. For example, SBML, PySB, and Antimony all use biological terms tied to the biochemical level to describe the parts of a model. This makes the language easier to understand and use for domain experts, but may prove challenging when building a multi-scale model that has to extend beyond the biochemical level.

##### Guidelines

A model should clearly indicate which variables are discrete and which are continuous. Event triggers, which define the transition between discrete and continuous parts of the model, should be examined and tested with extra care. If the language allows them, acausal connections should be preferred over causal input-output relationships, since acausality facilitates reuse.

##### Importance in SHM modeling task

At first, it was not clear for us whether the contractility of the heart in the SHM was a continuous or discrete variable. This confusion led to a severe error in an early version of the model. Our current implementation defines the variable with the keywords **discrete** `Real` to clearly indicate this distinction. Discrete variables were also required to disentangle the semantics of the cardiac conduction system, which is introduced in detail in Section 2.3.

##### References

In their 2017 review Bardini *et al.* [2] argue that multi-scale models in systems biology in general should strive towards a hybrid approach. The same has also previously been stated by other researchers [50–52]. In particular, the authors of PyDSTool argue that hybrid models based on DAEs are well-suited to represent multi-scale models [32].

#### Open

As a prerequisite for reproducibility and collaboration, models and simulation tools need to be accessible for everybody. In particular, the hurdle to run a quick simulation of a model to determine its usefulness for a specific task should be as low as possible. An openly accessible model definition also means that readers can offer feedback and corrections to improve the model. Preferably the language itself, the compiler and associated tools should all have an open-source license. Additionally, collaboration is also facilitated if the language can be used on different platforms.

##### Guidelines

Readers of a paper should be able to download the model code and to simulate it with open-source tools. The download should also include explicit licensing information. The model repository should include everything necessary to reproduce plots and other results of the corresponding paper. It should also be under version control and include a human-readable changelog. Other researchers should be able to point out errors and suggest corrections.

##### Importance in SHM modeling task

Without the reference code of Seidel, our re-implementation of the SHM would not have achieved a perfect agreement with the original. To weed out our last errors, which only showed quantitative and not qualitative differences in the plots, we needed to simulate both models with identical solver settings and manually compare the output data. Additionally, some small errors in the published formulas became only apparent when we compared them with their C implementation.

##### References

Many large projects and databases such as the Physiome Model Repository of the IUPS Physiome project [53], the NSR Physiome project [54], the BioModels database [27], the virtual liver network [55], Plants *in silico* [43] and SEEK [23] already provide open-source implementations of models. Mulugeta *et al.* [30] also specifically advocate for more version control and changelogs (in the form of e-notebooks).

#### Declarative

The mathematical formalism for biological models can already be complicated in itself. A modeling language should not require the adaptation of the model to the execution logic of the language, obscuring the original definition. Instead, the language should adapt to the model if it is presented in a clean mathematical formulation. This way the code can focus on expressing meaning rather than structure, which facilitates understanding. This also includes the possibility to formulate ODEs and DAEs not only in explicit form, i.e., with a single variable on the left-hand side of the equation, but also in implicit form, i.e., with arbitrary mathematical terms on the left and right-hand side. For example, specifying `u = r * i` should be equivalent to the equation `u / r = i` and the solver should decide for which variable this equation needs to be solved.

As a consequence of such a declarative style, errors reported by the compiler can also focus on meaning rather than just grammar, increasing the soundness of the model. One important example of this is that declarative languages usually allow the declaration and automatic checking of proper units for variables and parameters. Missing or wrong unit conversions are a common source of error in modeling that can be all but eliminated this way. Unit definitions also add semantic information and therefore make the model more understandable. Additionally, if the model is described in a declarative style, it is possible for automated tools to identify and extract meaningful parts of the model. This facilitates tool support—e.g., in the form of numerical solvers, optimization and verification toolchains—and also allows connecting model parts to standardized ontological terms. For the latter it is preferable if the support for ontologies is already included in the language itself.

##### Guidelines

Models should follow strict mathematical rules to fit nicely into the chosen formalism. If the language allows it, equations should be written exactly as one would write them in a scientific paper to convey their meaning, choosing freely between explicit and implicit form. If a model needs workarounds in the form of code that has to be added to make the model compile but that does not add new information about the modeled system, it may be worthwhile to revisit design choices and check the mathematical soundness of the model. In our own experience we found that most workarounds could be removed and the resulting model behaved more soundly and was easier to understand. Models should also specify units for all variables, preferably using the International System of Units (SI). If possible, automated unit consistency checks should be performed before publishing a model. Additionally, if the language supports semantic annotation with ontological terms, this feature should be used for all variables and components.

##### Importance in SHM modeling task

The original SHM was implemented in C, which is an imperative language. Most of the reference code that we consulted for our re-implementation was responsible for management tasks, such as storing a history of variables that enter equations with a delay, debugging output, or a manual implementation of an integration loop with the Runge-Kutta method. Although most equations were defined as separate functions, we sometimes had difficulties untangling the semantics of the model from the main integration loop.

One area of the model that was highlighted by the Modelica compiler as not mathematically sound was the systemic arterial blood pressure, which is given by an algebraic equation during systole and by an entirely separate differential equation in diastole. This issue only became apparent, because we had to translate the imperative C code, which simply used an if-expression to switch between the two states, into a declarative form, which required a consistent equation structure. This consistent structure could be established by manually differentiating the systolic equation and then only switching between two different expressions for the derivative.

##### References

Few researchers in systems biology explicitly distinguish between imperative and declarative languages. Zhu *et al.* [56] state that declarative languages are desirable, because it allows the description of the biological processes “in a natural way”. Several language authors also state that their respective modeling language is declarative [29, 33, 57, 58], but they do not explain why this is important. Of these, only the authors of JSim [58] and Myokit [29] state that declarative languages allow concentrating on what is modeled and not how the equations are solved, make models more understandable, and facilitate their analysis both by researchers and software tools.

#### Graphical

Discussing or even just understanding a model is difficult if the model is only described in the form of code or mathematical equations. This is especially true when the input of domain experts is required, who are not computer scientists or mathematicians. For this purpose most papers in biology use some kind of diagram to transport the general structure of the model in a graphical way. Here, there is a trade-off between two different visualization types:

1. Automatically generated abstract graphs of variable dependencies are an exact representation of the model and are well-supported by tools, which reduces the effort required to build these representations. However, automatic graph layout is a nontrivial problem: Different algorithms or parameter settings can lead to large differences in the layout [59]. Most algorithms also do not scale well to large graphs and additional techniques are required to group nodes according to semantic similarity [60]. Additionally, to the lack of grouping capabilities, automatic graph visualizations also solely rely on the variable names to convey the role of a variable—e.g., whether it is the product, reactant or catalyst of a reaction—or the kind of variable interactions—e.g., if the correlation is positive or negative. Consequently, this approach is mainly suited to represent the mathematical dependencies of variables, but not to give an intuitive overview of the model structure or to analyze the biological relations between modeled concepts.
2. Manual drawings of the biological interactions with respective images and symbols capture the essence of the information required to understand the model and can quickly be processed by the reader. This also has the additional benefit that the model can be discussed with domain experts that are familiar with the biological concepts, but not with mathematical modeling. However, they are less accurate, not standardized and require a lot of manual effort. This can also mean that when a model is extended or otherwise updated, changes may not be immediately reflected in the drawing, since it may only be updated at a later stage or not at all.

There are multiple hybrid approaches that try to address the shortcomings of pure type 1 or type 2 visualization. The Systems Biology Graphical Notation (SBGN) [61] allows the illustration of models with standardized abstract structural diagrams, which serve a similar function as circuit diagrams in electrical engineering. SBGN diagrams do not display variables, but represent the actual physical entities and processes with unambiguous, standardized glyphs. While there exist tools that can generate SBGN diagrams automatically, like CySBGN [62], some manual arrangement is required to produce satisfactory results. The standardization of SBGN also comes at the expense of the biological intuitiveness of the resulting diagram. Instead of immediately recognizable biological icons, researchers have to learn and interpret a series of abstract glyphs. For metabolism pathways this is no issue, since species that are part of a reaction are typically identified by their name and not by any two- or three-dimensional structure that could be used as an icon. However, e.g., for action potential models it would be preferable to represent ion channels and pumps by schematic drawings and to have a visual separation between the inside and outside of a cell. Such a graphical representation is especially helpful if it is standardized across different models. For example, the Physiome Model Repository [53] uses the same icons for ion channels and pumps across all curated action potential models, which are drawn by the now discontinued tool OpenCell (http://physiomeproject.org/software/opencell/about). A similar standardized icon language could also be beneficial for models at the tissue or organ level.

Like type 1 and type 2 visualizations, SBGN graphs are independent of the capabilities of the language with which the model was written. They are generated by tools that do not need to have any connection with the modeling language itself. Modeling languages can support them by referencing image files or XML files containing SBGN as part of the model documentation, but they have to be maintained separately. An example of this can be seen on the left side in Figure 4.

**Figure 4:**
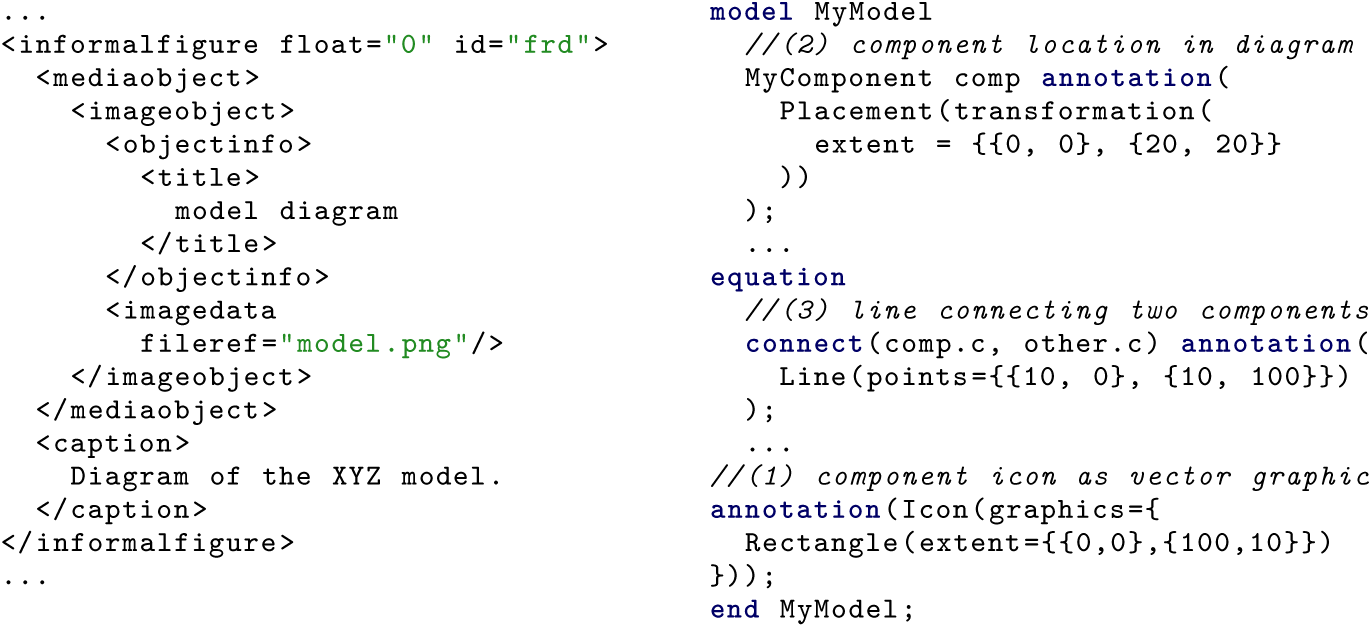
Two different ways in which modeling languages can support graphical representations of models as part of their syntax. Left: CellML allows to include diagrams or plots as figures in the model documentation. The image files remain separate from the model code and have no semantic connection to it except for the figure caption. Right: Modelica allows to add graphical annotations using a vector graphics syntax. Models and their components can have icons graphics (*//(1)*), which can be placed in a diagram coordinate system (*//(2)*) and connected with lines (*//(3)*). This graphical representation is tied to the structure of the model. If, e.g., a component is removed from a model, the placement annotation (*//(2)*) must also be removed, which automatically updates the diagram and ensures that it still accurately reflects the new model structure.

This is addressed by another hybrid approach that goes a step further towards the analogy with circuit diagrams and integrates layout and rendering information directly into the model structure. It is mainly prevalent in languages with an industrial background such as Modelica and MATLAB, but is also implemented in the SBML level 3 layout and rendering packages [63, 64]. In this approach, model components are assigned graphical annotations, which define how the component should look and where it should be placed in the diagram representation of the model. In a modular language, this information can be used to build tools that allow to construct models by dragging and dropping component icons and connecting them with lines, much like a circuit diagram. An example of this can be seen on the right side in Figure 4. The resulting diagrams are both an accurate reflection of the connections between model components, because they are intrinsically tied to the functional model code, and can be understood quickly, since they are arranged manually and use biological imagery. If the model changes and, e.g. a component is removed, the graphical annotation also has to be removed, because the compiler would otherwise produce an error message. This ensures that graphical representations stay up to date when a model is changed. Creating symbols and images for components requires effort, but this has to be done only once for each component and the arrangement and connection may even be easier than writing the equations that connect the components in code.

As becomes apparent, this last approach should be favored for multi-scale models, although it has to be noted that it is also possible to combine multiple approaches in the same language.

##### Guidelines

All interactions between the individual modules in a model should have a graphical representation in the corresponding diagram. Each diagram should only have a few components. If it becomes too crowded, some components should be grouped together to form a hierarchical structure. Each individual component in the diagram should be represented with an intuitive symbol that either corresponds to the appearance or function of its biological equivalent. Components should be visually grouped according to their function and interaction to facilitate understanding.

##### Importance in SHM modeling task

The original SHM features a graphical representation in the form of 23 text boxes that are connected by arrows. While this does give an overview of the physiological effects present in the model, one of our first steps to better understand the model was to augment this diagram by grouping the effects by the organs to which they belong and adding respective icons. Our Modelica implementation now features a fully visual diagram with 15 components that is guaranteed to be faithful, since it is tied to the equations in the code. It helped us on several occasions to discuss the model with domain experts, such as physicians and chemists.

##### References

The Physiolibrary is a Modelica library for physiological models that has graphical representations for each component [38]. ProMoT is a modeling tool that allows the composition of modular models in a graphical way [44]. Alves *et al.* [65] compare 12 different simulator tools, giving higher ratings to those that have graphical representations for model components. Mangourova *et al.* [66] state that it is preferable when a modeling tool for integrative physiology provides a graphical way of composing models since this can reduce development time.

To easily refer to these characteristics we form the mnemonic MoDROGH for Modular, Descriptive, human-Readable, Open, Graphical, and Hybrid. We will also use the term “MoDROGH language” for a language that exhibits all or most of these characteristics. A checklist for ensuring that a particular model utilizes the MoDROGH criteria to the fullest extent permitted by the chosen language can be found in Supplementary Note 3.

### 2.2 Existing languages exhibit MoDROGH criteria to varying extent

As mentioned in the introduction, there are multiple suitable languages available that implement the MoDROGH characteristics to some extent. In this section the most prominent examples will be discussed with respect to each characteristic (highlighted in *italics*). We consider a “modeling language” to be any language used for describing and distributing mathematical models. This includes exchange formats such as CellML and SBML, languages that are embedded in a general-purpose programming language like Python or Julia, and standalone languages like Modelica. We selected languages that are currently popular either in systems biology or in mathematical modeling in general with a tendency towards general-purpose modeling languages that are not restricted to a specific organizational level or model type. We also only chose candidates that exhibit at least some MoDROGH characteristics. We have to emphasize that the list is not comprehensive, but we tried to cover examples for all major trends in modeling languages.

#### MATLAB

MATLAB is perhaps the most widely known language used for solving ODEs (https://www.mathworks.com/products/matlab.html). The MoDROGH criteria can be best fulfilled when using the Simulink environment (https://www.mathworks.com/products/simulink.html) with the embedded language Simscape (https://www.mathworks.com/products/simscape.html). The SimBiology environment can be used as an alternative, which is comparable to SBML in its expressiveness regarding rules and reactions and can also export models to SBML (https://de.mathworks.com/products/simbiology.html). It is, however, tailored towards pharmacological models and not as feature-rich as Simulink and Simscape, which is why we restrict our analysis on the latter combination. Simscape realizes *modularity* through full object-orientation with class definitions, instantiation and inheritance, although Simscape classes can only have one parent class, in contrast to MATLAB classes, which allow multiple inheritance. Through Simulink, models can be imported from different languages using the FMI, but export of Simscape models with this interface is currently not supported. The language is also *declarative* allowing to freely mix between implicit and explicit formulation of DAEs, which are written in a concise syntax that focuses on *human-readability*. It supports documentation strings for components and human-readable labels for variables. Unfortunately, the readability of Simscape is hampered by the fact that Simscape has to be used in conjunction with Simulink, which saves models in a proprietary binary format, which is not readable in a text editor and not even openly documented. This issue is further aggravated by the fact that backward compatibility to older versions of the model format is not guaranteed [66]. Units are supported and a mandatory consistency check is performed at the interfaces between components. There is no built-in support for ontologies, but since Simscape supports object-orientation, “is-a” relationships, which designate a component as an instance of a concept, might be expressed by building a large type hierarchy of ontological terms. This would require all models to use this type hierarchy and therefore reduce flexibility in designing generic base classes, since Simscape only allows single inheritance. Simscape classes can be used as *graphical* components within Simulink to create larger systems by arranging and connecting them via drag and drop. *Hybrid* systems are supported with index-reduction for DAEs and discrete events and variables. Unfortunately MATLAB, Simulink, and Simscape are proprietary tools that are not *open* in any way, requiring license fees and prohibiting custom extensions.

#### SBML

In systems biology, the SBML is a widely used *open* language for describing biological models—mostly at the level of biochemical pathways [67]. SBML level 3 includes an optional language module for hierarchical composition, which allows building *modular* models via the import of components during which individual variables can also be overwritten or deleted. Because it uses a subset of MathML to describe equations, SBML is *declarative* and *hybrid* and in theory allows the definition of arbitrary DAEs in explicit and implicit form. SBML is based on XML, which makes it highly machine-readable and in turn facilitates interoperability between tools, because support for model import or export can be implemented easily. Unit definition is possible but optional and tools are not required to interpret them. However, libSBML, the most popular library for working with SBML models, can perform automatic unit consistency checks [68]. Support for discrete events is limited to reinitialization of continuous variables. The reliance on MathML and XML is also a drawback, because it limits the *human-readability* of model files and presents challenges for version control software that is not equipped to distinguish structural from semantical changes. Individual components can be annotated with textual notes, Systems Biology Ontology (SBO) terms, or Minimal Information Required In the Annotation of Models (MIRIAM) metadata. Using the SBML level 3 packages for layout and rendering, *graphical* annotations can be assigned to model components. The high interoperability between SBML tools resulting from its focus on machine-readability is a major advantage, because researchers can use a tool that is designed to fit their specific use case and can reuse models across tools. Due to the wide acceptance of SBML, it can be expected that most researchers will have at least one such tool available so that the visual clutter of the XML files is no issue for model reuse. However, most of these tools do not support all optional SBML packages with the consequence that in practice support for modularity, graphical annotations, and DAEs in implicit or explicit form may be limited to specific tools.

#### CellML

CellML is similar to SBML, but focuses on building more general component-based models [69]. It is also *open*, *declarative* and *hybrid* with the same considerations for being based on XML and MathML. In contrast to SBML, it does not only support units, but enforces that every variable in a valid CellML model must have a unit definition. Modelers can still choose the special value dimensionless to designate that a variable does not have a unit, but they have to make this choice consciously and explicitly. The language itself does not require tools to check the consistency of these units, but OpenCOR, one of the main tools for creating and simulating CellML models, does perform automated consistency checks when a model is loaded or saved [47]. OpenCOR can also somewhat alleviate the downside in *human-readability*, because it defines a so called “CellML Text” language, which can be used to view and manipulate the model in a more human-readable text format [47]. However, “CellML Text” has limited expressiveness only allowing the definition of explicit and not implicit equations and it is only used for viewing and editing and not for model storage. It also does not contain annotations, which can be defined in CellML through embedded metadata files in Resource Description Framework (RDF) format, which can also contain ontological annotations. Since version 1.1, *modular* CellML models can be hierarchically composed of sub-models [24, 70]. To support the graphical representation of models, CellML provides constructs for referencing externally-stored graphical files and formatting figure captions. This feature allows modelers to link models with an associated image and is used by the curators of the Physiome Model Repository—the primary clearinghouse for CellML models—to display relevant figures on a model’s webpage. However, as there is no semantic link between figure elements and model code, it is the responsibility of the modeler to keep the figure up to date when the model is changed.

#### Python

Python is an open-source programming language that is popular in data science (http://www.python.org). The language itself is imperative, but it can be extended with some declarative features for special purposes. In systems biology, notable efforts include PySB [33] and the PySCeS [31]. These packages define their own declarative domain-specific languages (DSLs) within Python to tackle specific biological use cases. PySB focuses on rule-based reaction models while PySCeS focuses on ODEs, structural analysis, and metabolic control analysis. There also exist general-purpose packages, such as SimuPy [71] and PyDSTool [32], that allow users to create and analyze models built with ODEs, DAEs and discrete events.

All aforementioned python-based solutions are *open* and *declarative* and Python itself focuses on *human-readability*. However, the modeling packages mainly rely on the modeler to use the features of Python to implement *modularity* concepts and to document their models by themselves. Also, none of them support any *graphical* representation of models. Notably, SimuPy and PyDSTool lack slightly in human-readability and declarativeness because they require a very specific and low-level technical format for defining equations. Exceptions in terms of modularity are SimuPy’s block diagrams and PySB’s macros. The fact that the environment is not declarative in itself leads to the drawback that only PySB supports ontological annotations and only PySCeS supports the definition (but not consistency checks) of units. Regarding the *hybrid* characteristic, the differences are most pronounced since PySB is not hybrid at all, featuring only specific biochemical rules without events, while PySCeS and SimuPy allow discrete events as well as ODEs and only PyDSTool is able to also handle DAEs. None of these packages support the explicit declaration of discrete variables.

#### Antimony

Antimony [35] is a *declarative* modeling language with an emphasis on *human-readability* used by the *open* Python-based environment Tellurium [34], which can be used for model building, simulation and analysis. Since Tellurium version 2, Antimony also supports the structural annotation of models with terms from the SBO or general MIRIAM metadata. Antimony is *modular* by design, allowing the definition of components that can be imported in other models. As in SBML, individual variables and equations can be overwritten or deleted during import. It is *hybrid* in the sense that it allows discrete events, but it only supports explicit ODEs and not DAEs and it lacks support for declaring discrete variables. Like SBML, Antimony focuses on models on the level of biochemical pathways by providing a special syntax for reactions. It has no support for embedding any form of *graphical* model representations.

#### Modelica

Modelica is an open-source declarative modeling language primarily used in engineering [37]. It is very similar to MATLAB’s Simulink environment and the Simscape language. In fact, Simulink was developed before Modelica and Modelica before Simscape, which suggests some influence between the languages in both directions. Modelica supports *modularity* via object orientation including the overwriting of variables and explicit equations, and, in contrast to Simscape, multiple inheritance. Most Modelica tools support the FMI, allowing the reuse of models across different languages. Like Simscape, it also allows grouping of interface variables to connectors, which can be used to connect models *graphically* via drag and drop. It is *human-readable* and *declarative* allowing to define a model with a mix of explicit and implicit DAEs. Models can be annotated with documentation strings for individual components, a full HTML documentation for classes and machine-readable annotations, which do not support ontologies by default but have a flexible extension mechanism with so-called vendor-specific annotations. As in Simscape, “is-a” relationships between model components and ontological terms can also be implemented via a type hierarchy. While this introduces design restrictions in Simscape, Modelica supports multiple inheritance and therefore allows maintaining ontological type hierarchies in addition to generic base classes. Units are supported and optional consistency checks can be performed. Modelica is also fully *hybrid* with support for discrete events and variables as well as arbitrary DAEs. With additional, open-source libraries, it is also capable to express models, e.g., as bond graphs, Petri nets and finite state machines. The Modelica ecosystem is a mix of *open* academic tools and commercial tools for use in industry while most libraries are open-source. There are two actively maintained open-source Modelica compilers called JModelica and OpenModelica, the latter including a fully-fledged integrated development environment (IDE) [72, 73]. However, Dymola, the most widely used Modelica IDE, is proprietary and not fully compatible with opensource alternatives (https://www.3ds.com/products-services/catia/products/dymola/). Therefore, Dymola models may need to be adjusted slightly to run with open-source compilers or vice versa.

#### Julia

Julia is an *open* programming language that is mainly used for data science [36]. It is imperative by nature, but the language can be extended with macros, which are more powerful than the respective capabilities of Python, to allow *declarative* modeling. Elmqvist *et al.* use this feature for Modia, an implementation of the Modelica syntax within Julia [74]. Modia is currently still experimental. It is *modular*, *hybrid*, and focuses on *human-readability*, but lacks, for example, the *graphical* features of Modelica.

In general, Julia offers strong support for differential equations with packages such as DifferentialEquations.jl which supports *hybrid* systems including DAEs, partial differential equations (PDEs) and discrete events, but not the declaration of discrete variables [75]. The syntax of this package focuses on *human-readability* and is *declarative*, but only allows either a fully-implicit or semi-explicit formulation of the whole system of DAEs with a mass matrix. Like Python-based solutions, annotation and *modularization* in this package is up to the modeler using the features of the language Julia which supports a multiple dispatch mechanism, which can be used to accomplish the same functionality as object orientation except for encapsulation. However, units with automated consistency checks can be supported though the Unitful.jl package [75]. Like Modia, DifferentialEquations.jl offers no *graphical* representation of models.

A summary of the available languages and their features can be found in Table 1. Notably, Simscape and Modelica stand out by supporting full object-oriented design of models, explicit declaration of discrete variables, an integrated graphical representation, which allows biological drawings and manual arrangement, acausal connections between components, cross-language import of models via the FMI, grouping of interface variables as connectors, and unrestricted mixing of implicit and explicit equation formats. The feature-richness of these languages is not surprising, since both are established industry standards, which are used in multiple disciplines to build large and complex models. Between the two candidates, Modelica additionally provides a mostly open environment, multiple inheritance, overwriting of variables and some equations during instantiation and inheritance, export to other languages via the FMI, and machine-readable annotations, which can, in theory, be used to implement support for ontologies. On the downside, this ontology support must be implemented manually and is not included in major tools and while open source tools do exist, they only make up a part of the Modelica ecosystem and are not necessarily fully compatible with proprietary solutions.

**Table 1:**
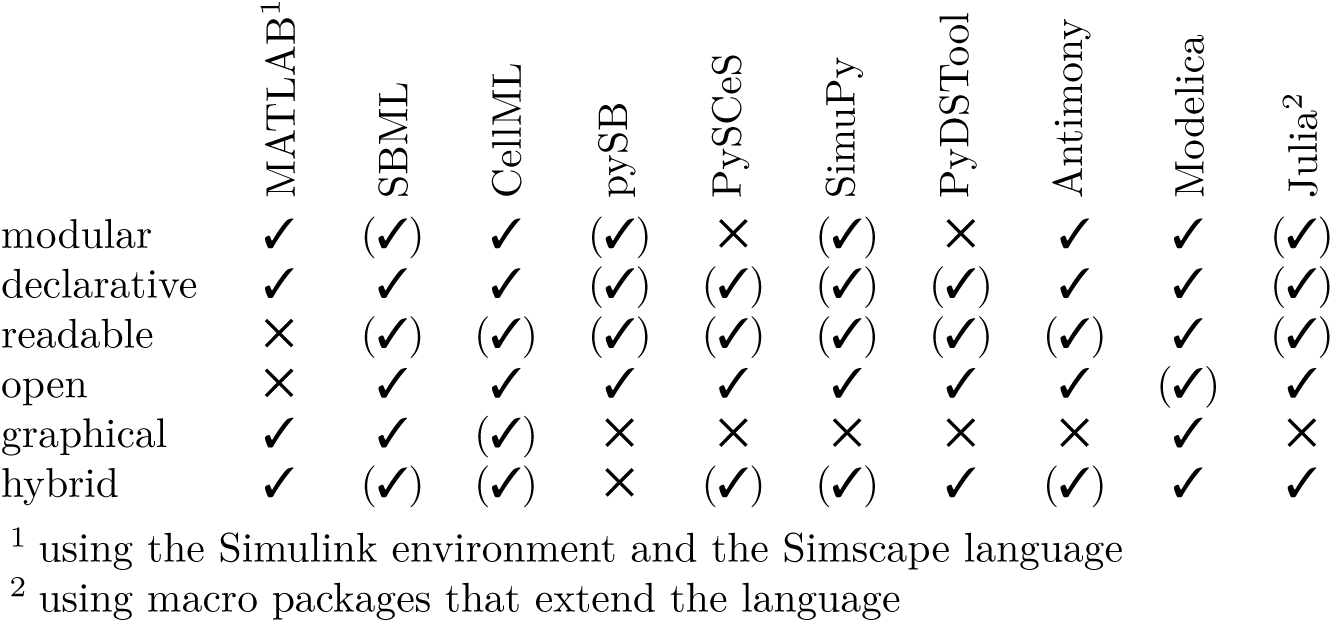
Evaluation of language candidates with respect to the desirable characteristics established in this paper. A check mark in parentheses means the language has the respective characteristic in principle, but not to its full extent or with noticeable drawbacks. A more detailed version of this table with regard to individual language features can be found in Supplementary Table 1.

Although it is certainly not the only option and it is as of now foreign to the systems biology ecosystem, we think that Modelica is a suitable choice to demonstrate the benefits of the MoDROGH characteristics since it implements them to the fullest extent among our selection of languages.

### 2.3 Modularizing a model of the human cardiac conduction system facilitates reuse

The Seidel-Herzel model (SHM) describes the autonomic control of the heart rate in humans at a high level of abstraction [10]. It was developed and implemented by Henrik Seidel in 1997 using the programming language C. We chose this model because preliminary^1^ versions, which have been published as individual peer-reviewed articles [11, 76], have gained substantial research interest and are able to simulate several relevant disease conditions such as first and second degree atrioventricular block [10], carotid sinus hypersensitivity [11], congestive heart failure [77], and primary autonomic failure [77] as well as treatment options such as the administration of atropine or metoprolol [77]. It is also especially interesting with regard to its dynamical properties such as the emergence of Mayer waves [10], bifurcations [11] and cardiorespiratory synchronization [78]. In a previous paper, we translated the SHM to Modelica [12], and we recently also published our full model code as an open-source reference implementation [79]. The model is therefore freely available, able to produce physiologically relevant results, large enough to benefit from engineering methodology, and yet small enough to allow an in-depth analysis at the source code level. It is not representative of lower-level metabolism and cell-signaling models, which are currently the most common type of models encountered in systems biology, but it is well suited to showcase what is needed for future multi-scale models, which inevitably have to leave these well-explored levels behind to generate new insights. In fact, the model can already be considered to span multiple scales of time since it includes effects at the sub-second level as well as on the level of multiple minutes [80].

To be more specific, the SHM can be classified as a hybrid (discrete and continuous), deterministic, quantitative, macro-level model. All effects in the model are described on the organ level, including the time course of systemic arterial blood pressure generated by the pumping of the heart; the Windkessel effect of the expanding arteries dampening the initial rise in blood pressure; the arterial baroreceptors generating a neural signal depending on the absolute value and the increase in blood pressure; the autonomic nervous system emitting norepinephrine and acetylcholine as hormone and neurotransmitter based on signals from the baroreceptor and the lungs; and the cardiac conduction system with the sinoatrial node (SA node) as main pacemaker and the atrioventricular node (AV node) as a fallback system.

In the following, only the conduction system is examined. It takes an input signal from the SA node (based on norepinephrine and acetylcholine concentrations) and includes the refractory behavior of the SA node^2^ limiting the maximum signal frequency, the delay between a signal from the SA node and the actual ventricular contraction, and the AV node generating a signal if no signal has been received for a given period of time. In the original model, these effects were tightly coupled within a single piece of code comprising five parameters, and 13 variables and equations—not counting additional parameters and variables for initial conditions. We found that this complexity makes it hard to understand and modify the model, which is why we translated it into a modular structure using Modelica. We will explain Modelica-specific language constructs as they appear in the code examples, but for a more complete introduction to Modelica in a biological context the reader is referred to the Lotka-Volterra examples in [81] as well as our own implementation of the Hodgkin-Huxley model [82].

The modular version separates the code into the three components `RefractoryGate`, `Pacemaker` and `AVConductionDelay`. These components are connected via a unifying interface using a base class `UnidirectionalConductionComponent`, which takes a Boolean signal as an input and produces a Boolean output. These inputs and outputs are only `true` for the exact point in time when a signal is issued (i.e., , they behave as a sum of Kronecker deltas). In Modelica, this behavior can be indicated by defining a type alias `InstantSignal`.

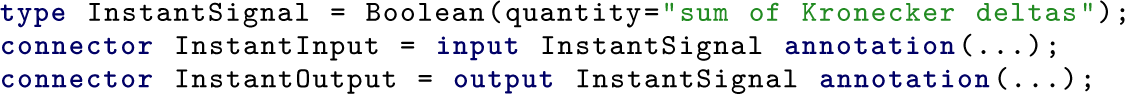

The new type is functionally identical to the base type `Boolean`, but by overwriting the built-in variable `quantity` it includes additional information that is both human-readable and can be interpreted by graphical tools to enhance understandability. The next two lines achieve two separate goals: First, the keyword **connector** designates `InstantInput` and `InstantOutput` as part of the interface of a class to the outside world. Second, specifying the **input** and **output** causalities ensures that input signals can only be connected to output signals and vice versa. This distinction can also be reflected in the graphical representation, which is defined in **annotation()** statements, which are shown here without their content for the sake of brevity. The base model `UnidirectionalConductionComponent`, which has one input and one output, then becomes

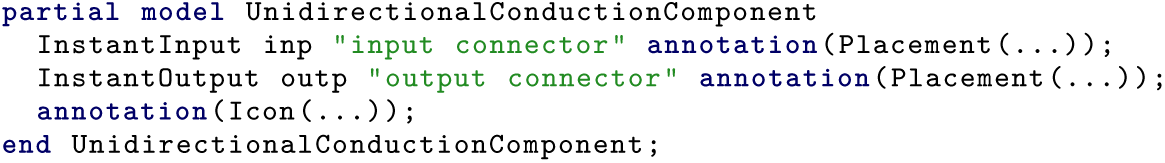

Note that the model is declared as **partial** which indicates that it is only a template that cannot be used on its own but must be extended by defining other models that include the following declaration.

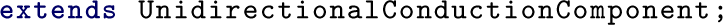

This statement imports all variables and equations of the base class into the current model, which ensures that all components will have an input and output connector named `inp` and `outp` without the need to define these variables multiple times. Models can also inherit graphical annotations from base classes, which can define a common look and connector placement for the graphical representation.

The three main components `RefractoryGate`, `Pacemaker` and `AVConductionDelay` all extend `UnidirectionalConductionComponent`. For the sake of brevity we will only show the code for the `RefractoryGate` here while the code for the other two components can be found in Section 4.2. The `RefractoryGate` represents the refractory behavior of the SA node which cannot be excited for a certain time period after it has fired a signal. For our model this means that the output equals the input except that after each signal there is a time period `d_refrac` in which incoming signals are ignored. This results in the following definition:

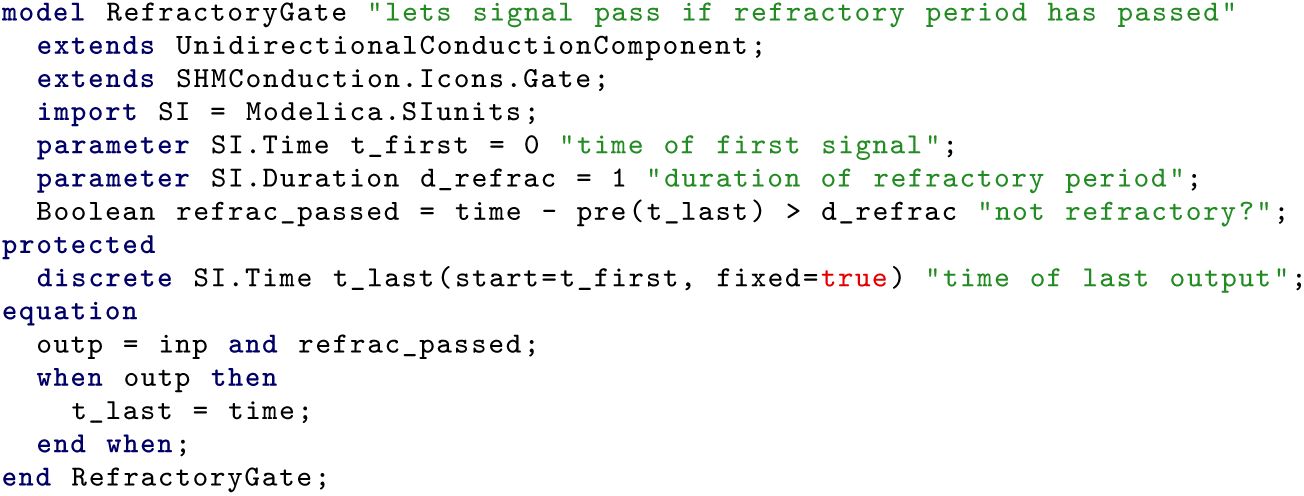

This model showcases several language features: It designates parameters with the **parameter** keyword, indicating that their value will not change during the simulation. It uses the `Modelica.SIunits` package, which contains types with unit definitions according to the SI. It documents each variable with a short informative explanation. It defines the helper variable `t_last` in a **protected** environment, which indicates that this variable is only relevant inside this component and should be hidden from other components. It contains an event using the **when** keyword, which can be used to assign values to discrete variables and to reinitialize continuous variables. It uses the `pre()` function to distinguish between the value of `t_last` *before* and *after* the event, which is required, because equations do not assume any causality. It explicitly marks `t_last` as **discrete**, which ensures that it must be defined within a **when** equation and indicates to the reader that it remains constant between events. It also employs multiple inheritance by including two **extends** statements: one for the base class containing the interface connectors, and one for an icon class containing the graphical annotation code. The latter is not strictly required, but it is convenient for readability, because it allows keeping verbose icon annotations in a separate file.

The other two components follow a similar design structure. The `Pacemaker` represents the capability of the AV node to generate spontaneous action potentials in the absence of a signal from the SA node. This means it lets incoming signals pass through but also issues a signal on its own when the output has been silent for the duration of its period `period`. To ensure that signals during the refractory period do not prematurely reset the pacemaker, it needs a separate reset input, which is only triggered when the output signal has also passed the refractory gate. The `AVConductionDelay` represents the time delay that occurs due to the slow conduction between AV node cells. It delays an incoming signal by a duration that depends on the elapsed time since the last output signal has been issued. As mentioned above, the code for both of these components can be found in section 4.2.

To form the full model of the cardiac conduction system, the components have to be connected through their interface variables. In Modelica, this is usually done in graphical tools like OpenModelica through a drag and drop interface. For this, the aforementioned **annotation()** statements come into play. They define the icons and the placement of components and connection lines in a vector graphics format. An example for the placement of the `inp` connector may look as follows:

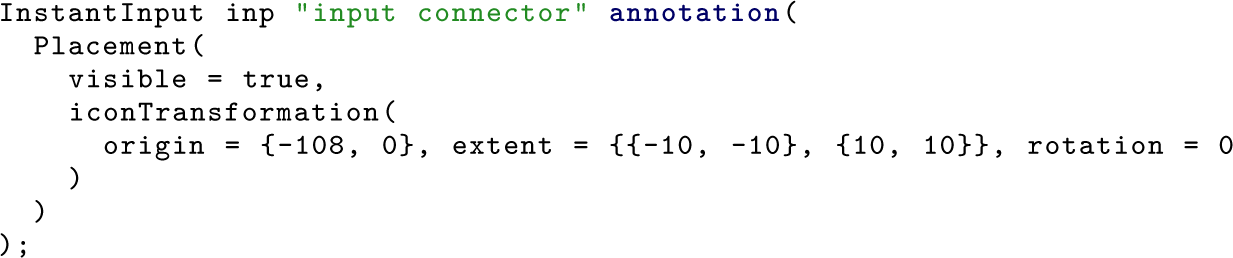

This ensures that the resulting diagram in Figure 5 is not a separate image file that has to be maintained separately, but is instead directly tied to the actual model structure. To keep the model code simple and short we defined the icon annotations in separate classes whose code can be found in Supplementary Listing 23–27. As seen in Figure 5 we chose an open fence gate for the refractory gate, a metronome for the pacemaker and an hourglass for the delay. The components are simply connected in order with the exception that the reset of the pacemaker component is only triggered if the signal also passed the refractory component. The resulting composite Modelica model `ModularConduction` looks as follows:

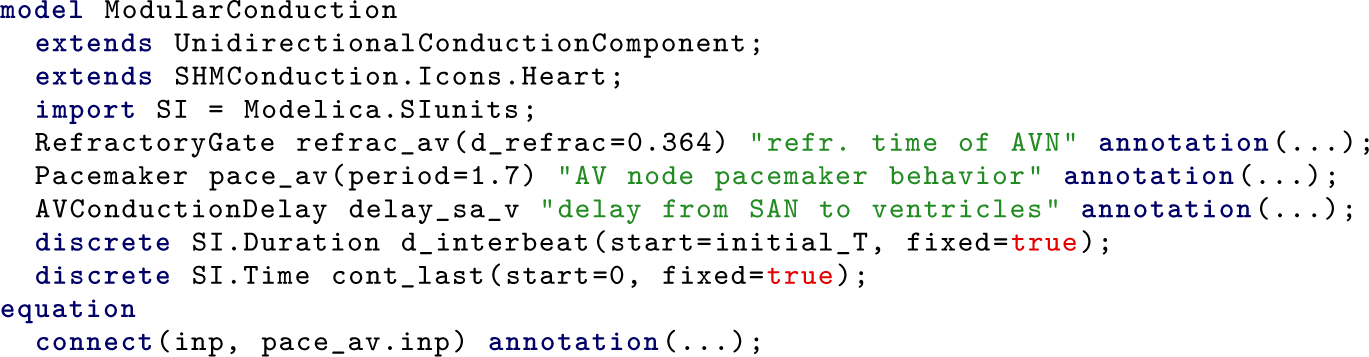
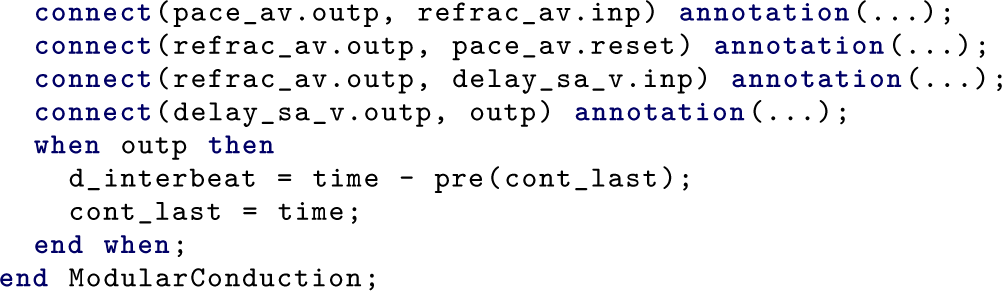

**Figure 5:**
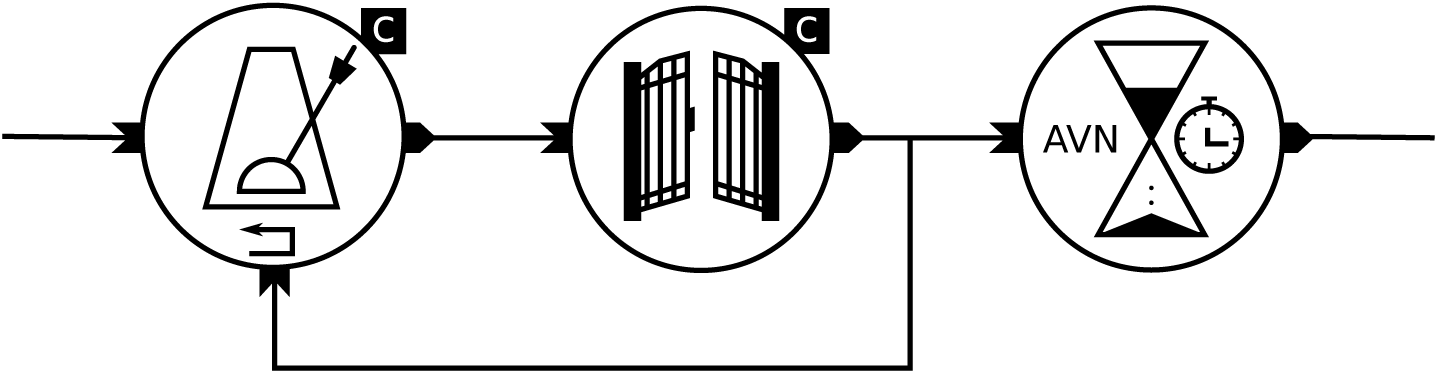
Diagram of the modular conduction model with symbols for the components. From left to right: `Pacemaker` for the pacemaker effect of the AV node, `RefractoryGate` for the refractory behavior of the SA node and `AVConductionDelay` for the combined delay between the SA node and the ventricles. The C in a black box indicates that the main variable of the component is held constant while the stopwatch symbol for the delay should indicate that the duration is time-dependent. Components have their input on the left, their output on the right and the pacemaker has the additional reset input at the bottom.

Note that we do not show the content of the **annotation()** statements here for the sake of brevity. The full code can be found on GitHub and in Supplementary Listing 1–27. Since the model itself receives a Boolean input from the SA node and provides a Boolean output for the Ventricles, it is itself a `UnidirectionalConductionComponent`. Components are used by defining variables of the types `RefractoryGate`, `Pacemaker`, and `AVConductionDelay`. The definitions also overwrite the parameters `d_refrac` and `period` to adjust the general `Pacemaker` and `RefractoryGate` models to their specific use case in this model. The inputs and outputs of the components are connected via **connect()** equations. In this case, **connect**`(a, b)` is synonymous with the equation `a = b`, but more complex connectors can connect multiple variables within a single statement and can also handle conservation laws. The model also introduces the additional variable `d_interbeat`, which allows using the interbeat intervals as a higher-level feature.

The structure defined in this model (and seen in Figure 5) deviates from the original SHM because the refractory behavior is situated at the AV node instead of the SA node. Additionally, the delay component models the complete delay from the SA node to the ventricles but is actually applied *after* the components for the AV node. To remain closer to physiology, one could split the delay component into two delays—one before and one after the AV node—and similarly add another refractory gate for the SA node. However, in Supplementary Note 1 we show that this simplified structure closely replicates the behavior of the SHM and even reveals some minor inconsistencies in the original model.

We also used our modular version to implement the trigger for PVC, which initially uncovered the problems with the monolithic version. It turned out that this extension now becomes possible without much effort, since it is easier to determine the effect of a PVC on the individual components one by one than to describe its effect on the whole system at once. A complete discussion of the extension can be found in Supplementary Note 2 and a diagram of the resulting model can be seen in Figure 6.

**Figure 6:**
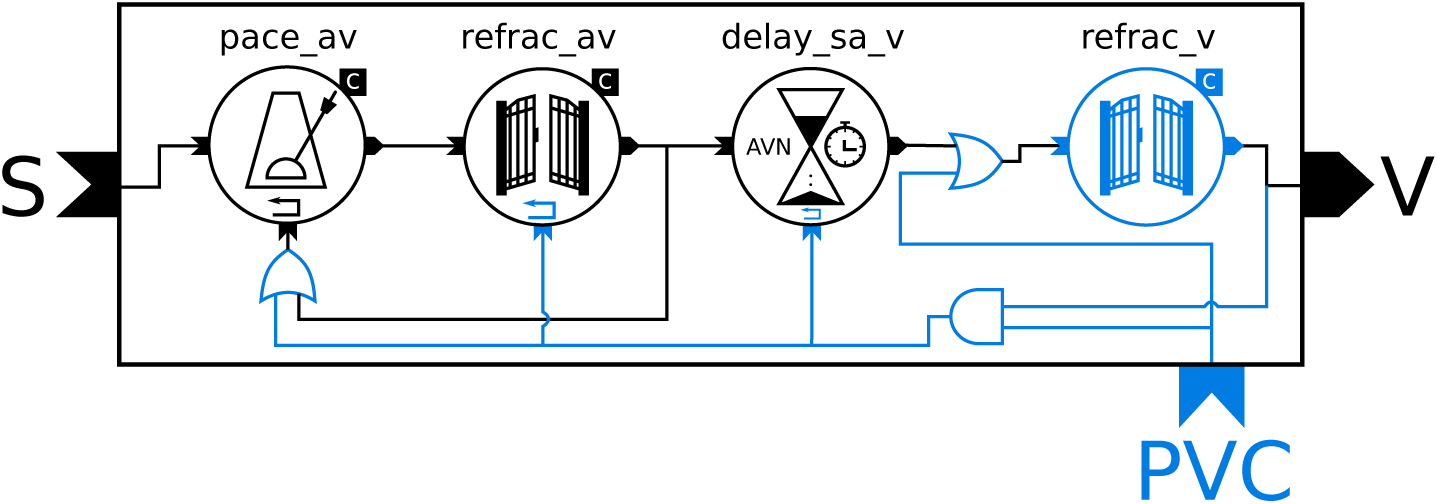
Diagram of the PVC model. The components are the same as in Figure 5 with additional components and connections highlighted in blue: reset inputs, second `RefractoryGate` (right) for the refractory period of the ventricles, two logical OR gates and one AND gate. The letters on the outside of the rectangle represent the connections of the model to the outside world: the input from the SA node (S), the output to the ventricles (V) and the trigger signal for PVCs (PVC).

## 3 Discussion

The model that we chose to demonstrate the benefit of the MoDROGH characteristics is quite small as compared to, e.g., , current whole-cell models, which can involve 28 or more individual interconnected components [83]. It can be argued that one needs to look at models of this scale to really assess the impact of model engineering decisions and language choice. However, we think that more than the size or structure of the model, the context of its reuse is the most important factor that allows us to generalize our findings to different areas of mathematical modeling. To extend the SHM, we needed to identify the correct integration points for the new effect in the model, which in turn required us to first separate the model into modules that each represent only a single physiological effect. We think that this requirement to understand and break up existing models for reuse in a different context represents one of the main challenges of multi-scale modeling in general. Additionally, larger models would not allow an in-depth discussion of their code within a single research article, since there would be simply too much interrelated code to discuss. We therefore chose the cardiac conduction system of the SHM as a “minimal working example”, which is just large enough to show the effects that we want to discuss but still small enough to cover the whole code in this article. This is in accordance with common practices in computer science textbooks where general design patters for the construction of large software systems are discussed based on small examples [84]. It is also important to note that the language Modelica and the techniques that we discuss here can be, and to some extent have been, applied to build large models. Examples include the Physiolibrary, a library to build multi-organ or whole-body circulatory models [38], the Guyton model of physiological regulation [85] or even larger examples from industrial settings such as end-to-end simulation of launch vehicles [86] or electrical power systems with thousands of components [87]. For our specific example, an application to a model of relevant size is also tangible, because our model of the cardiac conduction system can be seamlessly integrated in our Modelica implementation of the full SHM, which features 15 interconnected components and also utilizes the MoDROGH characteristics [12].

This leads to our initial research question to assess how the modular, declarative, readable, open, graphical and hybrid nature of a MoDROGH language helped in the modeling process of the conduction model. We discuss this for each individual characteristic and then sum up the impact on the model design goals of reproducibility, understandability, reusabilty, and extensibility and reflect on our choice to use the modeling language Modelica.

### Modular

The modular implementation of the cardiac conduction system consists of small components with at most three parameters and seven variables and equations including only two to three interface variables. This stands in contrast to the five parameters and 13 variables and equations of the monolithic version. The lower amount of items that have to be processed at the same time can be seen as an indicator for increased understandability [88]. The `Pacemaker`, `RefractoryGate` and `ConductionDelay` models all are quite generic and it is easy to imagine that they could be reused in a different model that requires these effects. This also facilitated the extension of the model with a trigger for PVCs, which required the incorporation of a second `RefractoryGate` to model not only the refractory behavior of the SA node but also of the ventricles (see Figure 6). In this case, the component could be reused without modification.

This extension was our initial goal, which sparked the discussion about language characteristics and design guidelines for mathematical models. When we originally tried to implement this behavior in the monolithic version, we found it extremely hard to pinpoint the lines of code that would need to change. Now, with the modular version, the question was not “Which variables do I have to change?” but “Which influence does a PVC have on physiological component X?”. The discussion shifted from technological considerations to physiological ones, which made the extension possible without much effort. In the original model, several variables and equations would have to be added, making the already complicated system almost unmanageable. The benefits of modularity become even more apparent when moving from the conduction model to the whole SHM.

Since the whole `ModularConduction` model is also encapsulated with a simple interface consisting of an input, an output, the interbeat interval and the timestamp of the last contraction it can seamlessly be integrated into our modular version of the SHM. In fact, switching between the monolithic and modular implementation becomes as simple as changing the type of a variable from `MonolithicConduction` to `ModularConduction`. This is in stark contrast to the original implementation by Seidel in C, where the variables and equations for the conduction system where scattered throughout the code of the whole model. As a welcome side effect, the separation of the model into individual physiological effects also revealed some design flaws in the original. For example, it was not quite clear if the refractory term referred to the refractory behavior of the SA node, the AV node, or the ventricles and in Supplementary Note 1 we discovered that the C implementation introduces a seemingly unphysiological time-dependence in the effective duration of the refractory period.

It has to be said that the components we developed are only reusable within their physiological context. In contrast to, e.g., reaction equations in metabolism models, physiology is not yet standardized enough to have a unifying theory that allows building libraries of components that can be used in multiple tissue or organ models. However, the Physiolibrary can be seen as a first approach in this direction, which also uses Modelica [38].

### Human-readable

All variables, parameters and components in our model have human-readable labels that clearly specify which physiological quantity they represent. The full version of the model code, which can be found in Supplementary Listing 1–27 and at https://github.com/CSchoel/shm-conduction, also contains additional documentation. This is important for the understandability of the model but also for reuse and extension. Reuse requires the identification of possible connection points between variables in different models based on their semantics. Extension could, for example, involve the replacement of one variable or component with a more complex representation, which models the same concept in more detail.

The new model also removed an undocumented technical workaround from the original where the interaction between refractory time, spontaneous beats by the AV node, and the time delay were resolved indirectly: A scheduling system kept track of the next time a beat would be issued, giving precedence to beats that enter the schedule at a later time but would take effect earlier. A diagram of this system can be seen in Supplementary Figure 1. This indirect implementation was hard to understand, because the schedules have no direct physiological equivalent. The AV node, for example, does not signal the sinus node ahead of time to indicate when it will issue the next beat. This systems was therefore replaced by an explicit, more readable version by only considering actual signals and no schedules.

Modelica focuses on human-readability over machine-readability, which enabled us to discuss code-level details in this article and makes it possible to quickly review changes in a version control tool like Git (https://gitscm.com/). This made it easy for us to spot bugs by tagging working versus broken versions and identifying the lines that changed between the working and the broken state. In contrast, editing software for XML-based formats with less focus on human-readability may re-order lines from one version to another without consequence to the overall functioning of the model and without these changes being apparent to the modeler. This can result in long line change messages in popular version control systems such as Git, which complicates locating the individual line that introduced an error.

Another advantage of fully human-readable code can be seen in the experiment setup, which we show in section 4.3. This model contains the code `__OpenModelica_simulationFlags (s = “dassl”)`, which tells the OpenModelica compiler to use the Differential Algebraic System SoLver (DASSL) [89] to solve the equation system. This information can only be interpreted by OpenModelica and not by other tools, which might not support this solver. However, since Modelica is designed to be written by humans directly, researchers who inspect the model can easily find this information without knowing that it is there. In contrast, if the model was written in an XML-based language, researchers would probably not look at the raw code, but load the model in a tool that uses an intermediary language or a graphical user interface to display the model content. It is likely that such a tool would just discard information that it cannot process, making it possible that details like these tool-dependent solver settings might be overlooked in reproduction attempts.

However, our model also has a downside with regard to its human-readability: The visual annotations, which are only helpful within a tool like OpenModelica, do introduce visual clutter when the model is only viewed in text form. To some extent, we could alleviate this issue by separating icon definitions into separate files and base classes and including them via multiple inheritance. Yet still the model `ModularConduction` has to include verbose **annotation()** statements for the placement and connection of components.

### Hybrid

The example model is purely discrete and therefore not hybrid in itself. However, it is important to note that we could use the same language to describe this discrete model that we also used for the rest of the SHM, which is mainly continuous. It was, for example, not needed to explicitly set the derivatives of the discrete variables to zero, which would introduce visual clutter and therefore reduce understandability. While our example cannot directly show the benefits of DAEs and acausality, these features *are* included in the implementation of the SHM that we published previously. One example of this are the acetylcholine kinetics, which use the following connector interface.

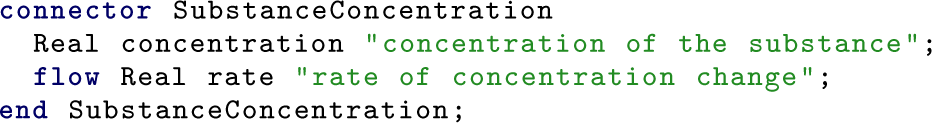

The keyword **flow** indicates that the variable `rate` is subject to a conservation law: At each connection point in the system, the sum of acetylcholine flow from and to all connected components must be zero. In the SHM, the acetylcholine concentration is only determined by a single `NeurotransmitterRelease` component, which is connected to the parasympathetic system, but it is easy to imagine an extension that includes multiple uptake sites. In such an example, the automatic generation of the conservation law by Modelica would allow to separate the effects of all connected components, which only have to declare their individual contribution to the acetylcholine concentration. This both leads to better encapsulation, making the model more understandable, and it facilitates extension since additional components that have an influence on the concentration can be added without changing any of the equations of the existing components.

### Open

Since the model discussed here is written in an open language with opensource tools, readers can easily run simulations themselves. They simply have to download the latest release from the Github repository at https://github.com/CSchoel/shm-conduction, download and install OpenModelica and load the models using the “Load library” option in the “File” menu. This means that our results can be easily reproduced regardless of available licenses or of the user’s operating system and that researchers who might want to reuse the model or components can quickly run simulations to assess the usefulness of the model for their use case.

However, there may still be some barriers. First, Modelica is not yet widely known in systems biology, which makes it likely that researchers will have to become familiar with a new tool and language in order to reproduce our findings. Second, engineers that use Modelica for industrial applications mostly turn towards proprietary solutions like Dymola (https://www.3ds.com/products-services/catia/products/dymola/), which can be more feature-rich than and not fully compatible with OpenModelica.

### Declarative

Even though the example model only contains time-related variables and no other physical quantities, the fact that we introduced proper SI units is still helpful for understanding. For example, it avoids errors, if the components are reused in another model that measures time in milliseconds instead of seconds. Unfortunately, Modelica does not enforce unit definitions or unit consistency checks. However, when these optional unit declarations are used consistently, they help to quickly identify and solve such order of magnitude errors.

In his C implementation, Seidel implemented a fourth order Runge-Kutta method himself, making this the only numerical method available. By using a declarative language, it becomes possible to easily switch between different solvers which can improve numerical accuracy. Not being tied to a specific numerical method also increases interoperability between models and thus reusability.

Another benefit of the declarative specification that becomes apparent in our example is the increase in mathematical soundness and clarity. The C implementation contained some design choices that were convenient for programming, but neither for understanding nor physiological plausibility. For example, the original model mixed variables that represent actual signals and time stamp variables that schedule signals for the future (see Supplementary Figure 1). To comprehend these formulas a context switch from the physiological meaning to the technical representation is required. Another hurdle for understanding the model is the unclear causality. In the SHM, every effect is triggered by the contraction, even if there is no actual signal feedback from the ventricles to the AV node on a physiological level. By separating the model into smaller physiologically meaningful modules with a unified interface, the Modelica compiler automatically hinted at these concerns, e.g., because variables where missing.

One downside of choosing Modelica is that we could not demonstrate the benefit of augmenting a model with semantic information using ontologies. However, the use of the `SIunits` package and the definition of the `InstantSignal` type to indicate Kronecker delta behavior of in- and outputs show how this could be achieved through a type hierarchy: The variables in the model do not only have units, but we also distinguish between the type `Time` for a point in time and `Duration` for a difference between two points in time. Similarly, `InstantSignal` is technically equivalent to the type `Boolean`, but carries additional semantic information about the shape of the signal. In much the same way, one could build large type hierarchies containing all terms of an ontology like the SBO, Chemical Entities of Biological Interest (ChEBI) [90] or Ontology of Physics for Biology (OPB) [91]. Another way to implement ontology support in Modelica would be so called vendor-specific annotations of the form **annotation**`(__VendorName(key=value, …))`, which could be added to components and variables. Since ontology terms are typically identified through Uniform Resource Identifiers (URIs), which are not human-readable, and because there are currently no graphical Modelica tools that support such ontologies, the first approach using the type system seems preferable for now.

### Graphical

The diagram in Figure 5 helps to understand the model at first glance. It can both be used as an entry point for understanding and for communicating the model to a domain expert who is not familiar with mathematical modeling or the language Modelica. The same would not be possible with more detailed SBGN graphs or automatically generated graphs of variable interactions, which would also include internal variables of components and helper variables. At the same time, the diagram is not just a separate image file but it is generated from **annotation()** statements in the individual components themselves. This means that it will remain up to date if components are added or removed or new connectors are included so that other researchers can rely on the accuracy of the diagram if they want to understand, reuse, or reproduce the model. The annotations also allow building more complex models or small test cases using drag and drop in a graphical tool like OpenModelica [72], which can facilitate reuse and extension. For example, the PVC extension required very little changes in the code. Most of the changes could be applied by adding a `RefractoryGate` component and three logical AND and OR gates to the diagram which can be seen in Figure 6.

On the downside, it can be argued that the connection in Figure 5 which points back from the refractory component to the pacemaker component is unintuitive and may be confusing when the model is interpreted physiologically. This can be remedied by introducing another layer of abstraction, which combines the components `Pacemaker` and `RefractoryGate` to a single component `RefractoryPacemaker`. We did not do this in our implementation to keep the model code as simple as possible, but in a larger model such an intermediary component may be advisable.

As of now our discussion was focused on Modelica, but in Section 2.2 it could be seen that there are multiple languages with MoDROGH characteristics. Additionally, our example revealed some shortcomings of Modelica with respect to modeling biological systems. We therefore want to recapitulate which features of the language were especially beneficial for our model design, which features were lacking, and what are the trade-offs that have to be made when switching to another language.

Our Modelica implementation made heavy use of object orientation, including multiple inheritance; it featured discrete variables with human-readable labels; it relied on the graphical representation for creating and communicating the toplevel model structure; it provided minimal interfaces by encapsulating helper variables and defining explicit connectors; and it used the built-in support for SI units. Unfortunately, unit definition are not enforced in Modelica, which means that it is up to the modeler to ensure their reliable use. Modelica also does not support semantic annotation of model components with ontological terms. We showed how this can be achieved with a type hierarchy or with vendor-specific annotations, but still ontology support would have to be added to open-source Modelica tools to be of practical use.

As an alternative, MATLAB with the Simulink environment and the Simscape language is the only other language presented in Section 2.2 that supports full object orientation, declaration of discrete variables, and integrated graphical annotations. In contrast to Modelica, it also enforces unit checks at the interfaces between components. The only technical downside of this language is that Simscape classes do not support multiple inheritance. However, this is probably no issue since we only used multiple inheritance to import the annotation code for icons. Simulink only supports icons as links to separate image files and not as verbose vector graphics code. This has the benefit of not cluttering the code and therefore removing the requirement for multiple inheritance, but it also has the drawback that it is not possible to define a common appearance for all components by inheriting parts of the graphical annotation from the base class. Apart from the technical aspects, the biggest drawback of MATLAB is that it is not open. If we had used MATLAB instead of Modelica, researchers who want to repeat our experiments could still download the code from GitHub, but they would need licenses for Matlab, Simulink, and Simscape to run the simulations.

From the open alternatives, Julia with the Modia package comes closest to the features we used for our example model. It has the drawback of not supporting any graphical representation and still being in an experimental stage. If one instead makes the switch to the more stable DifferentialEquations.jl, the support for object-orientation, declaration of discrete variables, variable labels and encapsulation is lost.

To compensate the shortcomings of Modelica, CellML might be the best fit, since it is the only language from those presented in Section 2.2 that enforces unit definitions. Additionally, it also supports model annotation with terms from an ontology and as an accepted COMBINE standard it is part of an established ecosystem for mathematical modeling in systems biology. With SemGen, there even exists a tool for semantics-based annotation and composition of CellML (and SBML) models [92]. On the downside, CellML does not support full object orientation for composing models. This means that base classes that define an interface need to be imported as *components* of the model, which requires a more verbose syntax. It also does not support the explicit declaration of discrete variables but does support variable reinitialization due to discrete events. To implement a variable like the interbeat interval `d_interbeat`, which stays constant between events, an additional equation would have to be added to the model, which sets the derivative of this variable to zero. This both introduces unnecessary code and makes the model less understandable as there is no clear distinction between discrete and continuous parts apart from the labels assigned to the variables. CellML also follows a different philosophy for graphical annotations, providing basic support for referencing and captioning externally-stored images as part of the documentation for the whole model. Finally, the language is not designed to be written directly by humans and instead relies on the use of appropriate tools to view and edit models.

It becomes clear that there is no single “best” language. Further development is needed to obtain a language that supports all the MoDROGH characteristics to their fullest extent. This development could start with Modelica, leveraging a flexible and industry-proven general-purpose modeling language and extending it to fit the needs of the systems biology community. It could also start with CellML or SBML, which are already proven languages with widespread support in the systems biology community, and which could borrow some software engineering features and standards from languages like Modelica or Simscape. Other approaches and foundations are also possible and it may even make sense to not pursue the “perfect” language at all, but to focus more on interoperability between languages that fit the specialized needs of smaller modeling domains.

In conclusion, using a modeling language that is Modular, Descriptive, human-Readable, Open, Graphical, and Hybrid (MoDROGH) can make models more reproducible, understandable, reusable, and extensible. Because there is no single best language, modelers have to decide which features are most important for them and which trade-offs they are willing to make. They should be aware of the beneficial characteristics of the language and use them consistently as we described in our guidelines and showed in our modular example model of the cardiac conduction system. The situation that a model needs to be dissected, modified and extended to be used in a different context is common in multi-scale, multi-level, and multi-class models and therefore it is likely that our findings translate to large areas of systems biology. Mathematical modeling in systems biology has become an engineering challenge that requires engineering solutions. Models should no longer be implemented with only a single purpose in mind, but as reliable parts of larger systems. We hope that this article can spark a discussion in the community to put more emphasis on these engineering aspects of mathematical modeling in the development, selection, and application of modeling languages.

## 4 Methods

### 4.1 Material

We used Mo|E version 0.6.3 [93] to write the code of our models and OpenModelica version 1.13.0 [72] as well as Inkscape version 0.91 (https://inkscape.org/) to add the component icons. OpenModelica was also used for all simulations.

In the following we will show and explain our Modelica code for the models and simulations. To keep it short, we do not show the code of the original monolithic version and of the graphical annotations. We also do not include most of the documentation strings, which are present in the full version. They can be found in Supplementary Listing 1–27 and at https://github.com/CSchoel/shm-conduction. Please also note that this article was previously published as a preprint [79].

### 4.2 Modular conduction model

The fundamental part of the modular model of the human cardiac conduction system is the interface component `UnidirectionalConductionComponent`, which serves as a base class for all other components. It has already been shown in Section 2.3. It defines the input and output connectors `inp` and `outp`, which are Booleans that are wrapped in a custom type `InstantSignal` to indicate that they behave as a sum of Kronecker deltas, meaning that they are only true for the exact instants in time when events occur:

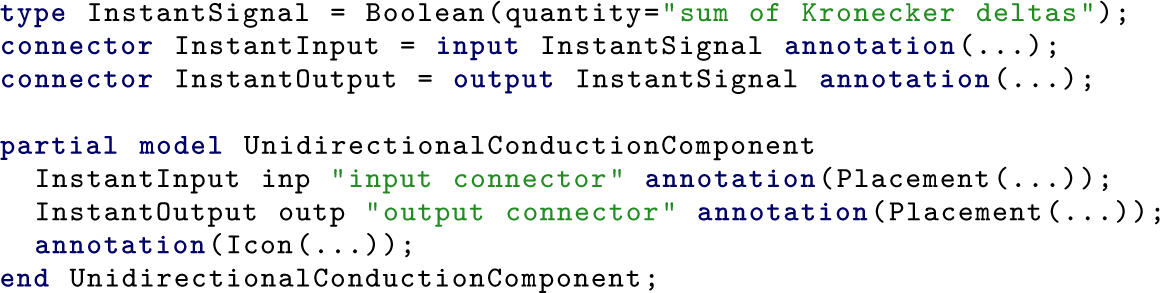

The keyword **connector** designates the types `InstantInput` and `InstantOutput` as part of the interface of a class and allows the assignment of a basic icon representation in the form of an **annotation()** statement. The content of these annotation statements can be quite verbose, which is why we only show them in the full code in Supplementary Listing 1–27 as well as on GitHub. The model `UnidirectionalConductionComponent` is defined as **partial** to designate that it is not designed to be used as a finished component but has to be extended in some way—in this case by defining the relationship between the input and the output.

The `RefractoryGate` has already been shown in Section 2.3. The component passes on its input signal as output signal, but only when the elapsed time since the last signal left the component is larger than the refractory period:

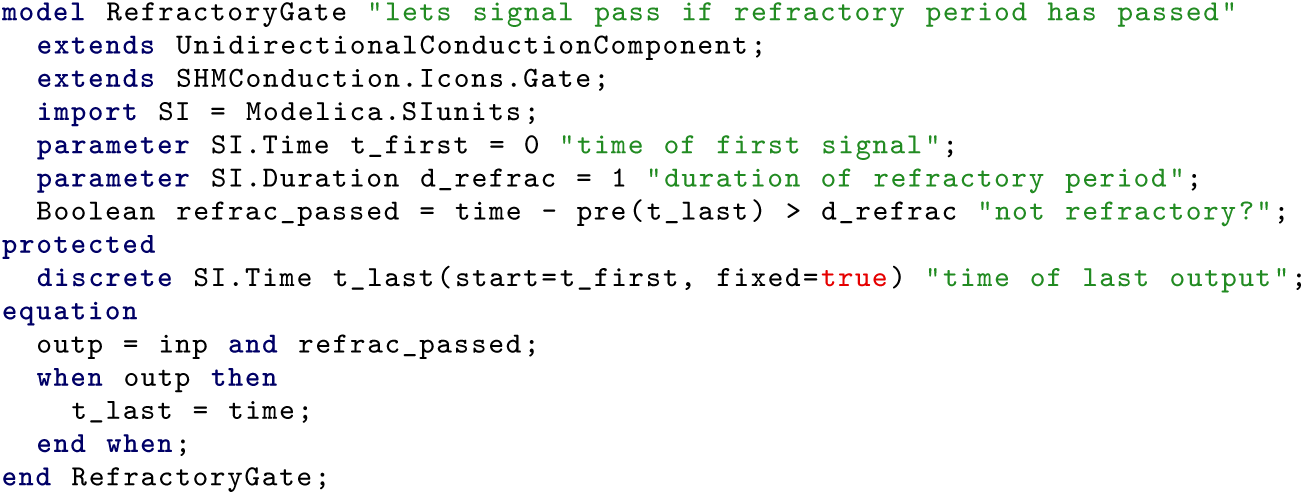

The function `pre()` is used here to denote the value right before an event instead of the value right after the event. The **when** statement describes an event which can define states of discrete variables and can reinitialize continuous variables. This model also introduces a **protected** section which contains variables and parameters that should not be visible from the outside.

The `Pacemaker` model propagates incoming signals, but also adds its own signal if there was no input for a certain period of time. Additionally, this component, too, has to ignore incoming signals during the refractory period. This can be implemented by decoupling the reset of the pacemaker timer from the output of the component and instead treating the reset signal as an additional input. It is assumed that this reset signal is only triggered if the signal passes not only the pacemaker but also the subsequent `RefractoryGate` component. The pacemaker component itself still resets when a spontaneous output signal is generated to maintain the invariant that the output signal will not be true for a prolonged period of time:

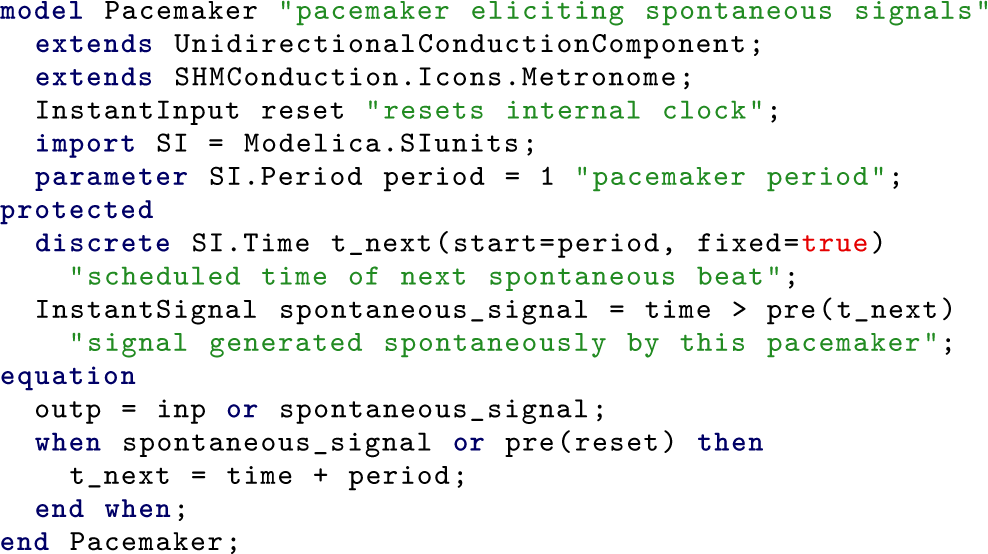

The `ConductionDelay` model puts incoming signals on hold and releases them after a certain time has passed. Physiologically the duration of the delay for each signal depends on the time that has passed between the last signal leaving the component and the current input signal. The original model silently assumed that there will never be a second input signal while a signal is put on hold. Therefore, this assumption is kept, but made more explicit by using the helper variable `delay_passed` in the **when** condition:

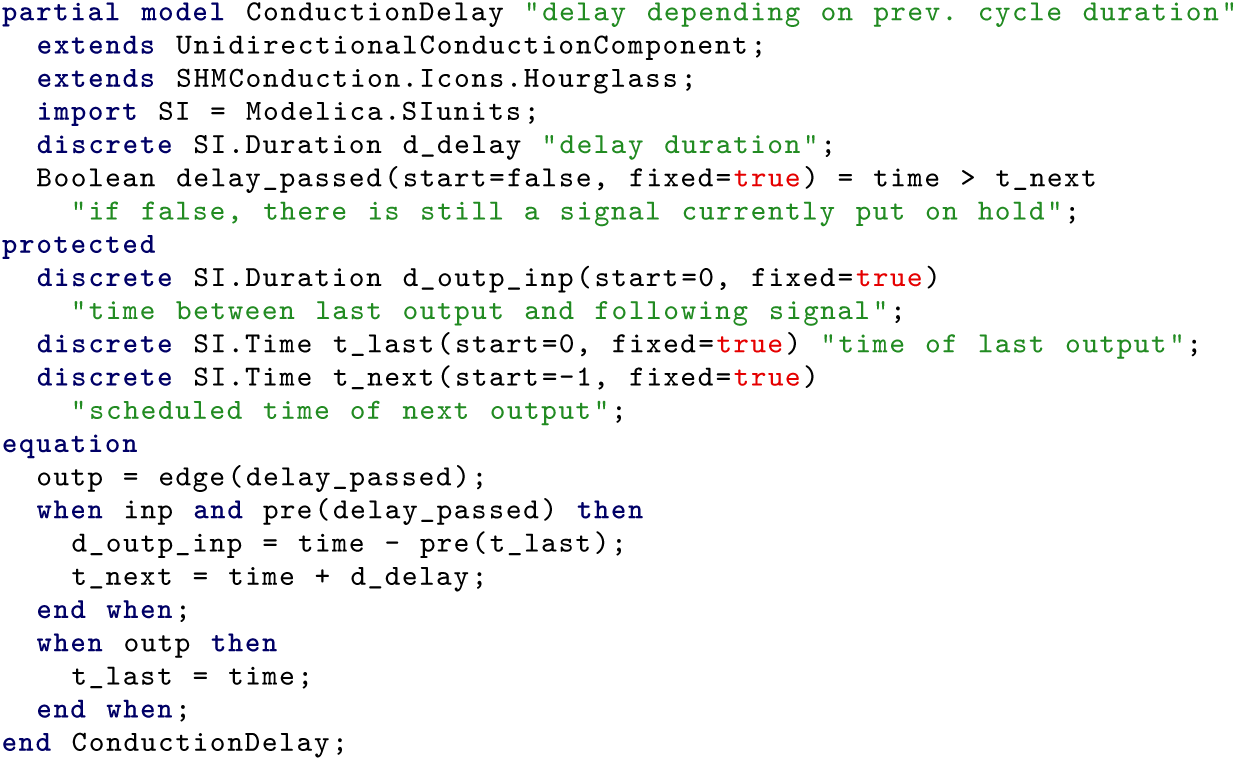

Modelica does already have support for explicit delays, but this feature is tailored towards continuous variables. Therefore, we use a scheduling solution with the variable `t_next`, which indicates the time when the next signal should leave the component. This is similar to the approach in the original C implementation of the SHM, but here this scheduling system is encapsulated in a single component and the respective helper variables are defined in a **protected** environment so that they do not show up in the simulation output.

Note that this is again only a **partial** model, which does not specify the behavior of the variable `d_delay`. This allows the separation of the general delay logic from the physiological equation for the AV node which is modeled in the `AVConductionDelay`:

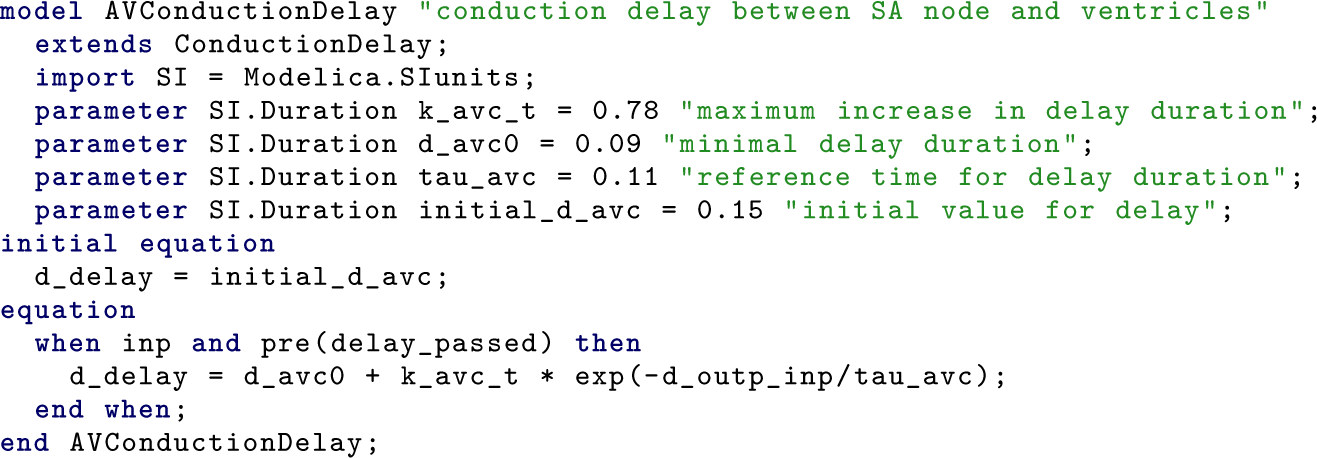

Currently, this separation is only performed to increase readability and to not further complicate the already complex `ConductionDelay` component. Additionally, if the delay was split into two components, as discussed in Section 2.3, the second delay component could also inherit the base equations from `ConductionDelay`, which would avoid code duplication.

Finally, the model `ModularConduction` combines the aforementioned components using **connect()** equations to connect the input and output variables. These equations are represented as lines in the graphical representation which are again defined in **annotation()** statements:

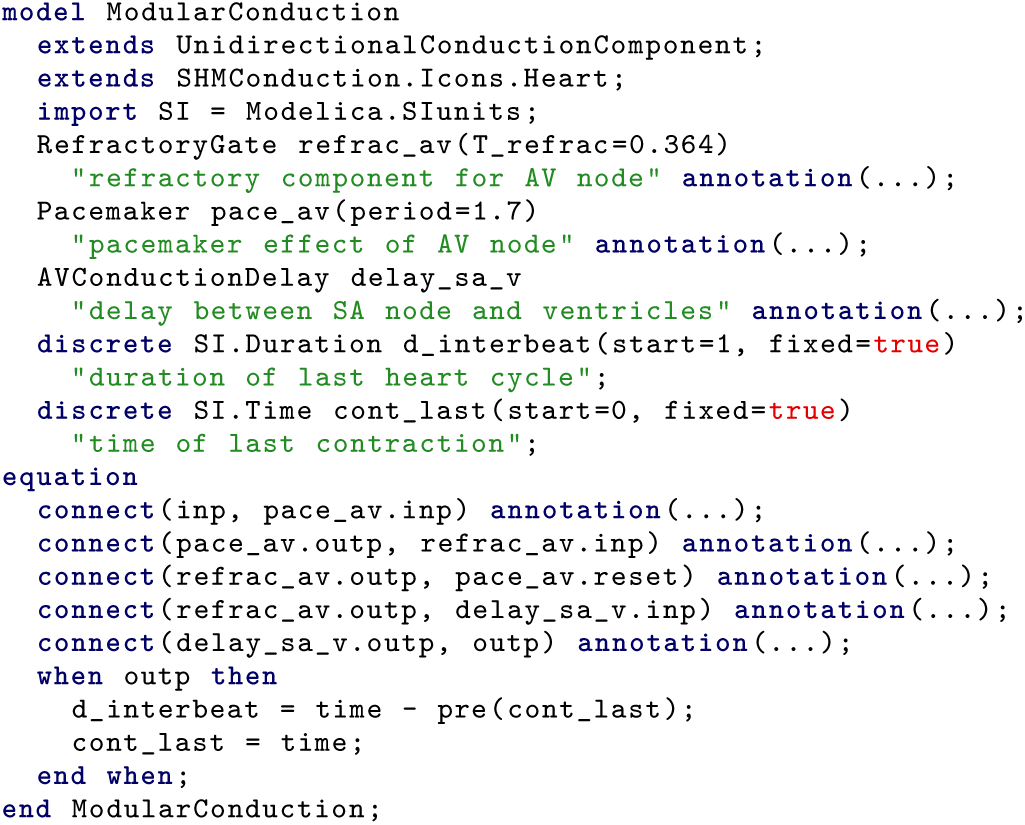

As already mentioned in Section 2.3, this model also shows how parameters like `d_refrac` and `period` can be adjusted when the components are imported. Note also that the model `ModularConduction` is again an `UnidirectionalConductionComponent` and can therefore be used as a component in a larger model such as the SHM.

### 4.3 Modular contraction experiment setup

Simulation experiments can also be defined directly in Modelica syntax. The following code was used to produce Supplementary Figure 2:

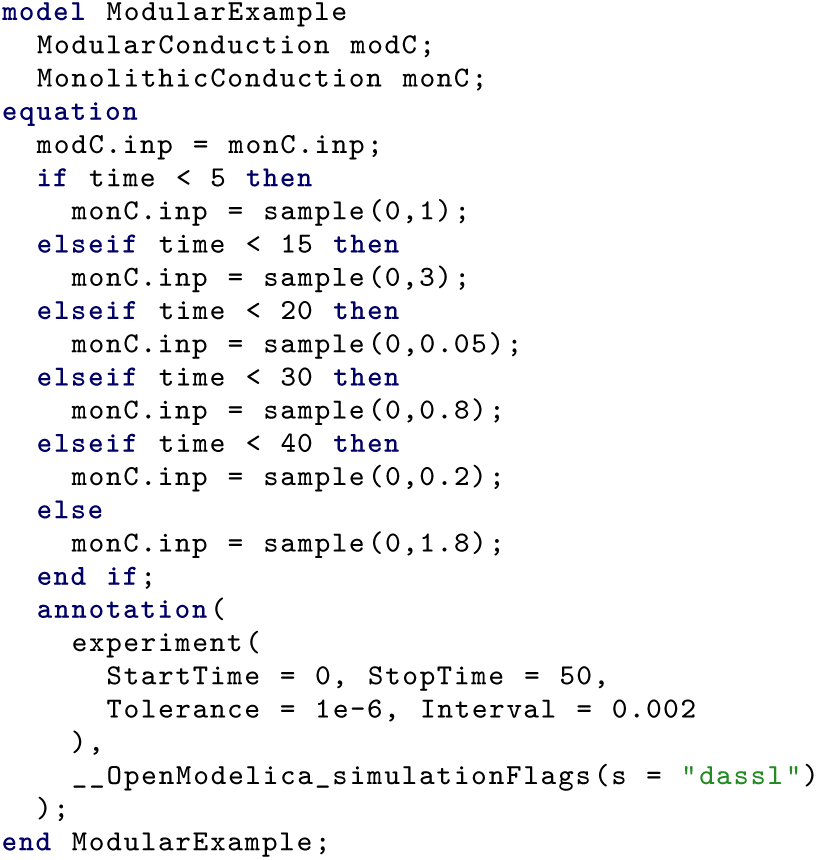

Here, the built-in function `sample(start, interval)` is used to issue signals from the SA node at a precise interval. The interval length is switched every five to ten seconds using an `if` statement. In addition to the experiment setup, the experiment protocol is also given by the `experiment()` annotation, which defines the start and stop times of the interval, the requested step size for the output and the tolerance used in the solver settings. The vendor-specific annotation `__OpenModelica_simulationFlags` is used to define the DASSL [89] as the default solver. For Supplementary Figure 2, the variables `monC.d_interbeat` and `modC.d_interbeat` were plotted against simulation time.

## Acknowledgements

We thank our four anonymous reviewers for their much valued input which greatly improved the quality of this article. C. Schölzel would like to thank Denis Noble, Peter Hunter, James Bassingthwaighte, Maxwell Neal and Herbert Sauro for insightful discussions about the IUPS and NSR Physiome projects, CellML and present and future challenges for systems biology. We also thank Alexander Goesmann for his advice regarding the focus of the article and Jochen Blom, Björn Pfarr and Annina Hofferberth for proofreading the manuscript.

## Author contributions

C.S., V.B. and A.D. conceived the project, C.S. implemented the models and performed the experiments, V.B. and G.E. provided physiological consultation and criticism, C.S. drafted the manuscript and V.B., G.E. and A.D. revised it.

## Data availability

The full code of the models and experiments used in this article can be found on GitHub, https://github.com/CSchoel/shm-conduction. The data for figures in the data supplement can be generated from this model code.

## Footnotes

1 Note that although the journal article [11] was published one year after the PhD thesis [10], the PhD thesis actually contains the latest version of the model with many small improvements.

2 Seidel probably meant to include the refractory behavior of the ventricles and not the SA node. The actual implementation, however, checks the refractory state *before* the delay between SA node and ventricles is applied.

## References

1. Hodgkin, A. L. & Huxley, A. F. A Quantitative Description of Membrane Current and Its Application to Conduction and Excitation in Nerve. The Journal of Physiology 117, 500–544 (1952).

2. Bardini, R., Politano, G., Benso, A. & Di Carlo, S. Multi-Level and Hybrid Modelling Approaches for Systems Biology. Computational and Structural Biotechnology Journal 15, 396–402 (2017).

3. Uhrmacher, A. M., Degenring, D. & Zeigler, B. Discrete Event Multi-Level Models for Systems Biology in Transactions on Computational Systems Biology I (ed Priami, C.) 66–89 (Springer, Berlin; Heidelberg, 2005).

4. Dada, J. O. & Mendes, P. Multi-Scale Modelling and Simulation in Systems Biology. Integrative Biology 3, 86 (2011).

5. Yu, J. S. & Bagheri, N. Multi-Class and Multi-Scale Models of Complex Biological Phenomena. Current Opinion in Biotechnology 39, 167– 173 (2016).

6. Waltemath, D. & Wolkenhauer, O. How Modeling Standards, Software, and Initiatives Support Reproducibility in Systems Biology and Systems Medicine. IEEE Transactions on Biomedical Engineering 63, 1999–2006 (2016).

7. Medley, J. K., Goldberg, A. P. & Karr, J. R. Guidelines for Reproducibly Building and Simulating Systems Biology Models. IEEE transactions on bio-medical engineering 63, 2015–2020 (2016).

8. Tiwari, K. et al. Reproducibility in Systems Biology Modelling preprint 10.1101/2020.08.07.239855 (bioRxiv, 2020).

9. Topalidou, M., Leblois, A., Boraud, T. & Rougier, N. P. A Long Journey into Reproducible Computational Neuroscience. Frontiers in Computational Neuroscience 9, 1–2 (2015).

10. Seidel, H. Nonlinear Dynamics of Physiological Rhythms PhD thesis (Technische Universität Berlin, Berlin, Germany, 1997).

11. Seidel, H. & Herzel, H. Bifurcations in a Nonlinear Model of the Baroreceptor-Cardiac Reflex. Physica D: Nonlinear Phenomena 115, 145–160 (1998).

12. Schölzel, C., Goesmann, A., Ernst, G. & Dominik, A. Modeling Biology in Modelica: The Human Baroreflex in Proceedings of the 11th International Modelica Conference (Versailles, France, 2015), 367–376.

13. Sarma, G. P. et al. Unit Testing, Model Validation, and Biological Simulation. F1000Research 5, 1946 (2016).

14. Miskovic, L., Tokic, M., Fengos, G. & Hatzimanikatis, V. Rites of Passage: Requirements and Standards for Building Kinetic Models of Metabolic Phenotypes. Current Opinion in Biotechnology 36, 146–153 (2015).

15. Hicks, J. L., Uchida, T. K., Seth, A., Rajagopal, A. & Delp, S. L. Is My Model Good Enough? Best Practices for Verification and Validation of Musculoskeletal Models and Simulations of Movement. Journal of Biomechanical Engineering 137, 020905 (2015).

16. Grimm, V. & Railsback, S. F. Pattern-Oriented Modelling: A ‘multi-Scope’ for Predictive Systems Ecology. Philosophical Transactions of the Royal Society B: Biological Sciences 367, 298–310 (2012).

17. Zhao, P., Rowland, M. & Huang, S.-M. Best Practice in the Use of Physiologically Based Pharmacokinetic Modeling and Simulation to Address Clinical Pharmacology Regulatory Questions. Clinical Pharmacology & Therapeutics 92, 17–20 (2012).

18. Smith, N. P., Crampin, E. J., Niederer, S. A., Bassingthwaighte, J. B. & Beard, D. A. Computational Biology of Cardiac Myocytes: Proposed Standards for the Physiome. Journal of Experimental Biology 210, 1576– 1583 (2007).

19. Goldberg, A. P. et al. Emerging Whole-Cell Modeling Principles and Methods. Current Opinion in Biotechnology 51, 97–102 (2018).

20. Bartocci, E. & Lió, P. Computational Modeling, Formal Analysis, and Tools for Systems Biology. PLOS Computational Biology 12, e1004591 (2016).

21. Walpole, J., Papin, J. A. & Peirce, S. M. Multiscale Computational Models of Complex Biological Systems. Annual Review of Biomedical Engineering 15, 137–154 (2013).

22. Hucka, M. et al. Promoting Coordinated Development of Community-Based Information Standards for Modeling in Biology: The COMBINE Initiative. Frontiers in Bioengineering and Biotechnology 3 (2015).

23. Wolstencroft, K. et al. SEEK: A Systems Biology Data and Model Management Platform. BMC Systems Biology 9 (2015).

24. Cooling, M. T., Hunter, P. & Crampin, E. J. Modelling Biological Modularity with CellML. IET Systems Biology 2, 73–79 (2008).

25. Neal, M. L. et al. A Reappraisal of How to Build Modular, Reusable Models of Biological Systems. PLOS Computational Biology 10, e1003849 (2014).

26. Waltemath, D. et al. The First 10 Years of the International Coordination Network for Standards in Systems and Synthetic Biology (COMBINE). Journal of Integrative Bioinformatics 17, 20200005 (2020).

27. Malik-Sheriff, R. S. et al. BioModels—15 Years of Sharing Computational Models in Life Science. Nucleic Acids Research 48, D407–D415 (2019).

28. Cooling, M. T. et al. Standard Virtual Biological Parts: A Repository of Modular Modeling Components for Synthetic Biology. Bioinformatics 26, 925–931 (2010).

29. Clerx, M., Collins, P., de Lange, E. & Volders, P. G. Myokit: A Simple Interface to Cardiac Cellular Electrophysiology. Progress in Biophysics and Molecular Biology 120, 100–114 (2016).

30. Mulugeta, L. et al. Credibility, Replicability, and Reproducibility in Simulation for Biomedicine and Clinical Applications in Neuroscience. Frontiers in Neuroinformatics 12 (2018).

31. Olivier, B. G., Rohwer, J. M. & Hofmeyr, J.-H. S. Modelling Cellular Systems with PySCeS. Bioinformatics 21, 560–561 (2005).

32. Clewley, R. Hybrid Models and Biological Model Reduction with PyDSTool. PLoS Computational Biology 8, e1002628 (2012).

33. Lopez, C. F., Muhlich, J. L., Bachman, J. A. & Sorger, P. K. Programming Biological Models in Python Using PySB. Molecular Systems Biology 9, 646 (2013).

34. Choi, K. et al. Tellurium: An Extensible Python-Based Modeling Environment for Systems and Synthetic Biology. Biosystems 171, 74–79 (2018).

35. Smith, L. P., Bergmann, F. T., Chandran, D. & Sauro, H. M. Antimony: A Modular Model Definition Language. Bioinformatics 25, 2452–2454 (2009).

36. Bezanson, J., Edelman, A., Karpinski, S. & Shah, V. B. Julia: A Fresh Approach to Numerical Computing. SIAM Review 59, 65–98 (2017).

37. Mattsson, S. E. & Elmqvist, H. Modelica – An International Effort to Design the next Generation Modeling Language in 7th IFAC Symposium on Computer Aided Control Systems Design, CACSD’97 30 (Gent, Belgium, 1997), 151–155.

38. Mateják, M. et al. Physiolibrary - Modelica Library for Physiology in Proceedings of the 10th International Modelica Conference 96 (Lund, Sweden, 2014), 499–505.

39. Maggioli, F., Mancini, T. & Tronci, E. SBML2Modelica: Integrating Biochemical Models within Open-Standard Simulation Ecosystems. Bioinformatics 36, 2165–2172 (2019).

40. Hellerstein, J. L., Gu, S., Choi, K. & Sauro, H. M. Recent Advances in Biomedical Simulations: A Manifesto for Model Engineering. F1000Research 8, 261 (2019).

41. Blochwitz, T. et al. The Functional Mockup Interface for Tool Independent Exchange of Simulation Models in Proceedings of the 8th International Modelica Conference (Dresden, Germany, 2011), 105–114.

42. Blochwitz, T. et al. Functional Mockup Interface 2.0: The Standard for Tool Independent Exchange of Simulation Models in Proceedings of the 9th International Modelica Conference (Munich, Germany, 2012), 173–184.

43. Zhu, X.-G. et al. Plants *in Silico*: Why, Why Now and What?-An Integrative Platform for Plant Systems Biology Research. Plant, Cell & Environment 39, 1049–1057 (2016).

44. Mirschel, S., Steinmetz, K., Rempel, M., Ginkel, M. & Gilles, E. D. ProMoT: Modular Modeling for Systems Biology. Bioinformatics 25, 687–689 (2009).

45. Kell, D. The Virtual Human: Towards a Global Systems Biology of Multi-scale, Distributed Biochemical Network Models. IUBMB Life 59, 689–695 (2007).

46. Hasenauer, J., Jagiella, N., Hross, S. & Theis, F. J. Data-Driven Modelling of Biological Multi-Scale Processes. Journal of Coupled Systems and Multi-scale Dynamics 3, 101–121 (2015).

47. Garny, A. & Hunter, P. J. OpenCOR: A Modular and Interoperable Approach to Computational Biology. Frontiers in Physiology 6, 26 (2015).

48. Oliveira, A., Kohwalter, T., Kalinowski, M., Murta, L. & Braganholo, V. XChange: A Semantic Diff Approach for XML Documents. Information Systems 94, 101610 (2020).

49. Drager, A. et al. SBML2LaTeX: Conversion of SBML Files into Human-Readable Reports. Bioinformatics 25, 1455–1456 (2009).

50. Lincoln, P. & Tiwari, A. Symbolic Systems Biology: Hybrid Modeling and Analysis of Biological Networks in Hybrid Systems: Computation and Control (eds Alur, R. & Pappas, G. J.) Lecture Notes in Computer Science 2993, 660–672 (Springer, Berlin; Heidelberg, 2004).

51. Bortolussi, L. & Policriti, A. Hybrid Systems and Biology in Formal Methods for Computational Systems Biology (eds Bernardo, M., Degano, P. & Zavattaro, G.) 424–448 (Springer, Berlin; Heidelberg, 2008).

52. Rejniak, K. A. & Anderson, A. R. A. Hybrid Models of Tumor Growth. Wiley Interdisciplinary Reviews: Systems Biology and Medicine 3, 115–125 (2011).

53. Yu, T. et al. The Physiome Model Repository 2. Bioinformatics 27, 743– 744 (2011).

54. Bassingthwaighte, J. B. Strategies for the Physiome Project. Annals of Biomedical Engineering 28, 1043–1058 (2000).

55. Holzhütter, H.-G., Drasdo, D., Preusser, T., Lippert, J. & Henney, A. M. The Virtual Liver: A Multidisciplinary, Multilevel Challenge for Systems Biology. Wiley Interdisciplinary Reviews: Systems Biology and Medicine 4, 221–235 (2012).

56. Zhu, H., Huang, S. & Dhar, P. The next Step in Systems Biology: Simulating the Temporospatial Dynamics of Molecular Network. BioEssays 26, 68–72 (2004).

57. Loew, L. M. & Schaff, J. C. The Virtual Cell: A Software Environment for Computational Cell Biology. Trends in Biotechnology 19, 401–406 (2001).

58. Butterworth, E., Jardine, B. E., Raymond, G. M., Neal, M. L. & Bassingthwaighte, J. B. JSim, an Open-Source Modeling System for Data Analysis. F1000Research 2, 288 (2013).

59. Yan, K. & Cui, W. Visualizing the Uncertainty Induced by Graph Layout Algorithms in 2017 IEEE Pacific Visualization Symposium (PacificVis) (Seoul, South Korea, 2017), 200–209.

60. Kerren, A. & Schreiber, F. Network Visualization for Integrative Bioinformatics in Approaches in Integrative Bioinformatics (eds Chen, M. & Hofestädt, R.) 173–202 (Springer, Berlin; Heidelberg, 2014).

61. Le Novère, N. et al. The Systems Biology Graphical Notation. Nature Biotechnology 27, 735–741 (2009).

62. Gonçalves, E., Iersel, M. & Saez-Rodriguez, J. CySBGN: A Cytoscape Plug-in to Integrate SBGN Maps. BMC Bioinformatics 14, 17 (2013).

63. Gauges, R., Rost, U., Sahle, S., Wengler, K. & Bergmann, F. T. The Systems Biology Markup Language (SBML) Level 3 Package: Layout, Version 1 Core. Journal of Integrative Bioinformatics 12, 550–602 (2015).

64. Bergmann, F. T., Keating, S. M., Gauges, R., Sahle, S. & Wengler, K. SBML Level 3 Package: Render, Version 1, Release 1. Journal of Integrative Bioinformatics 15 (2018).

65. Alves, R., Antunes, F. & Salvador, A. Tools for Kinetic Modeling of Biochemical Networks. Nature Biotechnology 24, 667–672 (2006).

66. Mangourova, V., Ringwood, J. & Van Vliet, B. Graphical Simulation Environments for Modelling and Simulation of Integrative Physiology. Computer Methods and Programs in Biomedicine 102, 295–304 (2011).

67. Keating, S. M. et al. SBML Level 3: An Extensible Format for the Exchange and Reuse of Biological Models. Molecular Systems Biology 16, e9110 (2020).

68. Bornstein, B. J., Keating, S. M., Jouraku, A. & Hucka, M. LibSBML: An API Library for SBML. Bioinformatics 24, 880–881 (2008).

69. Cuellar, A. A. et al. An Overview of CellML 1.1, a Biological Model Description Language. SIMULATION 79, 740–747 (2003).

70. Nickerson, D. & Buist, M. Practical Application of CellML 1.1: The Integration of New Mechanisms into a Human Ventricular Myocyte Model. Progress in Biophysics and Molecular Biology 98, 38–51 (2008).

71. Margolis, B. W. L. SimuPy: A Python Framework for Modeling and Simulating Dynamical Systems. The Journal of Open Source Software 2, 396 (2017).

72. Fritzson, P. et al. The OpenModelica Modeling, Simulation, *and Development Environ*ment in Proceedings of the 46th Conference on Simulation and Modelling of the Scandinavian Simulation Society (Trondheim, Norway, 2005).

73. Åkesson, J. R., Gäfvert, M. & Tummescheit, H. JModelica—An Open Source Platform for Optimization of Modelica Models in Proceedings of the 6th Vienna International Conference on Mathematical Modelling 34 (Vienna, Austria, 2009).

74. Elmqvist, H., Henningsson, T. & Otter, M. Systems Modeling and Programming in a Unified Environment Based on Julia in ISoLA 2016: Leveraging Applications of Formal Methods, Verification and Validation: Discussion, Dissemination, Applications 9953 (Corfu, Greece, 2016), 198–217.

75. Rackauckas, C. & Nie, Q. DifferentialEquations.Jl – A Performant and Feature-Rich Ecosystem for Solving Differential Equations in Julia. Journal of Open Research Software 5, 15 (2017).

76. Seidel, H. & Herzel, H. Modelling Heart Rate Variability Due to Respiration and Baroreflex in Modelling the Dynamics of Biological Systems (eds Mosekilde, E. & Mouritsen, O. G.) Springer Series in Synergetics 65, 205–229 (Springer, Berlin; Heidelberg, 1995).

77. Kotani, K., Struzik, Z., Takamasu, K., Stanley, H. & Yamamoto, Y. Model for Complex Heart Rate Dynamics in Health and Diseases. Physical Review E 72, 041904 (2005).

78. Kotani, K., Takamasu, K., Ashkenazy, Y., Stanley, H. & Yamamoto, Y. Model for Cardiorespiratory Synchronization in Humans. Physical Review E 65, 051923 (2002).

79. Schölzel, C. Modelica Implementation of the Seidel-Herzel Model of the Human Baroreflex version 1.6. Zenodo, 2020. https://zenodo.org/record/4110400 (2020).

80. Duggento, A., Toschi, N. & Guerrisi, M. Modeling of Human Baroreflex: Considerations on the Seidel–Herzel Model. Fluctuation and Noise Letters 11, 1240017 (2012).

81. Tiller, M. Modelica by Example https://mbe.modelica.university/ (2020) (Michael Tiller, 2020).

82. Schölzel, C., Blesius, V., Ernst, G. & Dominik, A. An Understandable, Extensible, and Reusable Implementation of the Hodgkin-Huxley Equations Using Modelica. Frontiers in Physiology 11, 583203 (2020).

83. Karr, J. R. et al. A Whole-Cell Computational Model Predicts Phenotype from Genotype. Cell 150, 389–401 (2012).

84. Freeman, E., Robson, E., Bates, B. & Sierra, K. Head First Design Patterns 638 pp. (O’Reilly, Sebastopol, CA, 2004).

85. Kofránek, J., Rusz, J. & Matoušek, S. Guyton’s Diagram Brought to Life - From Graphic Chart to Simulation Model for Teaching Physiology in Technical Computing Prague 2007: 15th Annual Conference Proceedings (Prague, Czech Republic, 2007), 1–13.

86. Briese, L. E., Klöckner, A. & Reiner, M. The DLR Environment Library for Multi-Disciplinary Aerospace Applications in Proceedings of the 12th International Modelica Conference (Prague, Czech Republic, 2017), 929–938.

87. Casella, F., Bartolini, A., Pasquini, S. & Bonuglia, L. Object-Oriented Modelling and Simulation of Large-Scale Electrical Power Systems Using Modelica: A First Feasibility Study in Proceedings of the IECON 2016 - 42nd Annual Conference of the IEEE Industrial Electronics Society (Florence, Italy, 2016), 6298–6304.

88. Sweller, J. Cognitive Load Theory in Advances in Cognitive Load Theory: Rethinking Teaching (eds Tindall-Ford, S., Agostinho, S. & Sweller, J.) 1st ed., 1–11 (Routledge, Abingdon, England, 2019).

89. Petzold, L. R. Description of DASSL: A Differential/Algebraic System Solver Sandia Report SAND82-8637 (Sandia National Laboratories, Albuquerque, New Mexico; Livermore, California, 1982).

90. Hastings, J. et al. ChEBI in 2016: Improved Services and an Expanding Collection of Metabolites. Nucleic Acids Research 44, D1214–D1219 (2016).

91. Cook, D. L., Bookstein, F. L. & Gennari, J. H. Physical Properties of Biological Entities: An Introduction to the Ontology of Physics for Biology. PLoS ONE 6, e28708 (2011).

92. Neal, M. L. et al. Semantics-Based Composition of Integrated Cardiomyocyte Models Motivated by Real-World Use Cases. PLOS ONE 10, e0145621 (2015).

93. Justus, N., Schölzel, C., Dominik, A. & Letschert, T. Mo|E – A Communication Service between Modelica Compilers and Text Editors in Proceedings of the 12th International Modelica Conference (Prague, Czech Republic, 2017), 815–822.

