## Supplementary material for "Characteristics of mathematical modeling languages that facilitate model reuse in systems biology: A software engineering perspective": Data Supplement

February 12, 2021

### 1 Supplementary Note: Accuracy of the modular model with respect to original

#### 1.1 Result: The modular model behaves similar to the original version

Although the modular version of the conduction model presented in Section 2.3 covers the same physiological effects as the original version, the implementation differs not only in structure but also in the mathematical representation. The original model used the elapsed time since the last contraction as a reference for the refractory period and the pacemaker effect of the atrioventricular node (AV node) instead of the elapsed time since the last sinus signal was received. This means that the duration of the refractory period needs to be adjusted. In the old model, the performed check was whether $t - (t_{\text{sinus}} + T_{\text{delay}}) < T_{\text{refrac}}$ where $t$ is the current time stamp, $t_{\text{sinus}}$ is the time stamp of the last sinus signal, $T_{\text{delay}}$ is the duration of the delay and $T_{\text{refrac}}$ is the refractory period. Since the check is now whether $t - t_{\text{sinus}} < T'_{\text{refrac}}$ one can deduce that if the same behavior is desired, $T'_{\text{refrac}}$ should equal $T_{\text{refrac}} + T_{\text{delay}}$, that is $T_{\text{refrac}}$ must be increased by the average delay duration. The pacemaker component does not have to be changed at all, because, although the pacemaker signal is delayed, the pacemaker clock is also started earlier. Effectively the delay duration is added and then subtracted from the resulting time stamp of the next contraction.

Supplementary Figure 2 shows a comparison of the resulting interbeat intervals (i.e. the time passed between two contractions) for both models with input of varying frequency. For the most part both versions behave identically. Only when the sinus cycle length drops below the refractory period one can see a difference in the plots. In these areas the interbeat intervals of the modular model fluctuate with a higher amplitude and lower frequency compared to the original Seidel-Herzel model (SHM).

#### 1.2 Discussion

The simulation of the modular model shows a similar but not identical behavior with respect to the original version. Increased amplitude and lower frequency of fluctuations during very fast sinus rhythms are explained by the use of the average delay duration to adjust the refractory period of the AV node in the modular model. In the original model the delay duration $T_{\text{delay}}$ varies over time. As shown in Section 1.1, this variable also plays a role in the check for the refractory period, making the effective duration of the refractory period itself time-dependent. It is not clear if this behavior is intentional in the SHM or if it is a side effect of other design choices. If this behavior is desired, it can be emulated in the modular version by making the variable `d_refrac` in the component `RefractoryGate` time dependent much like the duration in `ConductionDelay`. Physiologically the refractory period indeed changes when the sinus frequency is increased, but the effect is a decrease instead of an increase as in the monolithic model [1]. It can therefore be said that the modular design both helps to identify plausibility issues and can be more easily adapted to the biological reality.

### 2 Supplementary Note: Extension of the cardiac conduction system with a trigger for PVC

#### 2.1 Result: The new model structure facilitates the extension with a trigger for premature ventricular contractions

To show that the modular structure and the use of a Modular, Descriptive, human-Readable, Open, Graphical, and Hybrid (MoDROGH) language facilitate reuse and extension we added a trigger for premature ventricular contractions (PVCs) to the model. When we tried to do add this extension to the monolithic model, we struggled to identify which equations would have to change and where exactly the effect of a PVC should be incorporated. In the modular model, it is now possible to separately address the effect of a PVC on each individual component.

PVCs arise if some part of the ventricular tissue generates a signal without stimulation from the AV node. This can happen in a healthy individual, but the heart rate response to such an ectopic beat can be used as a risk indicator during pathological conditions [2].

An ectopic beat in the ventricles leads to a stimulation of the AV node that travels back upwards to the sinoatrial node (SA node) either cancelling out an oncoming downward signal or (in rare cases) resetting the clock of the SA node. Therefore, the *correct* way to model a PVC would be to include components that are bidirectional.

To keep it simple, the unidirectional components are used and it is assumed that a PVC will always reset the pacemaker and refractory time of the AV node but never reach the SA node. We modeled this by extending the components `RefractoryGate` and `ConductionDelay` with a "reset" input similar to the one that already exists for the `Pacemaker` component.

With these changes, there could still be two beats arbitrarily close to each other when a PVC is triggered right after a normal beat. Therefore, we also modeled the refractory behavior of the ventricles themselves. The only change needed for this was the addition of a second instance of the already existing `RefractoryGate` component that receives input from the PVC trigger and the delay component (combined with a logical OR). The output of this additional component was used to ensure that the reset of the AV node would only happen if the PVC actually did trigger a contraction. This was achieved by an additional logical AND gate with input from the PVC signal and the contraction output. A graphical representation of the resulting model can be seen in Figure 2 in the main article.

#### 2.2 Result: The PVC extension shows plausible results

The behavior of the resulting model is shown in Supplementary Figure 3. For a normal sinus rhythm of 75 bpm the model behaves as expected. When a PVC happens while a beat is delayed, it replaces the normal beat, leading to one interbeat interval that is shorter than normal followed by one interval that is larger than normal by the same magnitude. The same behavior can be observed for a PVC happening directly before a sinus signal is issued. A PVC that follows a normal beat within the ventricular refractory period is completely ignored and a PVC right between two normal beats leads to two interbeat intervals that are shorter than normal since all three beats (two normal, one ectopic) lead to a contraction.

To test the behavior of the AV node in the presence of ectopic beats, the experiment was repeated without any sinus signal. Here, the behavior is the same for PVCs while an AV signal is delayed, during the ventricular refractory period and directly before an AV signal. A PVC right between two normal beats does not result in two reduced interbeat intervals but in one reduced and one increased interval. This is due to the reset of the pacemaker clock that will issue the next signal after the pacemaker period has passed. This signal has to travel through the delay component, which increases the interval duration.

#### 2.3 Discussion

The PVC model also behaves as expected. In the simulation with a sinus frequency of 75 bpm, the stimulations very close to a sinus signal effectively replace that signal, leading to a short coupling interval and a long compensatory pause. Stimulations during the refractory period are correctly ignored and a stimulation between two sinus signals is just treated as an additional contraction leading to a series of two subsequent interbeat intervals that are shorter than the sinus cycle length. The third interbeat interval is also a little shorter than normal, because, although the cycle length has not changed, the sinus signal immediately after the PVC has an increased delay duration shifting it closer to its successor. In the simulation without a sinus signal, the only qualitative difference can be observed in the case of an additional signal in between two AV node signals. The first interbeat interval is reduced, but the second interval is actually increased. Physiologically this can be explained by the signal of the PVC traveling upwards stimulating the AV node in the same way a signal from the sinus node would do. The next contraction can therefore only happen after the normal AV cycle length *and* the delay duration has passed.

#### 2.4 Methods: PVC model code

For the PVC model, we will now only discuss the changes required for the existing components. The full code can be found in Supplementary Listing 1–27.

In the `RefractoryGate`, the condition **`when`** `outp` simply has to be replaced with **`when`** `outp` **`or`** `reset`. For `ConductionDelay` the process is a little more complicated since oncoming signals have to be canceled. This is achieved by temporarily setting `t_next` to a very large value (larger than the total simulation time). The equations section changes as follows:

```
delay_passed = time > t_next or t_next > 1e99;
outp = edge(delay_passed);
when pre(reset) or (inp and pre(delay_passed)) then
  d_outp_inp = time - pre(t_last);
end when;
when pre(reset) then
  t_next = 1e100;
elsewhen inp and pre(delay_passed) then
  t_next = time + d_delay;
end when;
when outp or reset then
  t_last = time;
```

```
  end when;
```

The resulting extended model `ModularConductionX` looks as follows:

```
model ModularConductionX
  extends UnidirectionalConductionComponent(outp.fixed=true);
  extends SHMConduction.Icons.Heart;
  import SI = Modelica.SIunits;
  RefractoryGateX refrac_av(T_refrac=0.364)
    "refractory component for AV node" annotation(...);
  Pacemaker pace_av(T=1.7)
    "pacemaker effect of AV node" annotation(...);
  AVConductionDelayX delay_sa_v
    "delay between SA node and ventricles" annotation(...);
  RefractoryGate refrac_v(T_refrac=0.2)
    "refractory component for ventricles" annotation(...);
  InstantInput pvc(fixed=true)
    "trigger signal for a PVC" annotation(...);
  discrete SI.Period d_interbeat(start=1, fixed=true)
    "duration of last heart cycle";
  discrete SI.Time cont_last(start=0, fixed=true)
    "time of last contraction";f
  Modelica.Blocks.Logical.Or vcont
    "groups inputs for refrac_v" annotation(...);
  Modelica.Blocks.Logical.Or rpace
    "groups reset signals of pace_av" annotation(...);
  Modelica.Blocks.Logical.And pvc_upward
    "true if we have PVC that travels upward" annotation(...);
equation
  connect(inp, pace_av.inp) annotation(...);
  connect(pace_av.outp, refrac_av.inp) annotation(...);
  connect(refrac_av.outp, delay_sa_v.inp) annotation(...);
  connect(delay_sa_v.outp, vcont.u1) annotation(...);
  connect(refrac_av.outp, rpace.u2) annotation(...);
  connect(vcont.y, refrac_v.inp) annotation(...);
  connect(refrac_v.outp, outp) annotation(...);
  connect(outp, pvc_upward.u1) annotation(...);
  connect(pvc, pvc_upward.u2) annotation(...);
  connect(pvc_upward.y, refrac_av.reset) annotation(...);
  connect(pvc_upward.y, delay_sa_v.reset) annotation(...);
  connect(pvc_upward.y, rpace.u1) annotation(...);
  connect(rpace.y, pace_av.reset) annotation(...);
  connect(pvc, vcont.u2) annotation(...);
  when outp then
    d_interbeat = time - pre(cont_last);
    cont_last = time;
  end when;
end ModularConductionX;
```

#### 2.5 Methods: PVC experiment setup

The simulation experiment for the PVC model is also defined as a Modelica class. Here the variable `d_interbeat` was plotted once with `with_sinus = true` and once with `with_sinus = false`:

```
model PVCExample
    import SI = Modelica.SIunits;
  discrete SI.Time sig_last(start=0, fixed=true)
    "time where last SA/AV signal was received";
  Integer count_sig(start=0, fixed=true) "counts SA/AV signals";
  parameter Boolean with_sinus = true
    "if true, a sinus signal is applied, otherwise only the AV node is active";
  parameter SI.Period normal_interval = if with_sinus
    then 0.8 else con.pace_av.period "normal cycle duration without PVC";
  SI.Duration t_since_sig = time - pre(sig_last)
    "time since last signal from SA/AV node";
  Boolean pvc_a = pre(count_sig)==5 and t_since_sig > con.delay_sa_v.d_avc0/2
```

```
    "timer for PVC a): while 6th beat is delayed";
  Boolean pvc_b = pre(count_sig)==12 and t_since_sig > con.refrac_av.d_refrac/2
    "timer for PVC b): after 12th beat within refractory period";
  Boolean pvc_c = pre(count_sig)==19 and t_since_sig > normal_interval / 2
    "timer for PVC c): between 19th and 20th beat (after refractory period)";
  Boolean pvc_d = pre(count_sig)==26 and
    t_since_sig > normal_interval - con.delay_sa_v.d_avc0 / 2
    "timer for PVC d): just before the 27th beat was signalled";
  Boolean trigger(start=false, fixed=true) = pvc_a or pvc_b or pvc_c or pvc_d
    "pvc trigger signal";
equation
  con.pvc = edge(trigger);
  if with_sinus then
    con.inp = sample(0, normal_interval) "undisturbed normal sinus rhythm";
  else
    con.inp = false "no sinus, only AV node";
  end if;
  when con.refrac_av.outp then
    count_sig = pre(count_sig) + 1;
    sig_last = time;
  end when;
  annotation(
    experiment(
      StartTime = 0, StopTime = 55,
      Tolerance = 1e-6, Interval = 0.002
    ),
    __OpenModelica_simulationFlags(lv = "LOG_STATS", s = "dassl")
  );
end PVCExample;
```

#### References


1. Mendez, C., Gruhzit, C. C. & Moe, G. K. Influence of Cycle Length upon Refractory Period of Auricles, Ventricles, and A-V Node in the Dog. *American Journal of Physiology-Legacy Content* **184,** 287–295 (1956).
2. Schmidt, G. *et al.* Heart-Rate Turbulence after Ventricular Premature Beats as a Predictor of Mortality after Acute Myocardial Infarction. *The Lancet* **353,** 1390–1396 (1999).

#### Supplementary Figures

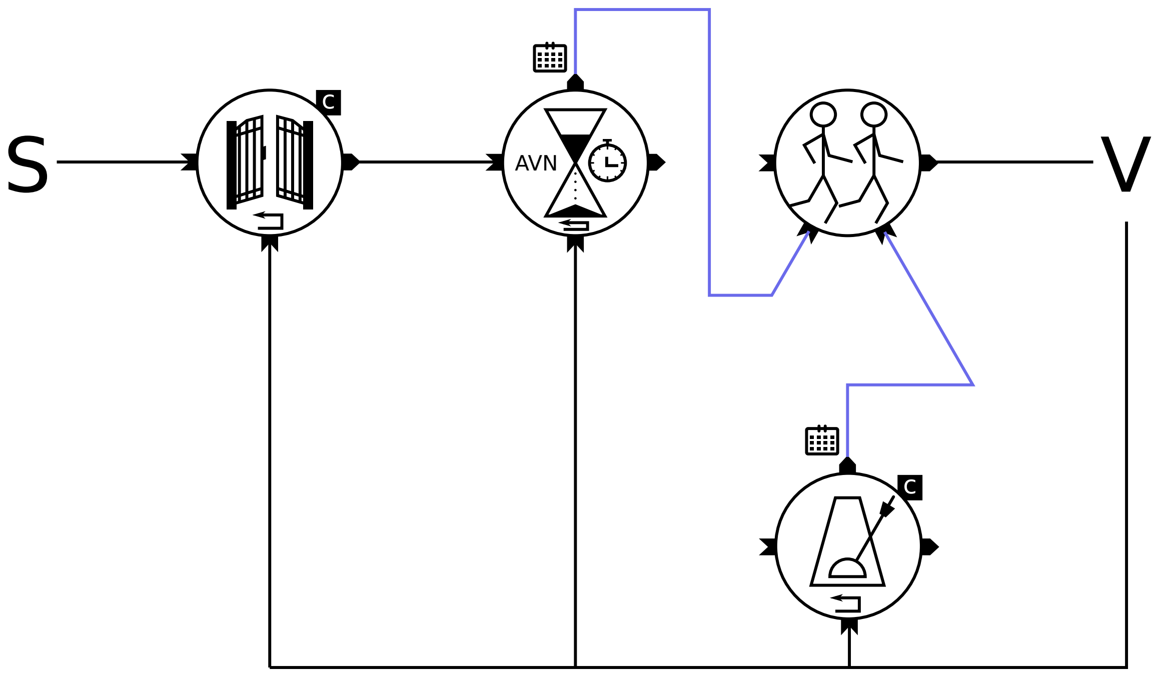


Supplementary Figure 1: Diagram of the monolithic model of the cardiac conduction system in the SHM. Sinus signals (S) are only propagated if enough time has passed since the last ventricular contraction (V). Propagated signals are delayed by a duration that also depends on the amount of time that has passed since the last ventricular contraction. The delay does result in a signal but in a scheduled time stamp that indicates the next time when a ventricular contraction could happen. This time stamp is compared to the time stamp produced by the pacemaker effect (a constant offset added to the time of the last ventricular contraction). The smaller time stamp wins the race condition (running stick figures symbol) and will trigger the next contraction while the larger time stamp is ignored.

This diagram illustrates why we had to change the model structure for our modular version. If we translated the monolithic version one-to-one into modules, we would have ended up with this quite convoluted diagram that does not separate the individual physiological effects very well. This would have had negative effects on the explanatory power and physiological soundness of the result.

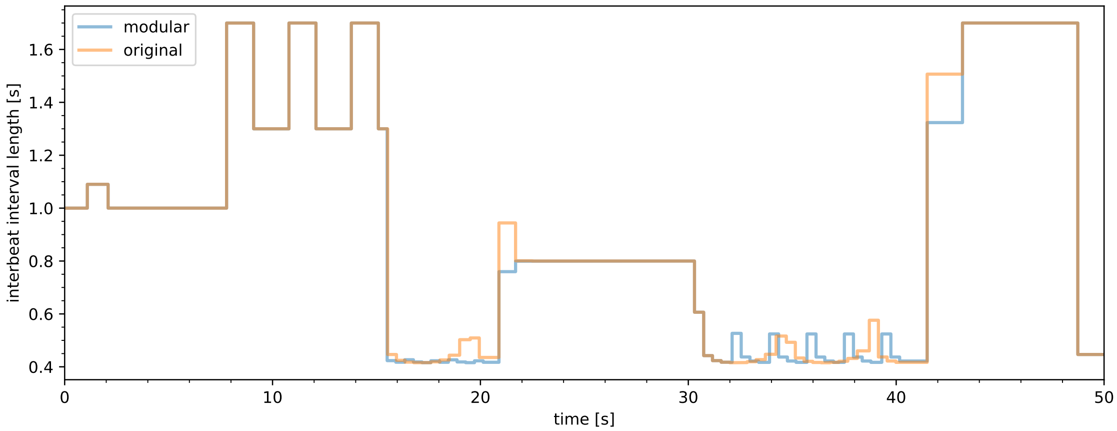


Supplementary Figure 2: Comparison of the interbeat intervals of the original conduction model (orange) and the modular version (blue). The plot was obtained with an artificial sinus signal that switches its cycle duration every ten seconds according to the following schedule: 1 s, 3 s, 0.05 s, 0.8 s, 0.2 s, 1.8 s. This schedule was chosen to cover a large range with different cycle durations that are a) below the refractory period of the AV node (0.2 s, 0.05 s), b) above the refractory period, but below the pacemaker period of the AV node (0.8 s, 1 s), or c) above the pacemaker period (3 s, 1.8 s). The schedule is not in any particular order so that both reactions to a sudden increase and a sudden decrease in sinus frequency can be observed. Only the cycle durations of 0.05 s and 0.2 s produce qualitative differences.

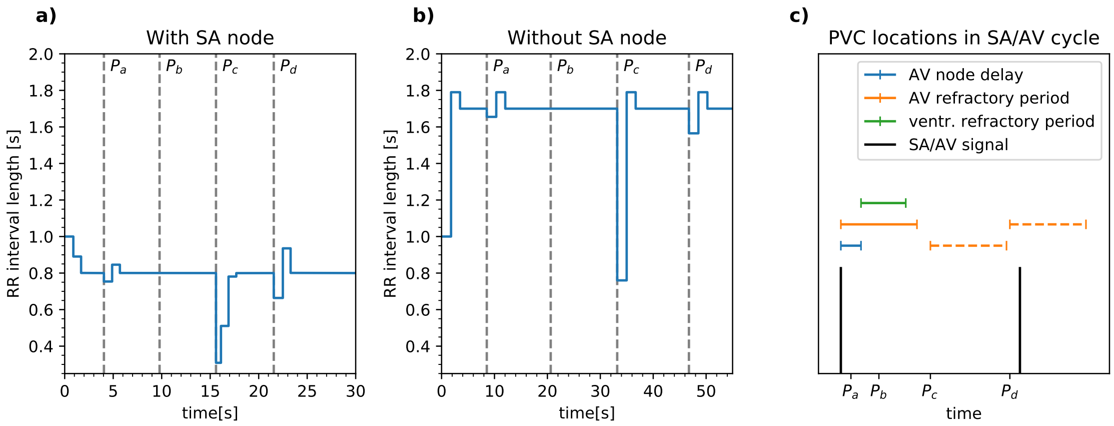


Supplementary Figure 3: Interbeat intervals of the modular model with PVCs with a) a normal sinus rhythm of 75 bpm and b) without sinus signal. Part c) shows the location of the ectopic beats in the normal cycle duration. PVCs are triggered with different prematurity: $P_a$ while a signal is delayed, $P_b$ within the ventricular refractory period, $P_c$ just between two beats, $P_d$ just before a beat would be triggered by the SA or AV node.

### 3 Supplementary Tables

| | | | | Python | | | | | | Julia | |
|---|---|---|---|---|---|---|---|---|---|---|---|
| | MATLAB[1] | SBML | CellML | pySB | PySCeS | SimuPy | PyDSTool | Antimony | Modelica | Modia | DiffEq.jl |
| modular | ✓ | (✓) | ✓ | (✓) | ✗ | (✓) | ✗ | ✓ | ✓ | ✓ | ✗ |
| instantiation / import | ✓ | ✓ | ✓ | (✓) | ✗ | ✓ | ✗ | ✓ | ✓ | ✓ | ✗ |
| object orientation | ✓ | ✗ | ✗ | ✗ | ✗ | ✗ | ✗ | ✗ | ✓ | ✓ | ✗ |
| multiple inheritance | (✓) | ✗ | ✗ | ✗ | ✗ | ✗ | ✗ | ✗ | ✓ | ✓ | ✗ |
| interface definition | ✓ | (✓) | ✓ | (✓) | ✗ | ✓ | ✗ | ✓ | ✓ | ✓ | ✗ |
| grouping of interface variables | ✓ | (✓) | ✗ | ✗ | ✗ | ✗ | ✗ | ✗ | ✓ | ✓ | ✗ |
| overwrite variables on import | ✗ | (✓) | ✗ | ✗ | ✗ | ✗ | ✗ | ✓ | ✓ | ✓ | ✗ |
| overwrite equations on import | ✗ | (✓) | ✗ | ✗ | ✗ | ✗ | ✗ | ✓ | (✓) | (✓) | ✗ |
| delete variables on import | ✗ | (✓) | ✗ | ✗ | ✗ | ✗ | ✗ | ✓ | ✗ | ✗ | ✗ |
| delete equations on import | ✗ | (✓) | ✗ | ✗ | ✗ | ✗ | ✗ | ✓ | ✗ | ✗ | ✗ |
| declarative | ✓ | ✓ | ✓ | (✓) | (✓) | (✓) | (✓) | ✓ | ✓ | ✓ | (✓) |
| mathematical semantics | ✓ | ✓ | ✓ | ✓ | ✓ | ✓ | ✓ | ✓ | ✓ | ✓ | ✓ |
| implicit ODE/DAE | ✓ | ✓ | (✓) | ✗ | ✗ | ✗ | ✗ | ✗ | ✓ | ✓ | (✓) |
| unit definition | ✓ | ✓ | ✓ | ✗ | ✓ | ✗ | ✗ | ✓ | ✓ | ✓ | ✓ |
| enforced unit definition | (✓) | ✗ | ✓ | ✗ | ✗ | ✗ | ✗ | ✗ | ✗ | ✗ | ✗ |
| ontology support | ✗ | ✓ | ✓ | (✓) | ✗ | ✗ | ✗ | ✗ | (✓) | (✓) | ✗ |
| readable | ✗ | (✓) | (✓) | (✓) | (✓) | (✓) | (✓) | (✓) | ✓ | ✓ | (✓) |
| model files focus on human-readability | ✗ [2] | (✓) | (✓) | ✓ | ✓ | (✓) | (✓) | ✓ | ✓ | ✓ | ✓ |
| format designed to be written by humans | ✗ [2] | ✗ | ✗ | ✓ | ✓ | ✓ | ✓ | ✓ | ✓ | ✓ | ✓ |
| labels for variables | ✓ | ✓ | ✓ | ✗ | ✗ | ✗ | ✗ | ✗ | ✓ | ✓ | ✗ |
| labels for models | ✓ | ✓ | ✓ | ✗ | ✗ | ✗ | ✗ | ✗ | ✓ | ✓ | ✗ |
| labels for equations | ✗ | ✓ | ✓ | ✗ | ✗ | ✗ | ✗ | ✗ | ✓ | ✓ | ✗ |
| rich text documentation | (✓) | ✓ | ✓ | ✗ | ✗ | ✗ | ✗ | ✗ | ✓ | ✓ | ✗ |
| open | ✗ | ✓ | ✓ | ✓ | ✓ | ✓ | ✓ | ✓ | (✓) | ✓ | ✓ |
| open-source language specification | ✗ | ✓ | ✓ | ✓ | ✓ | ✓ | ✓ | ✓ | ✓ | ✓ | ✓ |
| open-source compiler | ✗ | ✓ | ✓ | ✓ | ✓ | ✓ | ✓ | ✓ | (✓) | ✓ | ✓ |
| open-source tools and editors | ✗ | ✓ | ✓ | ✓ | ✓ | ✓ | ✓ | ✓ | (✓) | ✓ | ✓ |
| platform independent | ✗ | ✓ | ✓ | ✓ | ✓ | ✓ | ✓ | ✓ | ✓ | ✓ | ✓ |
| graphical | ✓ | ✓ | (✓) | ✗ | ✗ | ✗ | ✗ | ✗ | ✓ | ✗ | ✗ |
| image annotation | ✓ | ✓ | ✓ | ✗ | ✗ | ✗ | ✗ | ✗ | ✓ | ✗ | ✗ |
| vector graphics annotation | ✗ | ✓ | ✗ | ✗ | ✗ | ✗ | ✗ | ✗ | ✓ | ✗ | ✗ |
| annotations tied to model structure | ✓ | ✓ | ✗ | ✗ | ✗ | ✗ | ✗ | ✗ | ✓ | ✗ | ✗ |
| drag and drop composition | ✓ | (✓) | ✗ | ✗ | ✗ | ✗ | ✗ | ✗ | ✓ | ✗ | ✗ |
| hybrid | ✓ | (✓) | (✓) | ✗ | (✓) | (✓) | ✓ | (✓) | ✓ | ✓ | ✓ |
| ordinary differential equations (ODE) | ✓ | ✓ | ✓ | (✓) | ✓ | ✓ | ✓ | ✓ | ✓ | ✓ | ✓ |
| differential/algebraic equations (DAE) | ✓ | (✓) | (✓) | ✗ | (✓) | ✗ | ✓ | ✗ | ✓ | ✓ | ✓ |
| reinitialization due to discrete events | ✓ | ✓ | ✓ | ✗ | ✓ | ✓ | ✓ | ✓ | ✓ | ✓ | ✓ |
| explicit declaration of discrete variables | ✓ | ✗ | ✗ | ✗ | ✗ | ✗ | ✗ | ✗ | ✓ | ✓ | ✗ |
| other formalisms (Petri nets, FSA, ...) | ✓ | ✗ | ✗ | ✗ | ✗ | ✗ | ✗ | ✗ | ✓ | (✓) | ✗ |
| cross-language import/export (FMI) | (✓) | ✗ | ✗ | ✗ | ✗ | ✗ | ✗ | ✗ | ✓ | (✓) | ✗ |

[1] using the Simulink environment and the Simscape language
[2] while Simscape is human-readable, Simulink uses a proprietary binary file format

Supplementary Table 1: Evaluation of language candidates with respects to the desirable characteristics established in this paper broken up into individual features. A check mark in parentheses means the language has the respective feature in principle, but not to its full extent or with noticeable drawbacks. This table is an extended version of Table 1 in the main article.

### Supplementary Listings: Full code for monolithic and modular conduction models

The code snippets showcased in this paper where simplified for the convenience of the reader. This section of the supplement contains the full code that can also be obtained from GitHub [1]. The online version is the preferred source since it may contain fixes and updates applied after the publication of this paper. However, GitHub may not always be there in the future and therefore we provide this version to be archived along with the paper. Please note that most of the additional lines of code are introduced due to the graphical annotations.

#### Package structure

Modelica organizes code in packages. These packages can be defined within a single file or, as in this case, in a folder structure where each folder contains a file called `package.mo` that contains the package metadata. This project contains the following packages:

Supplementary Listing 1: `SHMConduction/package.mo`

```
package SHMConduction "modular and monolithic models of the human cardiac conduction system based on
      the PhD thesis of H. Seidel"
annotation(Documentation(info="<html>
  <p>Contains a modular version of the cardiac conduction system in the
  Seidel-Herzel-model (SHM).</p>

  <p>The SHM is a macro-level model of the human baroreflex that was
  originally written by Henrik Seidel in his PhD thesis in the language C
  <a href=\"#ref1\">[1]</a>.
  One part of this model is the cardiac conduction system that transfers
  signals from the sinus node to the ventricles where they trigger a
  ventricular contraction and thus the switch from diastole to systole.
  The cardiac conduction system is mainly controlled by the AV node that
  propagates sinus signals to the ventricles.</p>

  <p>The cardiac conduction submodel incorporates the following physiological
  effects.</p>

  <ul>
    <li><b>Refractory effect</b>: The atrioventricular (AV) node that reveives
    signals from the sinus node has a refractory period during which no
    excitation of the AV node can take place.</li>
    <li><b>Pacemaker effect</b>: If the sinus node does not send any signals
    for a prolonged time period, the AV node sends a signal on its own.</li>
    <li><b>Delay effect</b>: Signals traveling through the AV node are delayed.
    The duration of the delay depends on the time that has passed since the
    last signal has left the AV node.</li>
  </ul>

  <p>This package contains a monolithic version of the original model of the
  cardiac conduction system that was translated to Modelica, a modular version
  with small structural differences and an extension of this modular version
  that features a trigger for premature ventricular contractions (PVCs).</p>

  <p>The structural differences between the monolithic and modular versions
  are the following:</p>

  <table>
    <tr>
      <th>Monolithic version</th><th>Modular version</th>
    </tr>
```

[1] `https://github.com/CSchoel/shm-conduction`

```
    <tr>
      <td>Refractory time starts right after the contraction</td>
      <td>Refractory time starts right after sinus signal</td>
    </tr>
    <tr>
      <td>Pacemaker timer is reset after contraction</td>
      <td>Pacemaker timer is reset when sinus signal is received and AV node
      is not refractory</td>
    </tr>
    <tr>
      <td>Compares scheduled time stamps for next contraction that would be
      triggered by a delayed sinus signal or an intrinsic AV signal</td>
      <td>Every connection between components carries an actual signal;
      no scheduled time stamps are required</td>
    </tr>
  </table>

  <p>The Modelica version of the SHM model is described in detail in
  <a href=\"#ref2\">[2]</a>.
  The modular versions will be published in an upcoming paper
  <a href=\"#ref3\">[2]</a>.</p>

  <p><a name=\"ref1\">[1]</a> H. Seidel, &ldquo;Nonlinear dynamics of physiological
  rhythms,&rdquo; PhD thesis, Technische Universität Berlin, Berlin, Germany, 1997.
  </p>

  <p><a name=\"ref2\">[2]</a> C. Schölzel, A. Goesmann, G. Ernst, and A. Dominik,
  &ldquo;Modeling biology in Modelica: The human baroreflex,&rdquo; in Proceedings of the
  11th International Modelica Conference, Versailles, France, 2015, pp.-
  367376.</p>

  <p><a name=\"ref3\">[3]</a> C. Schölzel, V. Blesius, G. Ernst and
  A. Dominik, &ldquo;Required characteristics for modeling languages in systems
  biology: A software engineering perspective,&rdquo; unpublished.</p>
</html>"));
end SHMConduction;
```

Supplementary Listing 2: `SHMConduction/Components/package.mo`

```
within SHMConduction;
package Components "contains individual modules"
  type InstantSignal = Boolean(quantity="sum of Kronecker deltas")
    "signal that is only true for exact time instants (i.e. that behaves as a sum of Kronecker
         deltas)";
annotation(Documentation(info="<html>
  <p>Contains component models used in the examples.</p>
</html>"));
end Components;
```

Supplementary Listing 3: `SHMConduction/Components/Connectors/package.mo`

```
within SHMConduction.Components;
package Connectors "connector classes used as interfaces between components"
annotation(Documentation(info="<html>
  <p>Contains connector classes that define the interface between
  components.</p>
</html>"));
end Connectors;
```

Supplementary Listing 4: `SHMConduction/Components/PVC/package.mo`

```
within SHMConduction.Components;
package PVC "extended variants of components that allow to simulate premature ventricular
      contractions"
annotation(Documentation(info="<html>
  <p>Contains modified and additional models for simulating premature
```

```
  ventricular contractions (PVCs).</p>
</html>"));
end PVC;
```

Supplementary Listing 5: `SHMConduction/Examples/package.mo`

```
within SHMConduction;
package Examples "contains complete examples that are ready for simulation"
annotation(Documentation(info="<html>
  <p>Contains complete example models that can be simulated.</p>
</html>"));
end Examples;
```

Supplementary Listing 6: `SHMConduction/Icons/package.mo`

```
within SHMConduction;
package Icons "base models that contain no logic but only icon definitions"
annotation(Documentation(info="<html>
  <p>Contains empty classes with annotations that can be extended to use
  the respective icons in a model without cluttering the code.</p>
</html>"));
end Icons;
```

#### Monolithic model

Supplementary Listing 7: `SHMConduction/Components/MonolithicConduction.mo`

```
within SHMConduction.Components;
model MonolithicConduction "cardiac conduction system of the human heart adapted from the doctorate
    thesis of H. Seidel"
  import SI = Modelica.SIunits;
  input InstantSignal inp(start=false, fixed=true) "the sinus signal";
  output InstantSignal outp(start=false, fixed=true) "true when a contraction is triggered";
  parameter SI.Duration d_refrac = 0.22 "refractory period that has to pass until a signal from the
      sinus node can take effect again";
  parameter SI.Period av_period = 1.7 "av-node cycle duration";
  parameter SI.Duration k_avc_t = 0.78 "sensitivity of the atrioventricular conduction time to the
      time passed since the last ventricular conduction";
  parameter SI.Duration d_avc0 = 0.09 "base value for atrioventricular conduction time";
  parameter SI.Duration tau_avc = 0.11 "reference time for atrioventricular conduction time"; //TODO
       find better description
  parameter SI.Period initial_d_sinus_sinus = 1 "initial value for d_sinus_sinus";
  parameter SI.Period initial_d_interbeat = 1 "initial value for d_interbeat";
  parameter SI.Time initial_cont_last = 0 "initial value for last ventricular contraction time";
  parameter SI.Duration initial_d_delay = 0.15 "initial value for atrioventricular conduction time";
  output Boolean av_contraction "true when the av-node triggers a contraction";
  output Boolean sinus_contraction "true when the sinus node triggers a contraction";
  output Boolean refrac_passed(start=false, fixed=true) "true when the refractory period has passed"
      ;
  discrete output SI.Period d_sinus_sinus "time between the last two sinus signals that did trigger
      a contraction";
  discrete output SI.Period d_interbeat "time between the last two contractions";
protected
  discrete SI.Time cont_last "time of last contraction";
  discrete SI.Duration d_delay "atrioventricular conduction time (delay for sinus signal to trigger
      contraction)";
  SI.Duration since_cont "helper variable; time passed since last contraction";
  Boolean signal_received(start=false, fixed=true) "true, if a sinus signal has already been
      received since the last contraction";
  discrete SI.Time sinus_last "time of last received sinus signal";
  Boolean contraction_event(start=false, fixed=true);
initial equation
  cont_last = initial_cont_last;
  sinus_last = 0;
```

```
  d_sinus_sinus = initial_d_sinus_sinus;
  d_interbeat = initial_d_interbeat;
  d_delay = initial_d_delay;
equation
  signal_received = sinus_last > cont_last;
  refrac_passed = since_cont > d_refrac;
  contraction_event = (av_contraction or sinus_contraction) and refrac_passed "contraction can come
      from av-node or sinus node";
  outp = edge(contraction_event);
  av_contraction = since_cont > av_period "av-node contracts when av_period has passed since last
      contraction";
  sinus_contraction = signal_received and time > sinus_last + d_delay "sinus node contracts when
      d_delay has passed since last sinus signal";
  since_cont = time - cont_last;
  //sinus signal is recognized if refractory period has passed and there is no other sinus signal
      already in effect
  when inp and pre(refrac_passed) and not pre(signal_received) then
    d_delay = d_avc0 + k_avc_t * exp(-since_cont / tau_avc) "schedules next sinus_contraction";
    sinus_last = time "record timestamp of recognized sinus signal";
    d_sinus_sinus = time - pre(sinus_last);
  end when;
  when pre(outp) then
    cont_last = time "record timestamp of contraction";
    d_interbeat = time - pre(cont_last);
  end when;
  annotation(Documentation(info = "<html>
  <p>Models the contraction of the heart as described in Seidel's thesis.</p>
  <p>The model takes into account the following effects:</p>
  <ul>
    <li>There is a refractory period of duration <b>d_refrac</b> after a contraction during which
        signals of the sinus node are ignored.
    <li>If no sinus-induced contraction occurs for a prolonged time span (namely <b>av_period</b>)
        the av-node initiates a contraction by itself.
    <li>When a sinus signal is received, the upper heart contracts pumping the blood from the atrium
         into the ventricles. The systole does only
    begin with the second contraction of the heart. The time period between these two events is
        called the &quot;atrioventricular conduction time&quot;.
  </ul>
  <p><i>Note: The formulas in this model differ from the formulas found in the c-implementation by
      Seidel because OpenModelica is currently
  not capable of handling discrete equation systems. Therefore it was necessary to introduce the
      continuous phases <b>av_phase</b>,
  <b>sinus_phase</b> and <b>refrac_countdown</b>, as well as the continuous variable condition <b>
      signal_received_cont</b>.</i></p>
</html>"));
end MonolithicConduction;
```

### Modular model

Supplementary Listing 8: SHMConduction/Components/Connectors/InstantInput.mo

```
within SHMConduction.Components.Connectors;
connector InstantInput = input InstantSignal "input with Kronecker delta behavior"
  annotation(
    Icon(
      coordinateSystem(
        preserveAspectRatio= false,
        extent= {{-100,-100},{100,100}}
      ),
      graphics= {
        Polygon(
          origin= {-100,100},
          lineThickness= 5,
          pattern= LinePattern.None,
          points= {{194.32, -100}, {200, -200}, {0, -200}, {108.45, -100}, {0, 0}, {200, 0}},
          fillPattern= FillPattern.Solid,
          fillColor= {0,0,0},
          rotation= 0
```

```
        )
      }
    )
  )
;
```

Supplementary Listing 9: `SHMConduction/Components/Connectors/InstantOutput.mo`

```
within SHMConduction.Components.Connectors;
connector InstantOutput = output InstantSignal "output with Kronecker delta behavior"
  annotation(
    Icon(
      coordinateSystem(
        preserveAspectRatio= false,
        extent= {{-100,-100},{100,100}}
      ),
      graphics= {
        Polygon(
          origin= {-100,100},
          lineThickness= 5,
          pattern= LinePattern.None,
          points= {{200, -100}, {91.55, 0}, {0, 0}, {0, -200}, {91.55, -200}, {200, -100}},
          fillPattern= FillPattern.Solid,
          fillColor= {0,0,0},
          rotation= 0
        )
      }
    )
  )
;
```

Supplementary Listing 10: `SHMConduction/Components/UnidirectionalConductionComponent.mo`

```
within SHMConduction.Components;
partial model UnidirectionalConductionComponent "basic interface class with single input and output"
  import SHMConduction.Components.Connectors.{InstantInput, InstantOutput};
  InstantInput inp "input connector" annotation(
    Placement(
      visible = true,
      transformation(
        origin = {-100, 0},
        extent = {{-10, -10}, {10, 10}}, rotation = 0
      ),
      iconTransformation(
        origin = {-108, 0},
        extent = {{-10, -10}, {10, 10}}, rotation = 0
      )
    )
  );
  InstantOutput outp "output connector" annotation(
    Placement(
      visible = true,
      transformation(
        origin = {102, 0},
        extent = {{-10, -10}, {10, 10}}, rotation = 0
      ),
      iconTransformation(
        origin = {108, 0},
        extent = {{-10, -10}, {10, 10}}, rotation = 0
      )
    )
  );
  annotation(
    Icon(
      coordinateSystem(
        preserveAspectRatio= false,
        extent= {{-100,-100},{100,100}}
```

```
    ),
    graphics= {
      Ellipse(
        origin= {-100,100},
        lineThickness= 2,
        pattern= LinePattern.Solid,
        fillPattern= FillPattern.None,
        extent= {{2,-2},{198,-198}},
        rotation= 0.0
      )
    }
  )
);
end UnidirectionalConductionComponent;
```

Supplementary Listing 11: `SHMConduction/Components/Resettable.mo`

```
within SHMConduction.Components;
partial model Resettable "base class for all components that need a reset input"
  import SHMConduction.Components.Connectors.InstantInput;
  InstantInput reset "signal that resets internal variables" annotation(
    Placement(
      visible = true,
      transformation(
        origin = {-2, -98},
        extent = {{-10, -10}, {10, 10}}, rotation = 0
      ),
      iconTransformation(
        origin = {0, -108},
        extent = {{-10, -10}, {10, 10}}, rotation = 90
      )
    )
  );
  annotation(
    Icon(
      coordinateSystem(
        preserveAspectRatio= false,
        extent= {{-100,-100},{100,100}}
      ),
      graphics= {
        Line(
          origin= {-100,100},
          pattern= LinePattern.Solid,
          thickness= 1.5,
          arrowSize= 5,
          points= {{82.25, -192.58}, {114.50, -192.58}, {114.50, -180.64}, {81.84, -180.64}},
          arrow= {Arrow.None, Arrow.Open},
          rotation= 0
        )
      }
    )
  );
end Resettable;
```

Supplementary Listing 12: `SHMConduction/Components/Pacemaker.mo`

```
within SHMConduction.Components;
model Pacemaker "pacemaker that can elicit spontaneous signals and transmit incoming signals"
  extends UnidirectionalConductionComponent;
  extends SHMConduction.Icons.Metronome;
  extends SHMConduction.Icons.Constant;
  extends Resettable(reset.fixed=true); // resets internal clock
  import SI = Modelica.SIunits;
  parameter SI.Period period = 1 "pacemaker period";
protected
  discrete SI.Time t_next(start=period, fixed=true)
    "scheduled time of next spontaneous beat";
```

```
  InstantSignal spontaneous_signal = time > pre(t_next)
    "signal generated spontaneously by this pacemaker";
equation
  outp = inp or spontaneous_signal;
  when spontaneous_signal or pre(reset) then
    t_next = time + period;
  end when;
end Pacemaker;
```

Supplementary Listing 13: `SHMConduction/Components/RefractoryGate.mo`

```
within SHMConduction.Components;
model RefractoryGate "refractory gate that can block or transmit signals"
  extends UnidirectionalConductionComponent;
  extends SHMConduction.Icons.Gate;
  extends SHMConduction.Icons.Constant;
  import SI = Modelica.SIunits;
  parameter SI.Time t_first = 0 "time of first signal";
  parameter SI.Duration d_refrac = 1 "duration of refractory period";
  Boolean refrac_passed = time - pre(t_last) > d_refrac "true if component is ready to receive a
      signal";
protected
  discrete SI.Time t_last(start=t_first, fixed=true) "time of last output";
equation
  outp = inp and refrac_passed;
  when outp then
    t_last = time;
  end when;
end RefractoryGate;
```

Supplementary Listing 14: `SHMConduction/Components/ConductionDelay.mo`

```
within SHMConduction.Components;
partial model ConductionDelay "cardiac conduction delay that depends on previous cycle duration"
  extends UnidirectionalConductionComponent;
  extends SHMConduction.Icons.Hourglass;
  import SI = Modelica.SIunits;
  discrete SI.Duration d_delay "delay duration";
  Boolean delay_passed(start=false, fixed=true) = time > t_next "if false, there is still a signal
      currently put on hold";
protected
  discrete SI.Duration d_outp_inp(start=0, fixed=true) "time between last output and following
      signal";
  discrete SI.Time t_last(start=0, fixed=true) "time of last output";
  discrete SI.Time t_next(start=-1, fixed=true) "scheduled time of next output";
equation
  outp = edge(delay_passed);
  when inp and pre(delay_passed) then
    d_outp_inp = time - pre(t_last);
    t_next = time + d_delay;
  end when;
  when outp then
    t_last = time;
  end when;
end ConductionDelay;
```

Supplementary Listing 15: `SHMConduction/Components/AVConductionDelay.mo`

```
within SHMConduction.Components;
model AVConductionDelay "conduction delay between SA node and ventricles"
  extends ConductionDelay;
  import SI = Modelica.SIunits;
  parameter SI.Duration k_avc_t = 0.78 "maximum increase in delay duration";
  parameter SI.Duration d_avc0 = 0.09 "minimal delay duration";
  parameter SI.Duration tau_avc = 0.11 "reference time for delay duration";
  parameter SI.Duration initial_d_avc = 0.15 "initial value for delay duration";
```

```
initial equation
  d_delay = initial_d_avc;
equation
  when inp and pre(delay_passed) then
    d_delay = d_avc0 + k_avc_t * exp(-d_outp_inp/tau_avc);
  end when;
  annotation(
    Icon(
      coordinateSystem(
        preserveAspectRatio= false,
        extent= {{-100,-100},{100,100}}
      ),
      graphics= {
        Text(
          origin = {-55, 1},
          extent = {{-21, 19}, {27, -23}},
          textString = "AVN"
        )
      }
    )
  );
end AVConductionDelay;
```

Supplementary Listing 16: `SHMConduction/Components/ModularConduction.mo`

```
within SHMConduction.Components;
model ModularConduction "modular version of the model of the cardiac conduction system by H. Seidel"
  extends UnidirectionalConductionComponent;
  extends SHMConduction.Icons.Heart;
  import SI = Modelica.SIunits;
  RefractoryGate refrac_av(d_refrac=0.364) "refractory component for AV node" annotation(
    Placement(
      visible = true,
      transformation(
        origin = {6, 5.77316e-15},
        extent = {{-20, -20}, {20, 20}}, rotation = 0
      )
    )
  );
  Pacemaker pace_av(period=1.7) "pacemaker effect of AV node" annotation(
    Placement(
      visible = true,
      transformation(
        origin = {-52, 3.77476e-15},
        extent = {{-20, -20}, {20, 20}}, rotation = 0
      )
    )
  );
  AVConductionDelay delay_sa_v "total delay between SA node and ventricles" annotation(
    Placement(
      visible = true,
      transformation(
        origin = {62, 3.55271e-15},
        extent = {{-20, -20}, {20, 20}}, rotation = 0
      )
    )
  );
  discrete SI.Duration d_interbeat(start=1, fixed=true) "duration of last heart cycle (interbeat
      interval)";
  discrete SI.Time cont_last(start=0, fixed=true) "time of last contraction";
equation
  connect(inp, pace_av.inp) annotation(
    Line(thickness = 1, points = {{-74, 0}, {-96, 0}, {-96, 0}, {-100, 0}})
  );
  connect(pace_av.outp, refrac_av.inp) annotation(
    Line(thickness = 1, points = {{-30, 0}, {-16, 0}, {-16, 0}, {-16, 0}})
  );
  connect(refrac_av.outp, pace_av.reset) annotation(
```

```
    Line(thickness = 1, points = {{28, 0}, {34, 0}, {34, -44}, {-52, -44}, {-52, -22}, {-52, -22}})
  );
  connect(refrac_av.outp, delay_sa_v.inp) annotation(
    Line(thickness = 1, points = {{28, 0}, {42, 0}, {42, 0}, {40, 0}})
  );
  connect(delay_sa_v.outp, outp) annotation(
    Line(thickness = 1, points = {{84, 0}, {102, 0}, {102, 0}, {102, 0}})
  );
  when outp then
    d_interbeat = time - pre(cont_last);
    cont_last = time;
  end when;
  annotation(
    Icon(
      graphics = {
        Line(
          origin = {-75, 5},
          points = {{-17, -5}, {17, 5}},
          arrow = {Arrow.None, Arrow.Open},
          thickness = 1,
          arrowSize = 5
        ),
        Line(
          origin = {75, -20},
          points = {{-19, -18}, {19, 18}},
          arrow = {Arrow.None, Arrow.Open},
          thickness = 1,
          arrowSize = 5
        )
      }
    )
  );
end ModularConduction;
```

### PVC extension

Supplementary Listing 17: SHMConduction/Components/PVC/ConductionDelayX.mo

```
within SHMConduction.Components.PVC;
partial model ConductionDelayX "resettable variant of ConductionDelay (resetting cancels delayed
    signals)"
  extends UnidirectionalConductionComponent;
  extends SHMConduction.Icons.Hourglass;
  extends Resettable(reset.fixed=true); // cancels a signal that is currently on hold
  import SI = Modelica.SIunits;
  discrete SI.Duration d_delay "delay duration";
  Boolean delay_passed(start=false, fixed=true) = time > t_next or t_next > 1e99 "if false, there is
      still a signal currently put on hold";
protected
  discrete SI.Duration d_outp_inp(start=0, fixed=true) "time between last output and following
      signal";
  discrete SI.Time t_last(start=0, fixed=true) "time of last output";
  discrete SI.Time t_next(start=-1, fixed=true) "scheduled time of next output";
equation
  outp = edge(delay_passed);
  when pre(reset) or (inp and pre(delay_passed)) then
    d_outp_inp = time - pre(t_last);
  end when;
  when pre(reset) then
    t_next = 1e100;
  elsewhen inp and pre(delay_passed) then
    t_next = time + d_delay;
  end when;
  when outp or reset then
    t_last = time;
  end when;
end ConductionDelayX;
```

Supplementary Listing 18: `SHMConduction/Components/PVC/AVConductionDelayX.mo`

```
within SHMConduction.Components.PVC;
model AVConductionDelayX "resettable variant of AVConductionDelay (resetting cancels delayed signals
    )"
  extends ConductionDelayX;
  import SI = Modelica.SIunits;
  parameter SI.Duration k_avc_t = 0.78 "maximum increase in delay duration";
  parameter SI.Duration d_avc0 = 0.09 "minimal delay duration";
  parameter SI.Duration tau_avc = 0.11 "reference time for delay duration";
  parameter SI.Duration initial_d_avc = 0.15 "initial value for delay duration";
initial equation
  d_delay = initial_d_avc;
equation
  when inp and pre(delay_passed) then
    d_delay = d_avc0 + k_avc_t * exp(-d_outp_inp/tau_avc);
  end when;
  annotation(
    Icon(
      coordinateSystem(
        preserveAspectRatio= false,
        extent= {{-100,-100},{100,100}}
      ),
      graphics= {
        Text(
          origin = {-55, 1},
          extent = {{-21, 19}, {27, -23}},
          textString = "AVN"
        )
      }
    )
  );
end AVConductionDelayX;
```

Supplementary Listing 19: `SHMConduction/Components/PVC/RefractoryGateX.mo`

```
within SHMConduction.Components.PVC;
model RefractoryGateX "resettable variant of RefractoryGate"
  extends UnidirectionalConductionComponent;
  extends SHMConduction.Icons.Gate;
  extends SHMConduction.Icons.Constant;
  extends Resettable; // resets internal clock
  import SI = Modelica.SIunits;
  parameter SI.Time t_first = 0 "time of first signal";
  parameter SI.Duration d_refrac = 1 "refractory period";
  Boolean refrac_passed = time - pre(t_last) > d_refrac "true if component is ready to receive a
      signal";
protected
  discrete SI.Time t_last(start=t_first, fixed=true) "time of last output";
equation
  outp = inp and refrac_passed;
  when outp or reset then
    t_last = time;
  end when;
end RefractoryGateX;
```

Supplementary Listing 20: `SHMConduction/Components/PVC/ModularConductionX.mo`

```
within SHMConduction.Components.PVC;
model ModularConductionX "cardiac conduction system with trigger for PVCs"
  extends UnidirectionalConductionComponent(outp.fixed=true);
  // outp is used in a when equation, so we need an initial value
  extends SHMConduction.Icons.Heart;
  import SHMConduction.Components.Connectors.InstantInput;
  import SI = Modelica.SIunits;
  RefractoryGateX refrac_av(d_refrac=0.364) "refractory component for AV node" annotation(
    Placement(
      visible = true,
```

```
    transformation(
      origin = {-20, 0},
      extent = {{-10, -10}, {10, 10}}, rotation = 0
    )
  )
);
Pacemaker pace_av(period=1.7) "pacemaker effect of AV node" annotation(
  Placement(
    visible = true,
    transformation(
      origin = {-60, 0},
      extent = {{-10, -10}, {10, 10}}, rotation = 0
    )
  )
);
AVConductionDelayX delay_sa_v "total delay between SA node and ventricles" annotation(
  Placement(
    visible = true,
    transformation(
      origin = {20, 0},
      extent = {{-10, -10}, {10, 10}}, rotation = 0
    )
  )
);
RefractoryGate refrac_v(d_refrac=0.2) "refractory component for ventricles" annotation(
  Placement(
    visible = true,
    transformation(
      origin = {60, 0},
      extent = {{-10, -10}, {10, 10}}, rotation = 0
    )
  )
);
discrete SI.Period d_interbeat(start=1, fixed=true) "duration of last heart cycle";
discrete SI.Time cont_last(start=0, fixed=true) "time of last contraction";
InstantInput pvc(fixed=true) "trigger signal for a PVC" annotation(
  Placement(
    visible = true,
    transformation(
      origin = {76, -76},
      extent = {{-10, -10}, {10, 10}}, rotation = 135
    ),
    iconTransformation(
      origin = {76, -76},
      extent = {{-10, -10}, {10, 10}}, rotation = 135
    )
  )
);
Modelica.Blocks.Logical.Or vcont "groups inputs for refrac_v" annotation(
  Placement(
    visible = true,
    transformation(
      origin = {42, -36},
      extent = {{-10, -10}, {10, 10}}, rotation = 0
    )
  )
);
Modelica.Blocks.Logical.Or rpace "groups reset signals of pace_av" annotation(
  Placement(
    visible = true,
    transformation(
      origin = {-60, -38},
      extent = {{-10, -10}, {10, 10}}, rotation = 0
    )
  )
);
Modelica.Blocks.Logical.And pvc_upward "true if we have PVC that travels upward" annotation(
  Placement(
    visible = true,
```

```
      transformation(
        origin = {20, -72},
        extent = {{10, -10}, {-10, 10}}, rotation = 0
      )
    )
  );
equation
  connect(inp, pace_av.inp) annotation(
    Line(points = {{-100, 0}, {-72, 0}, {-72, 0}, {-70, 0}})
  );
  connect(pace_av.outp, refrac_av.inp) annotation(
    Line(points = {{-50, 0}, {-30, 0}, {-30, 0}, {-30, 0}})
  );
  connect(refrac_av.outp, delay_sa_v.inp) annotation(
    Line(points = {{-10, 0}, {10, 0}, {10, 0}, {10, 0}})
  );
  connect(delay_sa_v.outp, vcont.u1) annotation(
    Line(points = {{30, 0}, {34, 0}, {34, -22}, {16, -22}, {16, -36}, {30, -36}, {30, -36}})
  );
  connect(refrac_av.outp, rpace.u2) annotation(
    Line(points = {{-10, 0}, {-6, 0}, {-6, -58}, {-78, -58}, {-78, -46}, {-72, -46}, {-72, -46}})
  );
  connect(vcont.y, refrac_v.inp) annotation(
    Line(points = {{54, -36}, {60, -36}, {60, -12}, {42, -12}, {42, 0}, {50, 0}, {50, 0}}, color =
        {255, 0, 255})
  );
  connect(refrac_v.outp, outp) annotation(
    Line(points = {{70, 0}, {98, 0}, {98, 0}, {102, 0}})
  );
  connect(outp, pvc_upward.u1) annotation(
    Line(points = {{102, 0}, {84, 0}, {84, -54}, {50, -54}, {50, -72}, {32, -72}, {32, -72}})
  );
  connect(pvc, pvc_upward.u2) annotation(
    Line(points = {{76, -76}, {50, -76}, {50, -80}, {32, -80}, {32, -80}})
  );
  connect(pvc_upward.y, refrac_av.reset) annotation(
    Line(points = {{10, -72}, {-20, -72}, {-20, -10}, {-20, -10}}, color = {255, 0, 255})
  );
  connect(pvc_upward.y, delay_sa_v.reset) annotation(
    Line(points = {{10, -72}, {2, -72}, {2, -18}, {20, -18}, {20, -10}, {20, -10}}, color = {255, 0,
         255})
  );
  connect(pvc_upward.y, rpace.u1) annotation(
    Line(points = {{10, -72}, {-84, -72}, {-84, -38}, {-72, -38}, {-72, -38}}, color = {255, 0,
        255})
  );
  connect(rpace.y, pace_av.reset) annotation(
    Line(points = {{-48, -38}, {-44, -38}, {-44, -16}, {-60, -16}, {-60, -10}, {-60, -10}}, color =
        {255, 0, 255})
  );
  connect(pvc, vcont.u2) annotation(
    Line(points = {{76, -76}, {70, -76}, {70, -50}, {16, -50}, {16, -44}, {30, -44}})
  );
  when outp then
    d_interbeat = time - pre(cont_last);
    cont_last = time;
  end when;
  annotation(
    Icon(
      graphics = {
        Line(
          origin = {-75, 5},
          points = {{-17, -5}, {17, 5}},
          arrow = {Arrow.None, Arrow.Open},
          thickness = 1,
          arrowSize = 5
        ),
        Line(
          origin = {75, -20},
```

```
          points = {{-19, -18}, {19, 18}},
          arrow = {Arrow.None, Arrow.Open},
          thickness = 1,
          arrowSize = 5
        ),
        Line(
          origin = {60, -64},
          points = {{6, 0}, {-6, 0}},
          arrow = {Arrow.None, Arrow.Open},
          thickness = 1,
          arrowSize = 5
        )
      }
    )
  );
end ModularConductionX;
```

#### Simulation examples

Supplementary Listing 21: `SHMConduction/Examples/ModularExample.mo`

```
within SHMConduction.Examples;
// TODO: add ontolgy links with annotation(__Ontology(foo="bar"))
model ModularExample "experiment to compare Conduction and ModularConduction"
  SHMConduction.Components.ModularConduction modC "modular contraction model";
  SHMConduction.Components.MonolithicConduction monC "original monolithic contraction model";
equation
  modC.inp = monC.inp;
  if time < 5 then
    monC.inp = sample(0,1);
  elseif time < 15 then
    monC.inp = sample(0,3);
  elseif time < 20 then
    // during Afib, atrial impulses occur at up to 600/min => with distance 0.1s
    // source: https://my.clevelandclinic.org/health/diseases/16765-atrial-fibrillation-afib
    monC.inp = sample(0,0.05);
  elseif time < 30 then
    monC.inp = sample(0,0.8);
  elseif time < 40 then
    monC.inp = sample(0,0.2);
  else
    monC.inp = sample(0,1.8);
  end if;
  annotation(
    experiment(StartTime = 0, StopTime = 50, Tolerance = 1e-6, Interval = 0.002),
    __OpenModelica_simulationFlags(lv = "LOG_STATS", s = "dassl")
  );
end ModularExample;
```

Supplementary Listing 22: `SHMConduction/Examples/PVCExample.mo`

```
within SHMConduction.Examples;
model PVCExample "experiment to test response of ModularConductionX to PVCs"
  SHMConduction.Components.PVC.ModularConductionX con;
  import SI = Modelica.SIunits;
  discrete SI.Time sig_last(start=0, fixed=true) "time where last SA/AV signal was received";
  Integer count_sig(start=0, fixed=true) "counts SA/AV signals";
  parameter Boolean with_sinus = true "if true, a sinus signal is applied, otherwise only the AV
      node is active";
  parameter SI.Period normal_interval = if with_sinus then 0.8 else con.pace_av.period "normal cycle
       duration without PVC";
  SI.Duration t_since_sig = time - pre(sig_last) "time since last signal from SA/AV node";
  Boolean pvc_a = pre(count_sig) == 5 and t_since_sig > con.delay_sa_v.d_avc0 / 2
    "timer for PVC a): while 6th beat is delayed";
  Boolean pvc_b = pre(count_sig) == 12 and t_since_sig > con.refrac_av.d_refrac / 2
    "timer for PVC b): after 12th beat within refractory period";
  Boolean pvc_c = pre(count_sig) == 19 and t_since_sig > normal_interval / 2
```

```
    "timer for PVC c): between 19th and 20th beat (after refractory period)";
  Boolean pvc_d = pre(count_sig) == 26 and
    t_since_sig > normal_interval - con.delay_sa_v.d_avc0 / 2
    "timer for PVC d): just before the 27th beat was signalled";
  Boolean trigger(start=false, fixed=true) = pvc_a or pvc_b or pvc_c or pvc_d "pvc trigger signal";
equation
  con.pvc = edge(trigger);
  if with_sinus then
    con.inp = sample(0, normal_interval) "undisturbed normal sinus rhythm";
  else
    con.inp = false "no sinus, only AV node";
  end if;
  when con.refrac_av.outp then
    count_sig = pre(count_sig) + 1;
    sig_last = time;
  end when;
  annotation(
    experiment(StartTime = 0, StopTime = 55, Tolerance = 1e-6, Interval = 0.002),
    __OpenModelica_simulationFlags(lv = "LOG_STATS", s = "dassl")
  );
end PVCExample;
```

#### Icons

Supplementary Listing 23: `SHMConduction/Icons/Constant.mo`

```
within SHMConduction.Icons;
model Constant "small box with a C in upper right corner"
    annotation(
        Icon(
            coordinateSystem(
                preserveAspectRatio= false,
                extent= {{-100,-100},{100,100}}
            ),
            graphics= {
                Polygon(
                    origin= {-100,100},
                    lineThickness= 0.56,
                    pattern= LinePattern.None,
                    points= {{163.21, -24.39}, {163.21, -2.15}, {196.78, -2.15}, {196.78, -36.36},
                         {174.62, -36.36}},
                    fillPattern= FillPattern.Solid,
                    fillColor= {0,0,0},
                    rotation= 0
                ),
                Text(
                    origin = {81, 84},
                    lineColor = {255, 255, 255},
                    extent = {{-11, 22}, {11, -22}},
                    textString = "c"
                )
            }
        )
    );
end Constant;
```

Supplementary Listing 24: `SHMConduction/Icons/Gate.mo`

```
within SHMConduction.Icons;
model Gate "gate icon for refractory period"
  annotation(
    Icon(
      coordinateSystem(
        preserveAspectRatio= false,
        extent= {{-100,-100},{100,100}}
      ),
      graphics= {
```

```
Rectangle(
  origin= {-100,100},
  lineThickness= 1,
  pattern= LinePattern.None,
  fillPattern= FillPattern.Solid,
  fillColor= {0,0,0},
  extent= {{42.33,-42.01},{55.18,-160.69}},
  rotation= 0
),
Polygon(
  origin= {-100,100},
  lineThickness= 2,
  pattern= LinePattern.Solid,
  points= {{69.55, -41.25}, {49.33, -57.13}, {49.33, -153.89}, {89.79, -142.17}, {89.79,
       -35.96}},
  fillPattern= FillPattern.None,
  rotation= 0
),
Rectangle(
  origin= {-100,100},
  lineThickness= 1,
  pattern= LinePattern.None,
  fillPattern= FillPattern.Solid,
  fillColor= {0,0,0},
  extent= {{145.47,-42.01},{158.32,-160.69}},
  rotation= 0
),
Polygon(
  origin= {-100,100},
  lineThickness= 2,
  pattern= LinePattern.Solid,
  points= {{131.10, -41.25}, {151.33, -57.13}, {151.33, -153.89}, {110.86, -142.17},
       {110.86, -35.96}},
  fillPattern= FillPattern.None,
  rotation= 0
),
Polygon(
  origin= {-100,100},
  lineThickness= 1,
  pattern= LinePattern.None,
  points= {{90.34, -77}, {94.87, -75.65}, {94.87, -93.03}, {89.96, -94.54}},
  fillPattern= FillPattern.Solid,
  fillColor= {0,0,0},
  rotation= 0
),
Line(
  origin= {-100,100},
  pattern= LinePattern.Solid,
  rotation= 0,
  points= {{69.55, -41.25}, {69.55, -148.03}},
  thickness= 2
),
Line(
  origin= {-100,100},
  pattern= LinePattern.Solid,
  rotation= 0,
  points= {{79.67, -37.80}, {79.67, -144.02}},
  thickness= 2
),
Line(
  origin= {-100,100},
  pattern= LinePattern.Solid,
  rotation= 0,
  points= {{59.56, -48.82}, {59.56, -150.20}},
  thickness= 2
),
Line(
  origin= {-100,100},
  pattern= LinePattern.Solid,
```

```
        rotation= 0,
        points= {{49.33, -62.89}, {89.79, -50.86}},
        thickness= 2
      ),
      Line(
        origin= {-100,100},
        pattern= LinePattern.Solid,
        rotation= 0,
        points= {{89.79, -132.65}, {49.33, -145.21}},
        thickness= 2
      ),
      Line(
        origin= {-100,100},
        pattern= LinePattern.Solid,
        rotation= 0,
        points= {{131.10, -41.25}, {131.10, -148.03}},
        thickness= 2
      ),
      Line(
        origin= {-100,100},
        pattern= LinePattern.Solid,
        rotation= 0,
        points= {{120.98, -37.80}, {120.98, -144.02}},
        thickness= 2
      ),
      Line(
        origin= {-100,100},
        pattern= LinePattern.Solid,
        rotation= 0,
        points= {{141.10, -48.82}, {141.10, -150.20}},
        thickness= 2
      ),
      Line(
        origin= {-100,100},
        pattern= LinePattern.Solid,
        rotation= 0,
        points= {{151.33, -62.89}, {110.86, -50.86}},
        thickness= 2
      ),
      Line(
        origin= {-100,100},
        pattern= LinePattern.Solid,
        rotation= 0,
        points= {{110.86, -132.65}, {151.33, -145.21}},
        thickness= 2
      )
    }
  )
);
end Gate;
```

Supplementary Listing 25: `SHMConduction/Icons/Heart.mo`

```
within SHMConduction.Icons;
model Heart "heart icon for full model of cardiac conduction system"
  annotation(
    Icon(
      coordinateSystem(
        preserveAspectRatio= false,
        extent= {{-100,-100},{100,100}}
      ),
      graphics= {
        Polygon(
          origin= {-100,100},
          lineThickness= 0.53,
          pattern= LinePattern.Solid,
          points= {{63.36, -149.76}, {77.13, -163.94}, {94.56, -172.20}, {114.76, -177.40}, {132.19,
               -179.84}, {143.62, -175.56}, {149.94, -164.14}, {152.69, -148.64}, {150.65,
```

```
        -132.02}, {145.04, -114.89}, {138.62, -102.65}, {130.15, -100.71}, {119.96, -96.02},
        {106.90, -89.39}, {90.08, -90.72}, {77.53, -96.02}, {69.07, -106.52}, {60.71,
        -121.62}, {56.94, -135.18}, {63.36, -149.76}},
  fillPattern= FillPattern.Solid,
  fillColor= {192,192,192},
  lineColor= {0,0,0},
  rotation= 0,
  smooth=Smooth.Bezier
),
Polygon(
  origin= {-100,100},
  lineThickness= 0.53,
  pattern= LinePattern.Solid,
  points= {{55.91, -142.11}, {52.04, -135.48}, {48.37, -122.43}, {46.94, -107.03}, {48.37,
        -93.26}, {51.53, -83.37}, {56.63, -78.07}, {63.76, -77.35}, {68.35, -78.48}, {68.46,
        -82.56}, {69.27, -86.63}, {77.33, -87.76}, {84.67, -89.49}, {83.04, -93.67}, {76.21,
        -102.75}, {67.64, -115.70}, {61.42, -127.32}, {58.87, -135.38}, {58.05, -140.89},
        {55.91, -142.11}},
  fillPattern= FillPattern.Solid,
  fillColor= {219,219,219},
  lineColor= {0,0,0},
  rotation= 0,
  smooth=Smooth.Bezier
),
Polygon(
  origin= {-100,100},
  lineThickness= 0.53,
  pattern= LinePattern.Solid,
  points= {{117.61, -84.91}, {115.67, -87.76}, {114.04, -92.09}, {115.85, -94.90}, {120.77,
        -97.35}, {126.28, -101.94}, {133.42, -101.83}, {138.25, -102.31}, {138.35, -97.63},
        {135.76, -94.19}, {133.58, -87.53}, {129.38, -82.02}, {122.20, -82.25}, {117.61,
        -84.91}},
  fillPattern= FillPattern.Solid,
  fillColor= {191,191,191},
  lineColor= {0,0,0},
  rotation= 0,
  smooth=Smooth.Bezier
),
Polygon(
  origin= {-100,100},
  lineThickness= 0.53,
  pattern= LinePattern.Solid,
  points= {{106.70, -59.41}, {111.59, -64.72}, {116.08, -65.23}, {128.42, -63.90}, {144.23,
        -65.53}, {155.81, -67.21}, {158.98, -68.69}, {160.48, -72.87}, {160.38, -79.81},
        {158.10, -84.09}, {149.94, -84.09}, {135.56, -81.34}, {123.32, -81.44}, {116.80,
        -86.33}, {114.97, -93.31}, {110.58, -91.53}, {102.42, -90.72}, {93.34, -91.64},
        {92.01, -89.19}, {94.77, -82.15}, {96.50, -75.93}, {95.48, -68.49}, {98.44, -59.82},
        {106.70, -59.41}},
  fillPattern= FillPattern.Solid,
  fillColor= {231,231,231},
  lineColor= {0,0,0},
  rotation= 0,
  smooth=Smooth.Bezier
),
Polygon(
  origin= {-100,100},
  lineThickness= 0.53,
  pattern= LinePattern.Solid,
  points= {{92.72, -84.60}, {95.48, -71.86}, {98.33, -62.68}, {103.33, -58.80}, {108.84,
        -58.50}, {112.30, -61.45}, {113.53, -65.43}, {119.54, -65.12}, {131.27, -64.51},
        {134.77, -64.90}, {137.26, -65.02}, {136.54, -61.49}, {135.86, -56.66}, {133.01,
        -46.46}, {129.33, -39.32}, {129.23, -35.86}, {134.23, -31.17}, {140.35, -26.17},
        {141.88, -22.80}, {139.74, -20.36}, {137.59, -19.34}, {131.99, -23.01}, {125.36,
        -29.43}, {125.66, -29.03}, {129.13, -24.13}, {131.37, -20.87}, {132.09, -18.42},
        {130.56, -16.18}, {128.01, -14.75}, {125.66, -15.16}, {121.07, -19.85}, {114.96,
        -25.56}, {109.65, -27.60}, {104.45, -26.99}, {100.68, -25.15}, {98.44, -21.58},
        {96.60, -15.87}, {94.87, -10.67}, {93.95, -8.12}, {93.03, -7.92}, {90.58, -8.99},
        {87.12, -10.72}, {85.89, -14.54}, {86.91, -20.15}, {87.47, -24.84}, {87.57, -28.92},
        {84.26, -36.78}, {78.14, -48.50}, {73.55, -59.82}, {70.39, -72.06}, {68.35, -81.85},
```

```
      {67.43, -86.34}, {75.08, -88.17}, {83.55, -89.29}, {82.57, -93.21}, {85.06, -94.36},
      {89.26, -92.25}, {92.72, -84.60}},
  fillPattern= FillPattern.Solid,
  fillColor= {176,176,176},
  lineColor= {0,0,0},
  rotation= 0,
  smooth=Smooth.Bezier
),
Polygon(
  origin= {-100,100},
  lineThickness= 0.53,
  pattern= LinePattern.Solid,
  points= {{135.53, -93.01}, {139.82, -92.83}, {144.10, -92.60}, {144.39, -91.39}, {144.39,
      -88.38}, {144.10, -85.78}, {143.63, -84.79}, {142.36, -84.33}, {138.83, -84.33},
      {136.57, -84.04}, {137.21, -83.58}, {140.45, -83}, {140.10, -82.01}, {133.74,
      -81.15}, {125.68, -81.06}, {121.31, -81.82}, {124.45, -82.51}, {128.65, -83.26},
      {130.96, -84.79}, {132.58, -87.34}, {133.62, -89.83}, {134.55, -91.97}, {135.53,
      -93.01}},
  fillPattern= FillPattern.Solid,
  fillColor= {161,161,161},
  lineColor= {0,0,0},
  rotation= 0,
  smooth=Smooth.Bezier
),
Polygon(
  origin= {-100,100},
  lineThickness= 0.53,
  pattern= LinePattern.Solid,
  points= {{55.58, -75.66}, {56.49, -72.12}, {57.65, -65.18}, {59.04, -53.43}, {60.14,
      -43.07}, {60.83, -36.25}, {61.76, -29.94}, {62.86, -24.04}, {63.96, -17.85}, {64.71,
      -13.97}, {66.27, -12.87}, {68.99, -12.35}, {71.89, -12.64}, {74.32, -13.62}, {75.30,
      -14.43}, {75.65, -19.76}, {76.05, -26.93}, {76.63, -29.48}, {79.23, -28.49}, {83.63,
      -25.54}, {87.10, -23.57}, {89.75, -22.53}, {91.93, -23.74}, {93.64, -26.82}, {94.22,
      -29.71}, {93.53, -32.02}, {87.16, -36.65}, {80.22, -42.38}, {78.66, -46.20}, {76.40,
      -53.26}, {72.93, -63.27}, {70.56, -72.47}, {69.28, -78.60}, {62.11, -79.41}, {55.20,
      -79.63}},
  fillPattern= FillPattern.Solid,
  fillColor= {225,225,225},
  lineColor= {0,0,0},
  rotation= 0,
  smooth=Smooth.Bezier
),
Ellipse(
  origin= {-100,100},
  lineThickness= 0.53,
  extent= {{45.36,-82.07},{55.94,-92.65}},
  pattern= LinePattern.Solid,
  fillPattern= FillPattern.Solid,
  fillColor= {17,89,255},
  lineColor= {17,89,255},
  rotation= 0
),
Ellipse(
  origin= {-100,100},
  lineThickness= 0.53,
  extent= {{69.17,-98.32},{79.75,-108.91}},
  pattern= LinePattern.Solid,
  fillPattern= FillPattern.Solid,
  fillColor= {17,89,255},
  lineColor= {17,89,255},
  rotation= 0
),
Line(
  origin= {-100,100},
  color= {17,89,255},
  pattern= LinePattern.Solid,
  thickness= 1,
  points= {{50.65, -87.74}, {53.67, -99.08}, {59.72, -105.88}, {68.04, -107.02}, {74.84,
      -105.13}},
```

```
    rotation= 0,
    smooth=Smooth.Bezier
  ),
  Line(
    origin= {-100,100},
    color= {17,89,255},
    pattern= LinePattern.Solid,
    thickness= 1,
    points= {{52.16, -84.72}, {63.88, -85.85}, {73.33, -91.14}, {74.84, -100.59}},
    rotation= 0,
    smooth=Smooth.Bezier
  ),
  Line(
    origin= {-100,100},
    color= {17,89,255},
    pattern= LinePattern.Solid,
    thickness= 1,
    points= {{53.29, -88.87}, {65.39, -94.54}, {74.08, -102.48}},
    rotation= 0,
    smooth=Smooth.Bezier
  ),
  Line(
    origin= {-100,100},
    color= {17,89,255},
    pattern= LinePattern.Solid,
    thickness= 1,
    points= {{74.84, -103.99}, {77.49, -123.65}, {68.41, -140.28}, {68.04, -146.33}},
    rotation= 0,
    smooth=Smooth.Bezier
  ),
  Line(
    origin= {-100,100},
    color= {17,89,255},
    pattern= LinePattern.Solid,
    thickness= 1,
    points= {{76.35, -102.10}, {95.25, -117.22}, {110.37, -117.98}, {112.75, -115.78},
         {120.20, -108.91}},
    rotation= 0,
    smooth=Smooth.Bezier
  ),
  Line(
    origin= {-100,100},
    color= {17,89,255},
    pattern= LinePattern.Solid,
    thickness= 1,
    points= {{74.84, -104.37}, {86.93, -124.78}, {90.71, -162.96}},
    rotation= 0,
    smooth=Smooth.Bezier
  ),
  Line(
    origin= {-100,100},
    color= {17,89,255},
    pattern= LinePattern.Solid,
    thickness= 1,
    points= {{76.35, -101.73}, {97.14, -109.29}, {102.43, -105.13}},
    rotation= 0,
    smooth=Smooth.Bezier
  ),
  Line(
    origin= {-100,100},
    color= {17,89,255},
    pattern= LinePattern.Solid,
    thickness= 1,
    points= {{75.60, -103.99}, {85.23, -113.44}, {94.87, -122.89}, {104.51, -132.34}, {112.64,
          -141.04}, {116.04, -143.21}, {117.55, -140.47}, {120.57, -135.08}, {130.02,
         -128.94}},
    rotation= 0,
    smooth=Smooth.Bezier
  ),
```

```
Line(
  origin= {-100,100},
  color= {17,89,255},
  pattern= LinePattern.Solid,
  thickness= 1,
  points= {{104.51, -132.34}, {108.57, -136.69}, {111.88, -154.26}, {109.61, -165.23}},
  rotation= 0,
  smooth=Smooth.Bezier
),
Line(
  origin= {-100,100},
  color= {17,89,255},
  pattern= LinePattern.Solid,
  thickness= 1,
  points= {{105.44, -117.73}, {110.37, -117.98}, {116.41, -119.12}, {121.71, -119.87}, {127,
       -120.62}, {133.05, -116.85}},
  rotation= 0,
  smooth=Smooth.Bezier
),
Line(
  origin= {-100,100},
  color= {17,89,255},
  pattern= LinePattern.Solid,
  thickness= 1,
  points= {{88.56, -141.19}, {88.82, -143.87}, {81.26, -154.64}},
  rotation= 0,
  smooth=Smooth.Bezier
),
Line(
  origin= {-100,100},
  color= {17,89,255},
  pattern= LinePattern.Solid,
  thickness= 1,
  points= {{117.55, -140.47}, {119.06, -137.78}, {136.45, -145.19}, {147.03, -141.79}},
  rotation= 0,
  smooth=Smooth.Bezier
),
Line(
  origin= {-100,100},
  color= {17,89,255},
  pattern= LinePattern.Solid,
  thickness= 1,
  points= {{112.64, -141.04}, {116.04, -143.21}, {122.28, -150.67}, {126.34, -155.30},
      {130.40, -159.93}, {128.13, -173.92}},
  rotation= 0,
  smooth=Smooth.Bezier
),
Line(
  origin= {-100,100},
  color= {17,89,255},
  pattern= LinePattern.Solid,
  thickness= 1,
  points= {{124.31, -152.99}, {126.34, -155.30}, {137.21, -156.15}, {143.63, -152.75}},
  rotation= 0,
  smooth=Smooth.Bezier
),
Ellipse(
  origin= {-100,100},
  lineThickness= 0.53,
  pattern= LinePattern.None,
  fillPattern= FillPattern.Solid,
  fillColor= {107,107,107},
  extent= {{152.88,-69.07},{159.03,-82.90}},
  rotation= 0
),
Ellipse(
  origin= {-100,100},
  lineThickness= 0.53,
  pattern= LinePattern.None,
```

```
          fillPattern= FillPattern.Solid,
          fillColor= {107,107,107},
          extent= {{72.89,-54.28},{76.36,-62.10}},
          rotation= 20.90
        )
      }
    )
  );
end Heart;
```

Supplementary Listing 26: `SHMConduction/Icons/Hourglass.mo`

```
within SHMConduction.Icons;
model Hourglass "hourglass icon for delay"
  annotation(
    Icon(
      coordinateSystem(
        preserveAspectRatio= false,
        extent= {{-100,-100},{100,100}}
      ),
      graphics= {
        Polygon(
          origin= {-100,100},
          lineThickness= 2,
          pattern= LinePattern.Solid,
          points= {{60.25, -28.44}, {139.75, -28.44}, {60.25, -171.56}, {139.75, -171.56}},
          fillPattern= FillPattern.None,
          lineColor= {0,0,0},
          rotation= 0
        ),
        Polygon(
          origin= {-100,100},
          lineThickness= 1,
          pattern= LinePattern.None,
          points= {{78.80, -61.84}, {121.20, -61.84}, {100, -100}},
          fillPattern= FillPattern.Solid,
          fillColor= {0,0,0},
          rotation= 0
        ),
        Polygon(
          origin= {-100,100},
          lineThickness= 1,
          pattern= LinePattern.None,
          points= {{60.25, -171.56}, {100, -155.65}, {139.75, -171.56}},
          fillPattern= FillPattern.Solid,
          fillColor= {0,0,0},
          rotation= 0
        ),
        Ellipse(
          origin= {-100,100},
          lineThickness= 1,
          pattern= LinePattern.None,
          fillPattern= FillPattern.Solid,
          fillColor= {0,0,0},
          extent= {{98.41,-112.72},{101.59,-115.90}},
          rotation= 0
        ),
        Ellipse(
          origin= {-100,100},
          lineThickness= 1,
          pattern= LinePattern.None,
          fillPattern= FillPattern.Solid,
          fillColor= {0,0,0},
          extent= {{98.41,-123.85},{101.59,-127.03}},
          rotation= 0
        ),
        Ellipse(
          origin= {-100,100},
```

```
    lineThickness= 1,
    pattern= LinePattern.None,
    fillPattern= FillPattern.Solid,
    fillColor= {0,0,0},
    extent= {{98.41,-134.98},{101.59,-138.16}},
    rotation= 0
  ),
  Ellipse(
    origin= {-100,100},
    lineThickness= 1,
    pattern= LinePattern.None,
    fillPattern= FillPattern.Solid,
    fillColor= {0,0,0},
    extent= {{98.41,-146.11},{101.59,-149.29}},
    rotation= 0
  ),
  Ellipse(
    origin= {-100,100},
    lineThickness= 2,
    extent= {{2.05,-2.05},{197.95,-197.95}},
    pattern= LinePattern.Solid,
    fillPattern= FillPattern.None,
    lineColor= {0,0,0},
    rotation= 0
  ),
  Ellipse(
    origin= {-107,112.51},
    lineThickness= 2,
    extent= {{130.21,-88.38},{178.47,-136.64}},
    pattern= LinePattern.Solid,
    fillPattern= FillPattern.None,
    lineColor= {0,0,0},
    rotation= -0
  ),
  Line(
    origin= {-107,112.51},
    color= {0,0,0},
    pattern= LinePattern.Solid,
    thickness= 2,
    points= {{153.25, -104.27}, {153.25, -116.19}, {168.83, -116.19}},
    rotation= -0
  ),
  Line(
    origin= {-107,112.51},
    color= {0,0,0},
    pattern= LinePattern.Solid,
    thickness= 1,
    points= {{154.34, -130.60}, {154.34, -136.64}},
    rotation= -0
  ),
  Line(
    origin= {-107,112.51},
    color= {0,0,0},
    pattern= LinePattern.Solid,
    thickness= 1,
    points= {{154.34, -88.38}, {154.34, -94.42}},
    rotation= -0
  ),
  Line(
    origin= {-107,112.51},
    color= {0,0,0},
    pattern= LinePattern.Solid,
    thickness= 1,
    points= {{130.21, -112.51}, {136.25, -112.51}},
    rotation= -0
  ),
  Line(
    origin= {-107,112.51},
    color= {0,0,0},
```

```
  pattern= LinePattern.Solid,
  thickness= 1,
  points= {{172.42, -112.51}, {178.47, -112.51}},
  rotation= -0
),
Line(
  origin= {-107,112.51},
  color= {0,0,0},
  pattern= LinePattern.Solid,
  thickness= 1,
  points= {{166.40, -91.61}, {164.39, -95.09}},
  rotation= -0
),
Line(
  origin= {-107,112.51},
  color= {0,0,0},
  pattern= LinePattern.Solid,
  thickness= 1,
  points= {{175.23, -100.44}, {171.76, -102.45}},
  rotation= -0
),
Line(
  origin= {-107,112.51},
  color= {0,0,0},
  pattern= LinePattern.Solid,
  thickness= 1,
  points= {{175.23, -124.57}, {171.76, -122.57}},
  rotation= -0
),
Line(
  origin= {-107,112.51},
  color= {0,0,0},
  pattern= LinePattern.Solid,
  thickness= 1,
  points= {{166.40, -133.41}, {164.39, -129.93}},
  rotation= -0
),
Line(
  origin= {-107,112.51},
  color= {0,0,0},
  pattern= LinePattern.Solid,
  thickness= 1,
  points= {{142.27, -133.41}, {144.28, -129.93}},
  rotation= -0
),
Line(
  origin= {-107,112.51},
  color= {0,0,0},
  pattern= LinePattern.Solid,
  thickness= 1,
  points= {{133.44, -124.57}, {136.92, -122.57}},
  rotation= -0
),
Line(
  origin= {-107,112.51},
  color= {0,0,0},
  pattern= LinePattern.Solid,
  thickness= 1,
  points= {{133.44, -100.44}, {136.92, -102.45}},
  rotation= -0
),
Line(
  origin= {-107,112.51},
  color= {0,0,0},
  pattern= LinePattern.Solid,
  thickness= 1,
  points= {{142.27, -91.61}, {144.28, -95.09}},
  rotation= -0
),
```

```
        Ellipse(
          origin= {-107,112.51},
          pattern= LinePattern.None,
          fillPattern= FillPattern.Solid,
          fillColor= {0,0,0},
          extent= {{153.49,-87.53},{155.18,-89.23}},
          rotation= -0
        ),
        Ellipse(
          origin= {-107,112.51},
          pattern= LinePattern.None,
          fillPattern= FillPattern.Solid,
          fillColor= {0,0,0},
          extent= {{153.49,-87.53},{155.18,-89.23}},
          rotation= -0
        ),
        Line(
          origin= {-107,112.51},
          color= {0,0,0},
          pattern= LinePattern.Solid,
          thickness= 2,
          points= {{154.34, -88.38}, {154.34, -82.29}},
          rotation= -0
        ),
        Rectangle(
          origin= {-107,112.51},
          lineThickness= 2.12,
          pattern= LinePattern.None,
          fillPattern= FillPattern.Solid,
          fillColor= {0,0,0},
          extent= {{148.50,-80.83},{160.18,-83.75}},
          rotation= -0
        )
      }
    )
  );
end Hourglass;
```

Supplementary Listing 27: SHMConduction/Icons/Metronome.mo

```
within SHMConduction.Icons;
model Metronome "metronome icon for pacemaker"
  annotation(
    Icon(
      coordinateSystem(
        preserveAspectRatio= false,
        extent= {{-100,-100},{100,100}}
      ),
      graphics= {
        Line(
          origin= {-100,100},
          pattern= LinePattern.Solid,
          rotation= 0,
          points= {{98.41, -140.61}, {160.53, -33.61}},
          thickness= 2
        ),
        Polygon(
          origin= {-100,100},
          lineThickness= 2,
          pattern= LinePattern.Solid,
          points= {{82.31, -34.98}, {115.02, -34.98}, {147.73, -152.74}, {49.60, -152.74}},
          fillPattern= FillPattern.None,
          rotation= 0
        ),
        Ellipse(
          origin= {-100,100},
          lineThickness= 2,
          pattern= LinePattern.Solid,
```

```
        endAngle= 179.999997341,
        fillPattern= FillPattern.Solid,
        fillColor= {255,255,255},
        extent= {{74.88,-117.33},{122.44,-164.88}},
        startAngle= 0.0,
        rotation= 0
      ),
      Polygon(
        origin= {-100,100},
        lineThickness= 2,
        pattern= LinePattern.None,
        points= {{143.95, -55.42}, {149.79, -58.93}, {159.31, -49.78}, {147.56, -42.71}},
        fillPattern= FillPattern.Solid,
        fillColor= {0,0,0},
        rotation= 0
      )
    }
  )
);
end Metronome;
```
